# Evolution of Evolvability In Rapidly Adapting Populations

**DOI:** 10.1101/2023.07.12.548717

**Authors:** James T. Ferrare, Benjamin H. Good

## Abstract

Mutations can alter the short-term fitness of an organism, as well as the rates and benefits of future mutations. While numerous examples of these evolvability modifiers have been observed in rapidly adapting microbial populations, existing theory struggles to predict when they will be favored by natural selection. Here, we develop a mathematical framework for predicting the fates of genetic variants that modify the rates and benefits of future mutations in linked genomic regions. We derive analytical expressions showing how the fixation probabilities of these variants depend on the size of the population and the diversity of competing mutations. We find that competition between linked mutations can dramatically enhance selection for modifiers that increase the benefits of future mutations, even when they impose a strong direct cost on fitness. However, we also find that modest direct benefits can be sufficient to drive evolutionary dead-ends to fixation. Our results suggest that subtle differences in evolvability could play an important role in shaping the long-term success of genetic variants in rapidly evolving microbial populations.

## Main Text

The benefits of new mutations can manifest over multiple timescales. Some mutations alter the short-term fitness of an organism, while others can also affect the rates and fitness benefits of future mutations. Examples are common in the microbial world. Mutations in DNA repair genes can generate mutator strains with dramatically elevated mutation rates (1–3). These same variants also alter the molecular spectrum of new mutations, which can shift the relative probabilities of adaptive variants in addition to their overall rates (4–6). Other classes of mutations can open or close novel adaptive pathways through epistatic interactions with other genes (7–12). Striking examples of these “evolvability modifiers” have been observed in laboratory evolution experiments (8, 10, 11, 13; Fig. 1A), and they are thought to play a critical role in cancer (14–16) and the evolution of antibiotic resistance (17–19). But despite their potential importance, it is difficult predict when these long-term benefits should be favored by natural selection.

**Fig 1.**
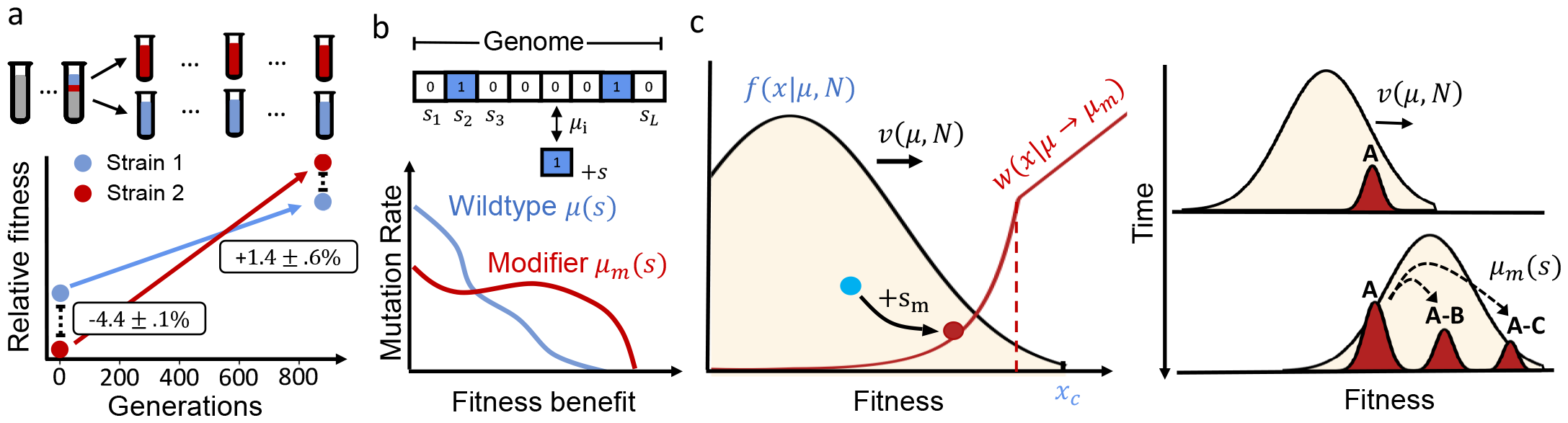
Modeling selection on evolvability in rapidly adapting asexual populations. **(a)** An empirical example of an evolvability modifier from Ref. (8). Two *E. coli* strains isolated from a long-term evolution experiment (42) had an initial fitness difference of ∼4%. The less-fit strain exhibited a higher rate of adaptation in replay experiments, which allowed it to consistently overcome its initial disadvantage after ∼900 generations of evolution. Fitness differences denote mean ± s.e.m. from 20 independent replays. **(b)** These long-term benefits can be modeled by the accumulation of beneficial mutations at a large number of linked genomic loci. The fitness benefits of the mutations are summarized by their distribution *μ s ds*, which denotes the total rate of producing mutations with fitness effects *s* ± *ds*/2. An evolvability modifier changes this distribution to a new value, *μ*_*m*_(*s*). **(c)** A modifier with a direct cost or benefit (*s*_*m*_) arises on a genetic background from the wildtype fitness distribution, *f*(*x*|*μ, N*), which has a maximum relativ fitness *x*_*c*_ (left). The modifier lineage competes with the wildtype population as they both acquire further mutations (right). The outcome of this competition is described by the conditional fixation probability *w*(*x* + *s*_*m*_| *μ* → *μ*_*m*_), which exhibits a sharp transition at a critical initial fitness *x*_*cm*_ ≠ *x*_*c*_.

Classical arguments from modifier theory suggest that asexual populations will select for mutations that maximize their long-term mean fitness (20, 21; SI Section 1). This simple result applies for infinite populations near mutation-selection equilibrium. Both conditions are frequently violated in adapting populations, since a beneficial variant can fix before its long-term costs or benefits are fully realized. This greediness creates an inherent tension between short-term and long-term fitness (22, 23).

While some of these trade-offs can be understood in simple cases where mutations accumulate one-by-one (24–27; SI Section 2), most microbial populations reside in a qualitatively different regime. In many cases of interest, from laboratory evolution experiments (10, 28–30) to natural populations of viruses (31–34), bacteria (35–37), and certain cancers (14, 15), multiple beneficial mutations can arise and compete within the population at the same time. The competition between these linked mutations (“clonal interference”; 38) ensures that a successful variant must often generate multiple additional mutations to fix, which amplifies the indirect selection for evolvability. While recent work has started to explore these effects for mutation-rate modifiers alone (24), we currently lack an analytical framework for understanding more general differences in evolvability in the high-diversity regimes most relevant for microbes.

Our limited understanding of these dynamics leaves many basic questions unresolved: How does natural selection balance the short-term costs or benefits of a mutation with its long-term impact on evolvability? Which evolvability differences matter most for determining a mutation’s long-term fate? And how do these answers depend on external factors like the size of the population or the diversity of competing mutations?

### Modeling second-order selection in rapidly adapting populations

To address these questions, we analyze the fates of general evolvability modifiers in a population genetic model that explicitly accounts for interference among competing beneficial mutations. We consider an asexual population of *N* individuals that can acquire beneficial mutations at a large number of linked genetic loci. The mutations accessible to each genotype 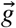 can be summarized by their distribution of fitness effects, 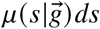, which represents the per generation rate of producing mutations with fitness effects *s* ± *ds*/2 (Fig. 1B). For most of this work, we will assume that the beneficial sites are sufficiently numerous and epistasis sufficiently weak that *μ*(*s*) remains approximately constant over the relevant timescales (which we determine self-consistently below). This implies that the rate of adaptation of the population will approach a steady-state value *v* (*μ, N)* that depends on the values of *μ(s*) and *N* (39, 40). Given these assumptions, the simplest possible evolvability modifier is one that changes the distribution of fitness effects to a new value, *μ*_*m*_*(s*), which is maintained for several additional substitutions (Fig. 1B). This can be viewed as the leading-order term in a more general expansion in the amount of “macroscopic epistasis” (41) in 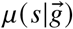 (Fig. S1; SI Section 3.1). It generalizes the notion of a mutator allele to capture more subtle changes in evolvability, while bypassing the tremendous complexity of the underlying fitness landscape.

If the modifier takes over the population, the rate of adaptation will shift to a new value *v* (*μ*_*m*_, *N*) that reflects the altered supply of mutations (e.g. Fig. 1A). To reach these high frequencies, the modifier must initially grow from a single founding individual, where stochastic fluctuations play a critical role. The outcome of this competition can be described by the fixation probability, *p*_fix_ (*μ* → *μ*_*m*_), which provides a quantitative measure of the mutant’s long-term reproductive value.

The fate of the modifier will strongly depend on its initial genetic background. While the distribution of backgrounds is often complicated at the genetic level, previous work has shown that progress can be made by grouping individuals by their relative fitness and modeling the resulting dynamics in fitness space (39, 40, 43). The distribution of fitnesses in the background population will approach a steady-state shape *f*(*x* |*μ, N*) that increases in fitness at rate *v*(*μ, N*) (Fig. 1C). This distribution has a characteristic width, *x*_*c*_, which roughly coincides with the fittest individuals in the population. A new modifier mutation will arise on a genetic background with fitness *x*, and will compete with the wildtype population while acquiring further mutations from *μ*_*m*_(*s*) (Fig. 1C, right). The outcome of this competition can be summarized by the conditional fixation probability *w*(*x*|*μ* →*μ*_*m*_) (Fig. 1C, left). In rapidly adapting populations, the fate of the modifier will often be determined while it is still at a low frequency in the population (SI Section 3). When this assumption holds, the conditional fixation probability can be approximated by a standard branching process recursion,

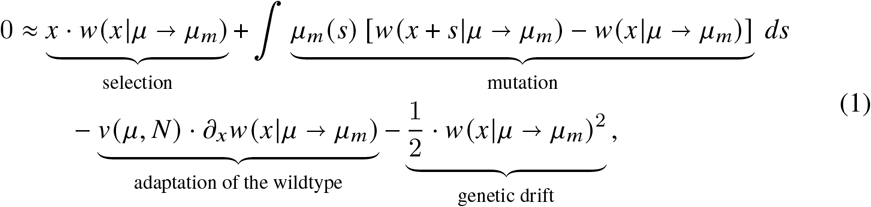

which represents a balance between (i) the growth of the lineage due to selection, (ii) the adaptation of the background population, (iii) the stochastic effects of genetic drift when the lineage is at low copy-number, and (iv) the production of further mutations (SI Section 3). The overall fixation probability of the modifier can then be obtained by averaging over its random genetic background,

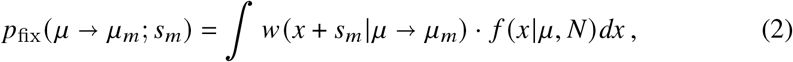

where we have also allowed the modifier to have a direct cost or benefit *s*_*m*_. Together, Eqs. (1) and (2) provide a quantitative framework for understanding the trade-offs between “first-order” selection on fitness and “second-order” selection on evolvability that generalizes the mutator model in Ref. (24).

The fixation probability of a neutral mutation must always satisfy *p*_fix_ (*μ* → *μ*; 0) = 1/*N* (44). This provides a natural scale for interpreting the fixation probability in Eq. (2). Variants with *p*_fix_(*μ* → *μ*_*m*_; *s*_*m*_) ≫ 1/*N* are strongly favored by natural selection, while variants with *p*_fix_(*μ* → *μ*_*m*_; *s*_*m*_) ≪ 1/*N* are effectively purged. For this reason, it will be convenient to examine the scaled fixation probability, 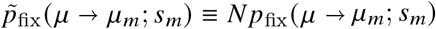, so that the sign of 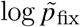coincides with the net “direction” of natural selection (45).

### Selection for evolvability in a simple fitness landscape

To understand the second-order selection pressures encoded in Eqs. (1) and (2), we start by considering a simple model for the distribution of fitness effects, where all mutations confer the same characteristic fitness benefit *s*_*b*_ and occur at a total rate *U*_*b*_. The most general evolvability modifier will either change the selection strength 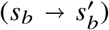, the mutation rate 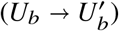, or both (Fig. 2A). This simplified model allows us to obtain an analytical solution for the fixation probability that is valid for empirically relevant scenarios where 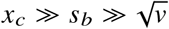 (SI Section 4).

**Fig 2.**
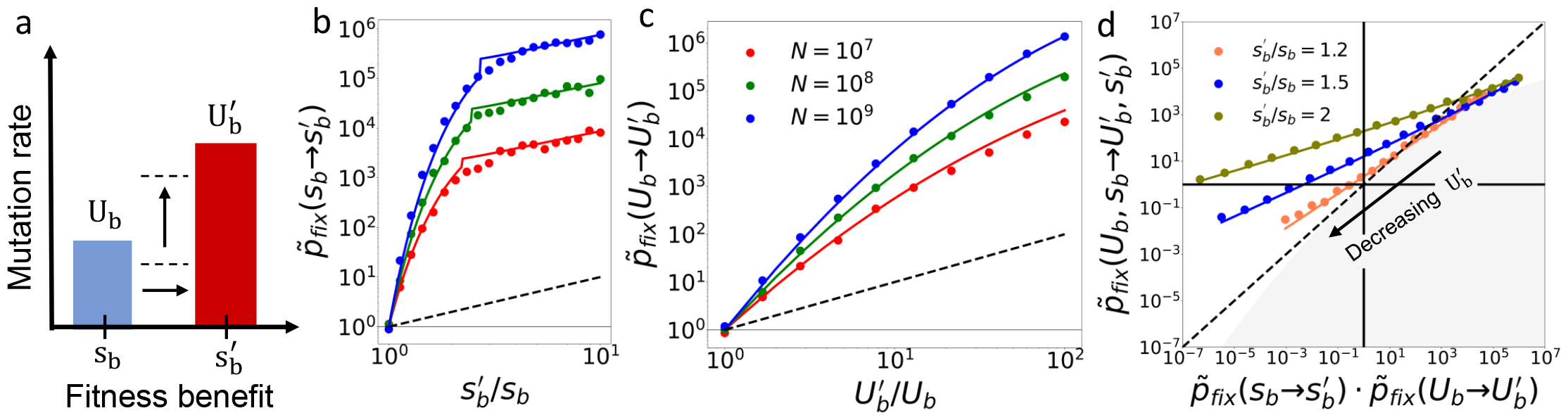
Interference between linked mutations enhances second-order selection for evolvability. **(a)** A simplified model for the distribution of fitness effects (Fig. 1B), where all mutations confer the same characteristic fitness benefit. An evolvability modifier will change the overall mutation rate 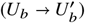, the overall selection strength 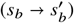, or both. **(b, c)** The fixation probability (scaled by the neutral expectation, 1/*N*) for a selection-strength modifier (b) or a mutation-rate modifier (c). Symbols denote the results of forward-time simulations (SI Section 8.1) for *s*_*b*_ = 10^−2^, *U*_*b*_ = 10^−5^, and *N* = 10^7^ − 10^9^. Solid lines denote our theoretical predictions (SI Section 8.2), while the dashed lines denote the null expectation in the absence of clonal interference (SI Section 2). **(d)** Fixation probability of a compound modifier (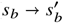 and 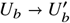) compared to an additive null model. Symbols denote the results of forward-time simulations for *s*_*b*_ = 10^−2^, *U*_*b*_ = 10^−5^, and *N* = 10^8^; the y-coordinates are obtained from simulations, while the x-coordinates are obtained from the theoretical predictions in panels b and c. Solid lines denote our theoretical predictions, which deviate substantially from the additive expectation (dashed lines). The gray region indicates forbidden combinations that cannot arise in our theory (SI Section 8.2).

For a pure selection-strength modifier 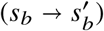, we find that the fixation probability initially grows as an exponential function of the new selection strength,

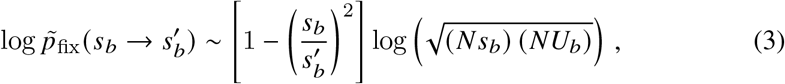

before saturating to a linear scaling when 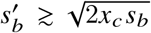 (Fig. 2B). This behavior is qualitatively different than that observed for mutation-rate modifiers (24),

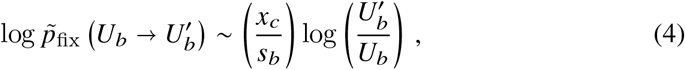

whose benefits increase more slowly with the mutation rate (Fig. 2C). In both cases, the fixation probabilities in Fig. 2B,C are significantly larger than the linear scaling expected when mutations accumulate via discrete selective sweeps (Fig. 2B,C; SI Section 2). This gap grows increasingly large as a function of *NU*_*b*_, demonstrating that the competition between linked mutations enhances second-order selection for evolvability.

The origin of this behavior has a simple heuristic explanation. For small changes in the selection coefficient 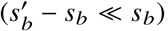, successful mutations typically arise in the high-fitness “nose” of the population (*x* ≈ *x*_*c*_ ≫ *s*_*b*_; Fig. 1C) and must acquire ∼*x*_*c*_/*s*_*b*_ additional mutations before they reach appreciable frequencies. In each of these steps (*j* = 1 … *x*_*c*_/*s*_*b*_), a selection-strength modifier produces 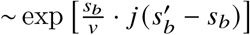 more mutations than a wildtype individual with the same fitness, leading to the exponential scaling observed in Eq. (3). The linear saturation at larger values of 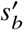 occurs when a single additional mutation is sufficient to ensure fixation. However, since successful modifiers still arise on anomalously fit genetic backgrounds 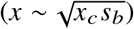, they are able to hitchhike to higher frequencies than they would on their own, thereby increasing their overall probability of producing a successful mutation. Both effects become increasingly important for larger values of *x*_*c*_/*s*_*b*_, which highlights how they critically depend on the genetic diversity that is maintained within the population.

The strength of second-order selection can be intuitively quantified by comparing Eqs. (3) and (4) with the corresponding fixation probability of a first-order mutation,

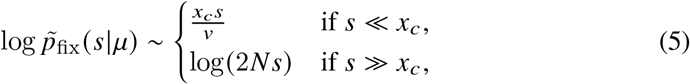

that has been studied in previous work (39, 40, 46; SI Section 4.1.3). Comparing this expression with Eq. (3) shows that even fractional changes in the selection coefficient 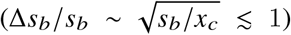 can generate fixation probabilities as large as a typical beneficial mutation (*s* ≈ *s*_*b*_). This contrasts with the behavior observed for a mutation-rate modifier in Eq. (4), where the mutation rate must increase by several orders of magnitude 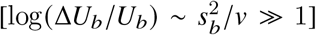 to achieve the same effect. This shows that large populations can more efficiently select on changes to the fitness benefits of future mutations, compared to the overall rate at which they occur.

The fixation probability of a compound modifier 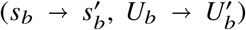 can be understood using these basic building blocks. We find that the leading-order contributions can be expressed as a linear combination,

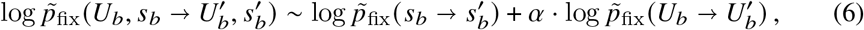

where the weighting factor *α* is given by 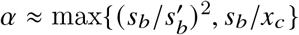. This additional weighting factor implies that mutation-rate and selection-strength modifiers do not additively combine: an increase in *s*_*b*_ will act to temper selection on the mutation rate, while decreases in *s*_*b*_ will amplify it. This non-additivity arises because larger selection-strength modifiers eventually lower the number of mutations required to fix 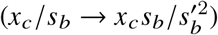, which diminishes the compounding effects of the altered mutation rate. These differences can be large, and can alter the sign of selection on the modifier (Fig. 2D). Moreover, since the individual components in Eq. (6) depend on the underlying parameters in different ways, the sign of selection can also vary as a function of the population size and the overall mutation rate (24, 26). These examples illustrate how selection-strength modifiers can lead to qualitatively different behavior than expected for mutation-rate changes alone, and that even modest shifts in *s*_*b*_ can frequently overpower order-of-magnitude differences in *U*_*b*_.

### Trade-offs between first- and second-order selection

We are now in a position to understand how natural selection balances the short-term costs and benefits of a mutation with its long-term impact on evolvability. When the evolvability deficits are not too large (SI Section 4.1.5), we find that the leading-order behavior can be decomposed into contributions from first- and second-order selection,

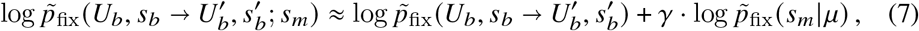

where 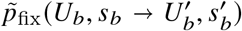 is the fixation probability of the modifier from Eq. (6), 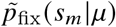 is the fixation probability of a first-order mutation from Eq. (5), and *γ* is an additional weighting factor satisfying 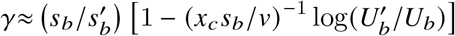. This decomposition shows that differences in evolvability will generally modulate the effects of first-order selection on fitness.

A modifier that is strongly beneficial in the absence of a direct cost or benefit 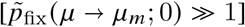 will always reduce the relative contribution from *s*_*m*_ (*γ* ≲ 1). As a result, strongly beneficial modifiers can remain positively selected even when they impose a large direct cost on fitness (|*s*_*m*_| ≫ *s*_*b*_; Fig. 3A,C). This contrasts with the classical behavior observed for discrete selective sweeps, where direct costs larger than *s*_*b*_ will typically prevent a modifier from fixing (SI Section 2). Larger populations are therefore better able to sacrifice short-term fitness for long-term gains in evolvability.

**Fig 3.**
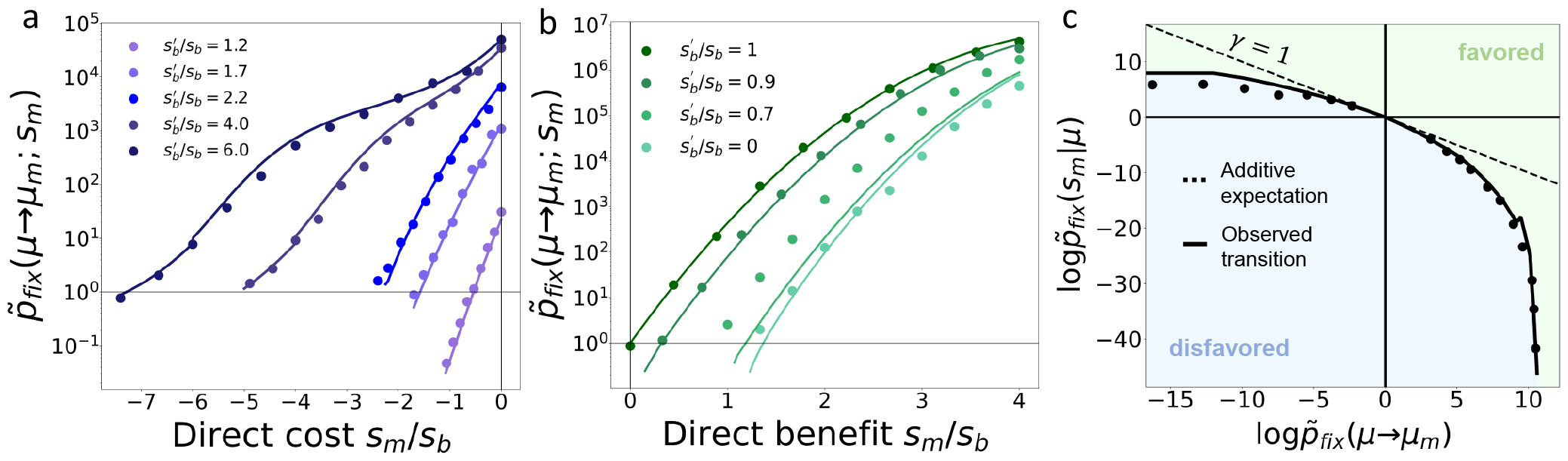
Trade-offs between first- and second-order selection. **(a, b)** Fixation probability of a selection-strength modifier with a direct cost (a) or benefit (b). Symbols denote the results of forward-time simulations for *s*_*b*_ = 10^−2^, 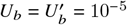, and *N* = 10^8^, while solid lines denote our theoretical predictions (SI Section 8.2). **(c)** Phase diagram illustrating the transition between favored (green) and unfavored modifiers (blue). Symbols denote results of forward time simulations for the selection-strength modifiers in panels a and b; x-coordinates show the measured fixation probabilities in the absence of a direct cost or benefit (y-intercepts in panels a and b), while the y-coordinates show the predicted fixation probability of a first-order mutation inferred from the x-intercept in panels a and b 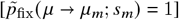. Solid lines show our theoretical predictions (SI Section 8.2), which exhibit large deviations from the additive expectation (*γ* = 1, dashed line).

The opposite behavior occurs when short-term fitness benefits are linked to long-term reductions in evolvability. In this case, strongly deleterious modifiers 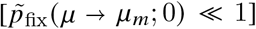 will generally amplify the relative contribution of a short-term fitness benefit (Fig. 3B,C). A striking example of this effect occurs in the extreme case of an evolutionary “dead end”, where further beneficial mutations are not available. These local fitness peaks cannot be captured by the linear approximation in Eq. (1), since their long-term success depends on their ability to influence the adaptation of the wildtype.

Remarkably, the strength of this feedback is still dominated by chance events that occur while the modifier lineage is rare. This allows us to generalize Eq. (1) to account for this non-linear behavior (SI Section 4.1.6). We find that moderate direct benefits as small as ≈0.4*x*_*c*_ are sufficient to cause an evolutionary dead-end to be positively selected 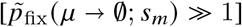, even though they cause the long-term rate of adaptation to drop to zero if they fix (Fig. 3B). This critical fitness benefit is often larger than the size of a single driver mutation (*s*_*b*_), but it is also smaller than the total fitness variation maintained within the population (*x*_*c*_) and is only weakly dependent on *NU*_*b*_. These examples illustrate how the evolutionary foresight of natural selection can be highly asymmetric: larger populations can greedily select for mutations that lower their rate of adaptation, even while they are better able to endure short-term fitness costs to realize long-term evolutionary benefits.

### Continuous distributions of fitness effects

We have so far focused on a simplified model of the fitness landscape, where all beneficial mutations confer the same characteristic fitness benefit. However, most organisms produce mutations with a range of different fitness benefits. A general evolvability modifier will therefore correspond to a continuous perturbation of the distribution of fitness effects, *δμ*(*s*) = *μ*_*m*_(*s*) − *μ*(*s*), representing the addition or subtraction of mutations with a range of fitness benefits (Fig. 4A). How does second-order selection on these more general evolvability differences relate to the selection-strength and mutation-rate and axes in Fig. 2?

**Fig 4.**
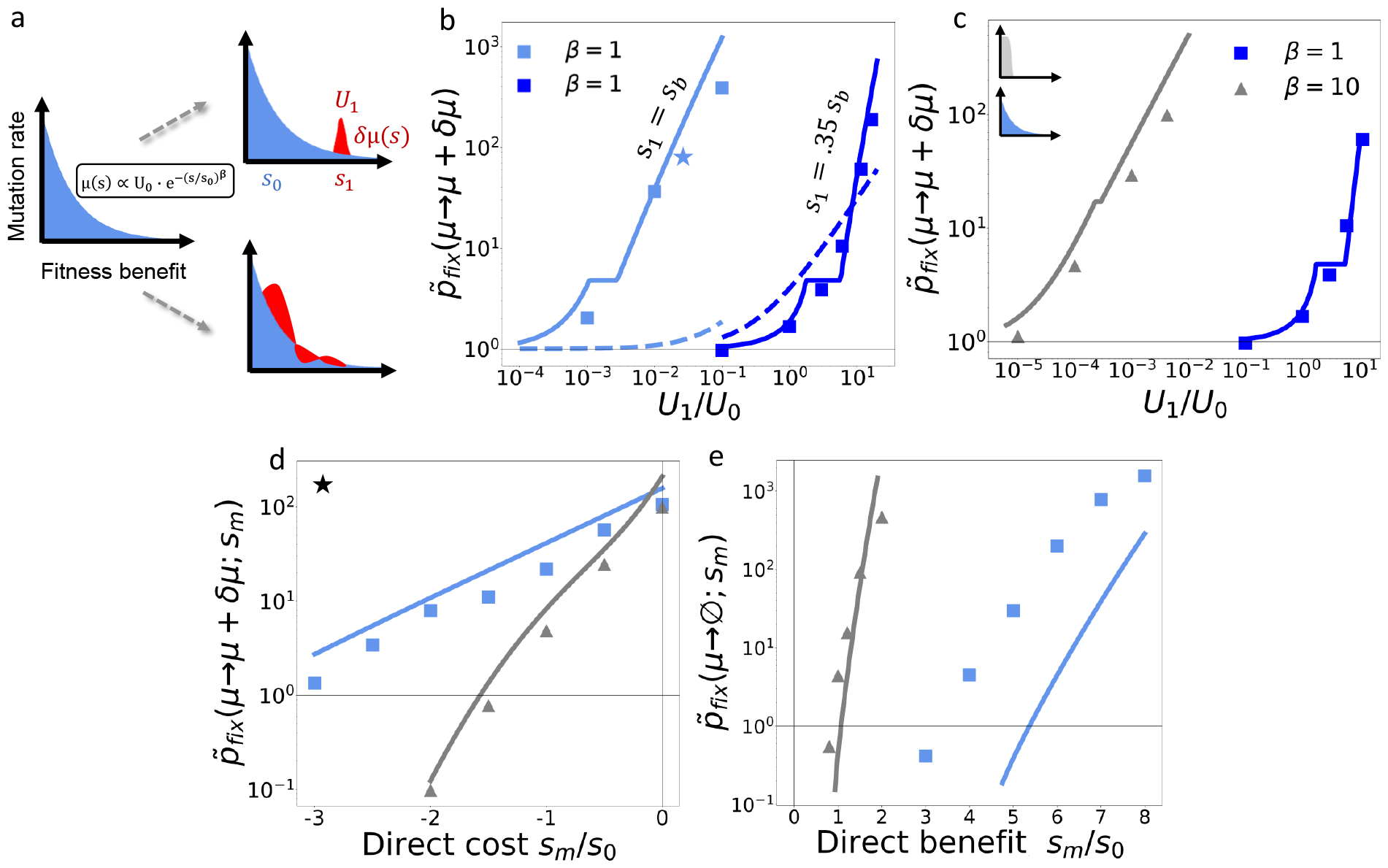
Second-order selection with a continuous distribution of fitness effects. **(a)** Examples of evolvability modifiers for a continuous distribution of fitness effects. Top right panel shows a small local addition *δμ* (*s*) ≈ *U*_1_*δ*(*s* − *s*_1_) (red), while bottom right panel shows a *δμ*(*s*) with a mixture of positive and negative changes. **(b, c)** Scaled fixation probability of the modifier in panel a (top right) for an exponential *μ* (*s*) (*β* = 1), as well as a distribution with a shorter tail (*β* = 10). Blue and gray curves in the inset of panel c illustrate differences in the shapes of these background distributions. In both panels, symbols denote the results of forward-time simulations for *s*_0_ = 10^−2^, *U*_0_ = 10^−5^, and *N* = 10^8^. Solid lines denote our theoretical predictions (SI Section 8.2), while dashed lines indicate the null expectation in the absence of clonal interference. **(d)** Fixation probability of a modifier with a direct cost. Base parameters are indicated by the stars in panel c. **(e)** Fixation probability of an evolutionary dead-end with a short-term fitness benefit. Symbols denote the results of forward-time simulations for *s*_0_ = 10^−2^, *U*_0_ = 10^−5^, and *N* = 10^8^, while solid lines denote our theoretical predictions.

We can extend our solution of Eq. (1) to a large class of wildtype mutation distributions that have been studied in previous work (SI Section 5). In these settings, the fitness benefits of fixed mutations are strongly peaked around a characteristic fitness benefit *s*_*b*_ (*μ, N*) [with a corresponding mutation rate *U*_*b*_ (*μ, N*)], even when *μ* (*s*) is more broadly distributed [e.g. an exponential distribution] (39, 40, 47, 48; SI Section 5.1). In terms of these parameters, we find that selection on a general evolvability modifier [*μ*(*s*) → *μ*(*s*) + *δμ*(*s*)] sensitively depends on the integral,

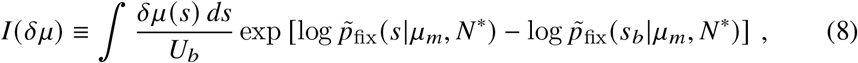

where *p*_fix_(*s*| *μ, N*) denotes the fixation probability of a first-order mutation from Eq. (5) and *N*^*^ is defined by *v* (*μ*_*m*_, *N*^*^) = *v* (SI Section 5.2). When *I* (*δμ*) ≲ 1, the fixation probability of the modifier initially grows as 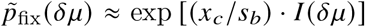. This shows that the relevant fitness scale for second-order selection is set by *s*_*b*_. For *s > s*_*b*_, even small increases in the net mutation rate 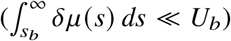 can generate large changes in the fixation probability (Fig. 4B). In contrast, fitness benefits slightly smaller than *s*_*b*_ require many multiples of *U*_*b*_ to have the same effect. Since *s*_*b*_ and *U*_*b*_ both emerge from the competition between linked mutations, the location of this transition can vary with the population size and the shape of *μ*(*s*) (Fig. 4C).

For more strongly beneficial modifiers 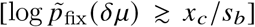, the integrand in Eq. (8), is often concentrated around a characteristic fitness benefit 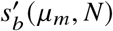, which depends on the perturbation *δμ* (*s*) as well as the wildtype parameters *s*_*b*_ and *U*_*b*_. In this case, we find that the fate of the modifier can be predicted from our single-*s* theory in Eq. (6),

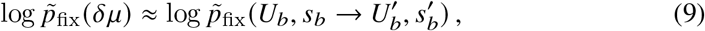

where 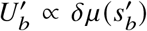 is the total mutation rate for mutations close to 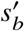 (Fig. 4B-E; SI Section 5). This effective selection coefficient coincides with the mutations most likely to fix in a successful modifier lineage (SI Section 5), providing a mechanistic interpretation for the effective parameters.

The equivalence principle in Eq. (9) shows that second-order selection can generally be understood as a combination of the mutation-rate and selection-strength axes in Fig. 2. However, the relevant parameters will not coincide with the mean and height of *μ*(*s*). Instead, due to the exponential weighting of mutations with *s* ≳ *s*_*b*_, otherwise subtle additions to *μ*(*s*) can be strongly favored by natural selection, even when they have a negligible impact on the overall mean or mutation rate (Fig. 4B-D). Conversely, large reductions in these global parameters will be invisible to natural selection unless they also deplete mutations near *s*_*b*_. This sensitivity to local changes could help explain previous experimental observations in *E. coli* (Fig. 1A), where the potentiation of just a few beneficial genes was sufficient to overcome a large short-term fitness cost (8).

## Discussion

Our results provide a framework for understanding how “first-order” selection for fitness interacts with “second-order” selection for evolvability in rapidly adapting populations. We have shown that when beneficial mutations are common, the competition between linked mutations can dramatically enhance selection for subtle differences in evolvability, leading to large deviations from the linear scaling predicted by classical evolutionary models (27). These results suggest that second-order selection could play a previously unappreciated role in driving the success of genetic variants in large microbial populations, from laboratory evolution experiments (8, 10, 13, 30, 49) to natural systems like cancer (14, 15), influenza (31), or SARS-CoV-2 (34, 50). This could have important consequences for evolutionary forecasting (32, 51–53), since it implies that the fitness effects of such variants might fail to explain their long-term evolutionary success.

Our theory indicates that it could be difficult to detect these evolvability differences using traditional metrics like the rate of adaptation (8, 54, 55) or the substitution rate (49), since the associated changes in these observables are not always large (SI Section 4.1.7). Previously documented cases like Fig. 1A might therefore only represent a fraction of the selectable variation in evolvability. Future efforts could instead focus on mapping the aggregate changes to the distribution of fitness effects (Fig. 1B), e.g. using barcoded lineage tracking (9, 56–58) or mutation trap experiments (59). However, we have also shown that the important changes in this distribution will often occur in its high-fitness tail, and are poorly captured by existing heuristics like the mean mutational effect (27, 60). They can also depend on external factors like the population size and the overall mutation rate. This suggests that the evolvability benefits of a mutation should not be viewed as an intrinsic property of the cell, but rather a collective effect that can vary across populations or within the same population over time. Our theory provides a framework for predicting where these evolutionarily important differences will occur.

While we have primarily focused on the supply of beneficial mutations, our results can also be extended to account for changes in the supply of strongly deleterious mutations (|*s*| ≫ *v*/*x*_*c*_; SI Section 6), which behave like an effective direct cost, 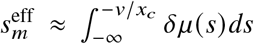 (Fig. S2). This mapping helps us understand how rapidly adapting populations balance the trade-offs (or synergies) between robustness and evolvability (61–63). It implies that rapidly adapting populations will be more willing to sacrifice short-term robustness for long-term gains in evolvability (Fig. 3A), but also that sufficiently large robustness gains can still be favored even if they eliminate the opportunities for future adaptation (Fig. 3B).

We have also assumed that the evolvability benefits of the modifier lineage remain fixed as it competes for dominance in the population. In reality, epistatic interactions could cause these benefits to attenuate – or even reverse – as the modifier acquires further mutations. Our heuristic analysis suggests that our current results will continue to hold as long as *μ*(*s*) and *μ*_*m*_(*s*) remain constant over a typical fixation time (roughly ∼*v*/*x*_*c*_ generations, or ∼*x*_*c*_/*s*_*b*_ additional mutations). This timescale is often modest in practice, allowing our minimal model to capture a broader range of epistatic scenarios than its idealized nature might originally suggest (Figs. S3 and S4; SI Section 7). Further extensions of this framework to allow for more rapidly varying distributions of fitness effects (“macroscopic epistasis”; 41, Fig. S1A) could be useful for understanding how large populations navigate rugged fitness landscapes (64).

Finally, while we have focused on the asexual dynamics common in laboratory experiments (8) and somatic evolution (14, 15), natural microbial populations often exhibit some degree of recombination (65). Widespread recombination will alter our predictions by decoupling the modifier locus from the future mutations that it produces (23, 66). Previous work suggests that some of our results may still apply on short genomic distances that remain tightly linked over the characteristic fixation time 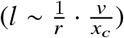 (67–69). However, modifiers can also benefit from transient linkage to mutations outside these asexual linkage blocks (70), and the fitness effects of the mutations could change when decoupled from the background that produced them (64). Understanding the interplay between these forces will be critical for understanding how second-order selection acts in other contexts.

## Acknowledgments

We thank Daniel Wong for useful discussions, and Sophie Walton, Olivia Ghosh, and Zhiru Liu for comments and feedback on the manuscript. This work was supported in part by the Alfred P. Sloan Foundation (FG-2021-15708) and a Terman Fellowship from Stanford University. B.H.G. is a Chan Zuckerberg Biohub – San Francisco Investigator.

## Author contributions

Conceptualization: J.T.F and B.H.G.; theory and methods development: J.T.F and B.H.G..; analysis: J.T.F and B.H.G.; writing: J.T.F. and B.H.G.

## Data and materials availability

Source code for forward-time simulations, numerical calculations, and figure generation are available at Github (https://github.com/bgoodlab/evolution_of_evolvability).

## Supplementary Information

**Fig S1.**
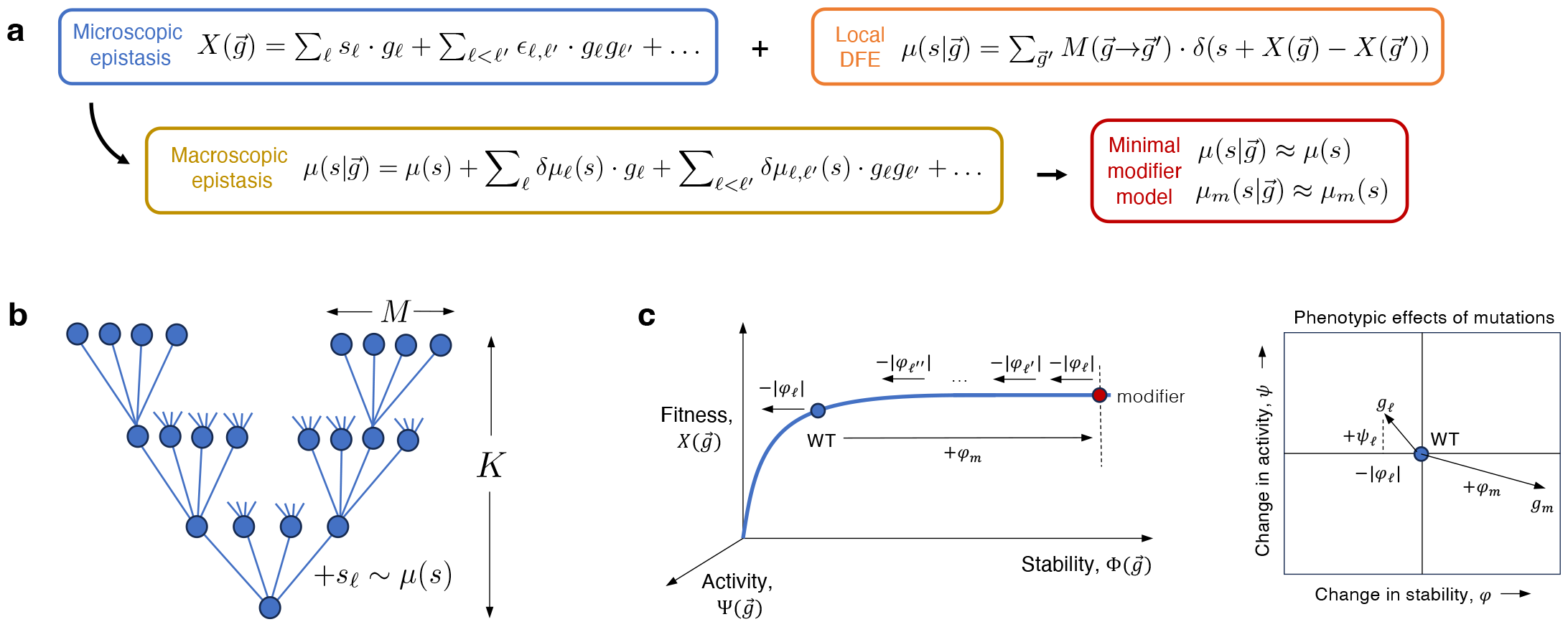
Examples of epistatic fitness landscapes that satisfy the minimal model in Fig. 1. (a) The evolvability modifier in Fig. 1B can be viewed as the lowest order term in a general macroscopic epistasis expansion (left, SI Section 3.1). Different fitness landscapes can produce the same macroscopic behavior. (b,c) Examples of highly epistatic fitness landscapes that satisfy the simple model above. (b) A “maximally epistatic” landscape of branching uphill paths, which generalizes the model in Ref. (71). Each step *k* = 1, …, *K* of a given path can access *M* ≪ *L* beneficial mutations; all other genotypes have fitness zero. (c) A fitness landscape formed by a non-linear combination of two global phenotypes, e.g., stability, 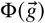, and activity, 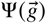. Individual mutations can affect both traits simultaneously (right). Stabilizing mutations can act like modifier alleles by potentiating the fitness benefits of mutations that would destabilize the protein on their own (left). In particular, a strongly stabilizing mutation can allow *K* ≈ *ϕ*_*m*_ /|*ϕ*_*ℓ*_| new mutations to accumulate before their effects on stability become important. See SI Section 3.1 for more details.

**Fig S2.**
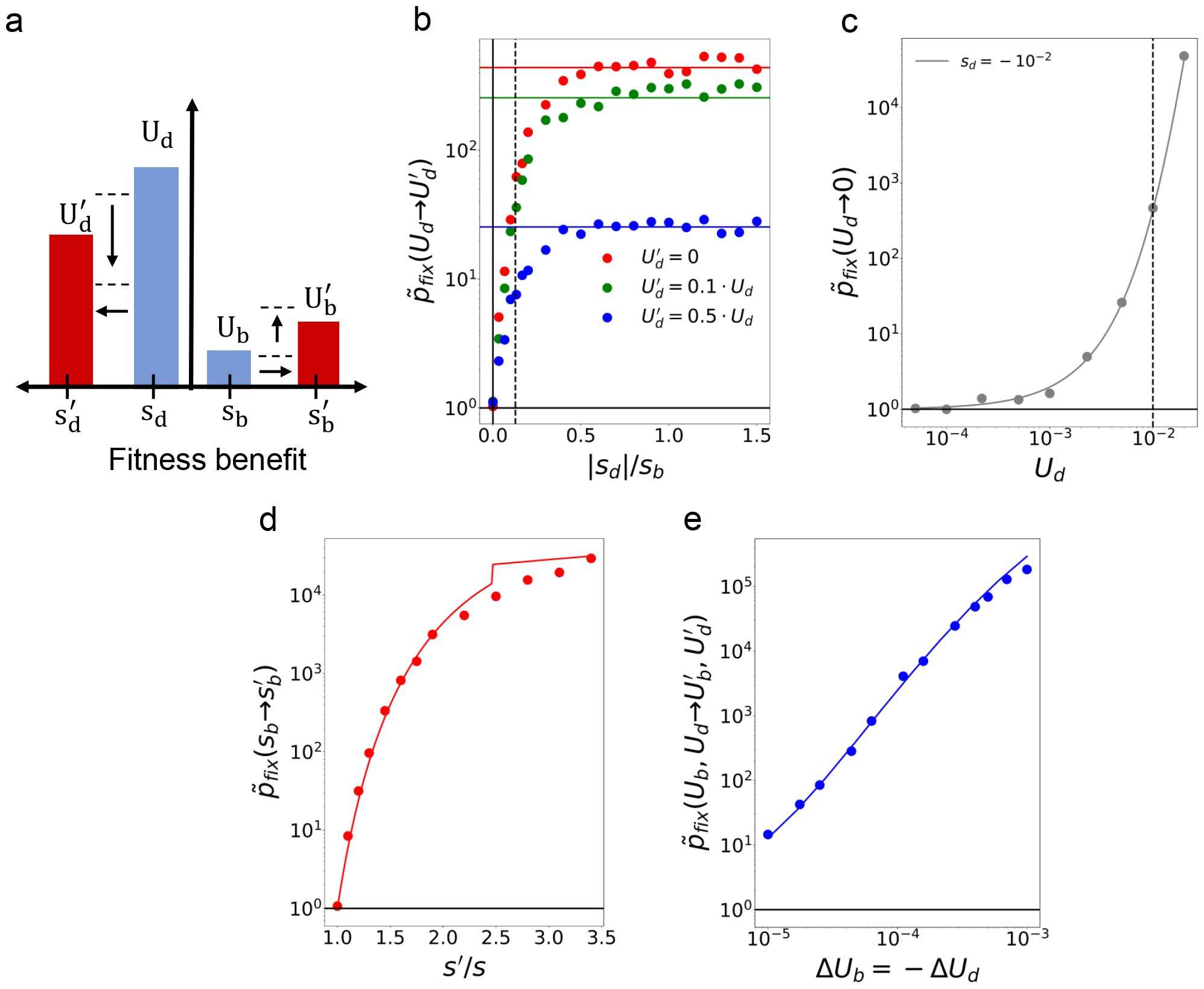
Deleterious mutations and second-order selection for robustness. **(a)** A generalization of the simplified model in Fig. 2A that incorporates deleterious mutations with a characteristic cost *s*_*d*_ and rate *U*_*d*_. A modifier mutation shifts these parameters to a new combination, 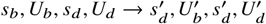. **(b, c)** Fixation probability of a robustness-enhancing modifier with 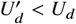 (and all other parameters are held fixed). Symbols denote the results of forward time simulations for *N* = 10^8^, *s*_*b*_ = 10^−2^, and *U*_*b*_ = 10^−5^, while solid lines denote our theoretical predictions in SI Section 6. Panel (b) shows that the purgeable mutations approximation holds across a broad range of fitness costs, with the dashed line marking the predicted transition to quasi-neutrality (|*s*_*d*_| ≈ *v*/*x*_*c*_). Panel (c) shows that selection for increased robustness is relatively weak unless *U*_*d*_ ≳ *s*_*b*_ (dashed line). **(d)** Fixation probability of a modifier that imposes a tradeoff between robustness and evolvability by increasing the strength of selection on beneficial and deleterious mutations simultaneously. Results are shown for *s*_*b*_ = |*s*_*d*_| = *s* and *U*_*d*_ = 10^−2^, with the remaining parameters the same as panel b. Since *U*_*d*_ ≫ *U*_*b*_, this example shows that strong selection for evolvability can occur for modifiers reduce the average fitness effect of mutations 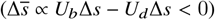. **(e)** Fixation probability of modifier that enhances robustness and evolvability at the same, by shifting mutations from deleterious to beneficial 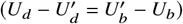. Symbols denote results of forward-time simulations with *s*_*d*_ = −10^−2^ and *U*_*d*_ = 10^−2^, with the remaining parameters the same as panel (b). Lines denote our theoretical predictions in the absence of deleterious mutations 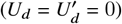. This example shows thatenhancements in evolvability are weighted more strongly than comparable increases in robustness, even when nearly all new mutations are deleterious (*U*_*b*_ ≪ *U*_*d*_).

**Fig S3.**
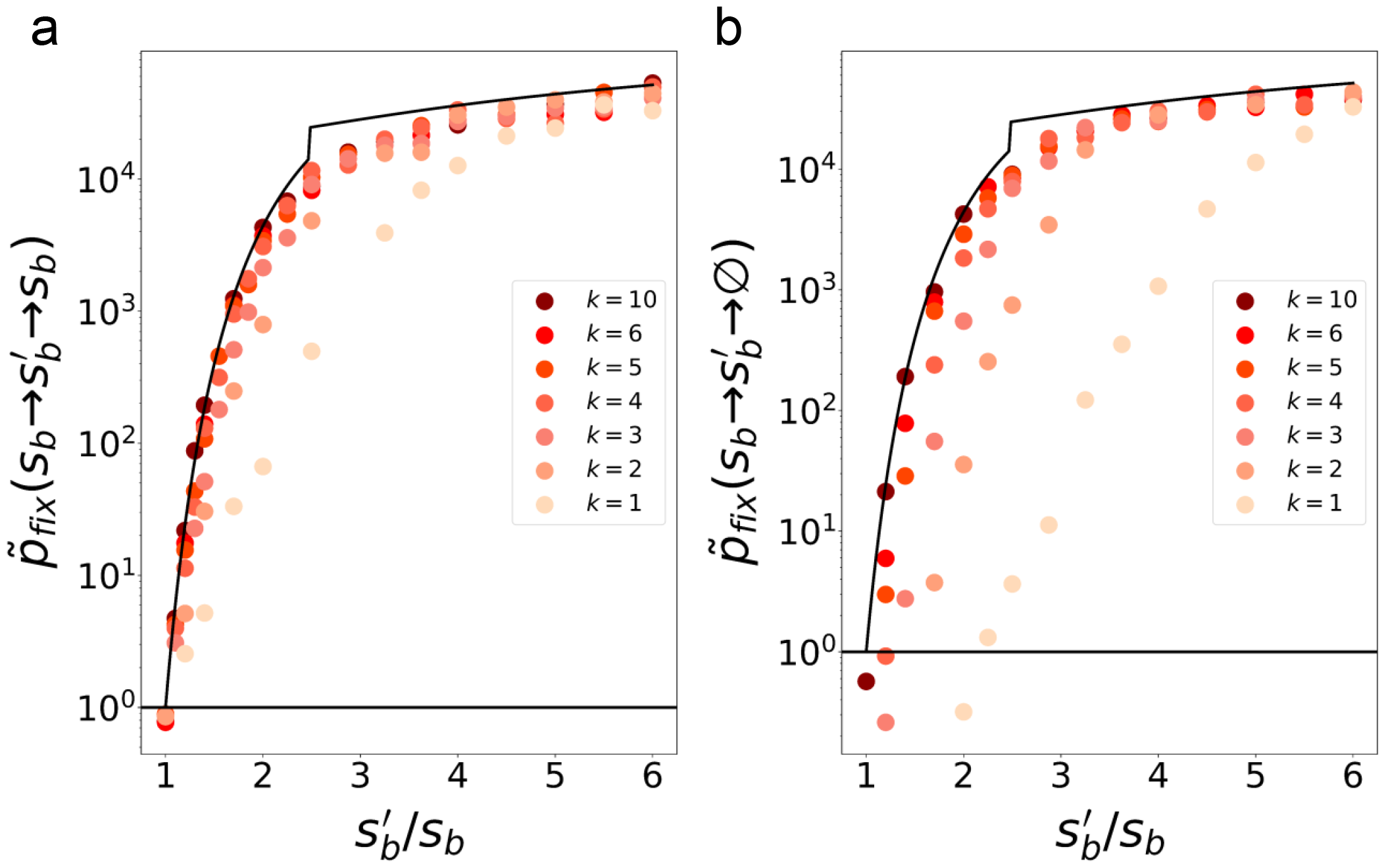
Relaxing the assumption that modifiers permanently change the mutation spectrum. **(a)** A generalization of the model in Fig. 2B, where the modifier reverts back to the wildtype distribution of fitness effects after acquiring *K* additional mutations (SI Section 7.2). Symbols denote the results of forward-time simulations for *N* = 10^8^, *s*_*b*_ = 10^−2^, and *U*_*b*_ = 10^−5^, while the line denotes our theoretical predictions for the minimal modifier model in Fig. 1B (i.e. *K* = ∞). Consistent with our calculations in SI Section 7.2, these results show that the *K* ≈ ∞ approximation remains highly accurate even for moderate values of *K* (e.g. 2-3), and for as little as a single mutation (*K* = 1) in the quasi-sweeps regime.**(b)** An alternative version of the model in panel a, where the modifier reverts to an evolutionary dead-end (*μ*_*m*_ = ∅) after *K* mutations; all other parameters are the same as panel a. Even in this extreme case, our minimal modifier model (solid line) remains highly accurate for moderate values of *K*, and as little as *K* = 1 in the quasi-sweeps regime. This demonstrates that large populations can only “see” across the fitness landscape for 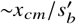 additional mutations (SI Section 7.2).

**Fig S4.**
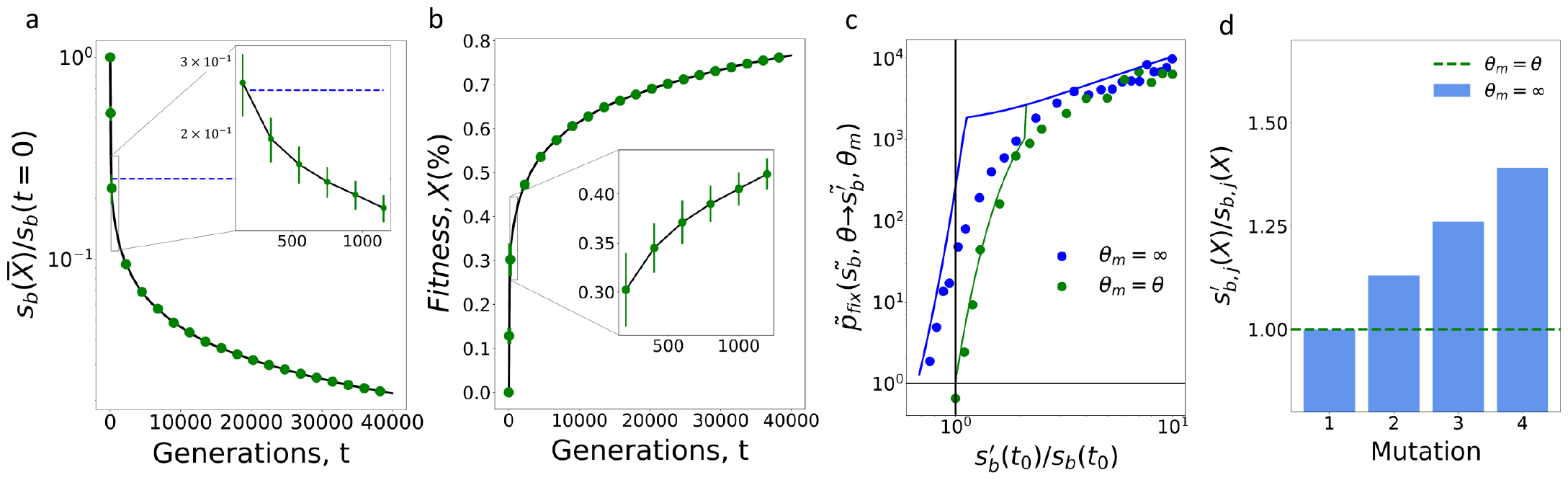
Selection for evolvability in the presence of diminishing returns epistasis. **(a, b)** A simple model of global diminishing returns epistasis motivated by the empirical example in Ref. (55) (SI Section 7.3). The fitness effects of new mutations shrink as the population adapts (panel a), leading to a decelerating rate of adaptation over time (panel b). Points denote the results of forward-time simulations for the distribution of fitness effects 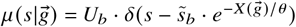, with 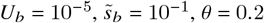, and *N* = 10^7^; points are connected by solid lines to aid visualization. **(c, d)** The fixation probability of an evolvability modifier that arises at the beginning of the inset in panel a, where the fitness trajectory is still decelerating. Green symbols in (c) show a selection-strength modifier with the same diminishing returns schedule as the background population (*θ*_*m*_ ≈ *θ*), while the blue symbols show an alternate example where the modifier avoids future diminishing returns once it arises (*θ*_*m*_ ≈ ∞). The green line illustrates the predictions from the “adiabatic” approximation in SI Section 7.3, demonstrating that the permanent modifier model 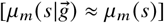 provides a good approximation when the local selection strengths are properly renormalized. The blue line shows the predictions from our heuristic analysis in SI Section 7.3, which accounts for the additional benefits that accrue for the modifier lineage when *θ*_*m*_ ≫ *θ* (panel d). This example illustrates that the evolvability advantages that accrue from large differences in diminishing returns epistasis can drive modest deviations from our existing theory when *θ* grows close to *x*_*c*_.

## 1 Comparison to deterministic modifier theory

In this section, we rederive a key result from classical modifier theory known as the “mean fitness principle” (20). This theory predicts that in an infinitely large asexual population, natural selection will favor modifiers that increase the long-term mean fitness of the population. Our derivation closely follows the one given in Ref. (21), as well as related work on mutators (72–74). We reproduce it here for completeness, using the same notation employed in our more general analysis below.

To establish this result, we note that in the absence of genetic drift (*N* = ∞), the deterministic dynamics of a well-mixed asexual population can be written in the general form,

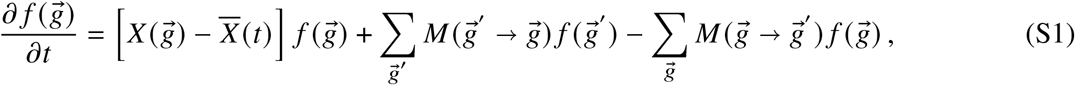

where 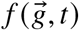 is the frequency of genotype 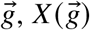 is the (log) fitness of genotype 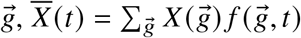 is the mean fitness of the population, and 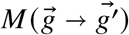 is the mutation rate from genotype 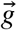 to 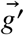. The fate of a general evolvability modifier allele can be analyzed by introducing an analogous set of equations,

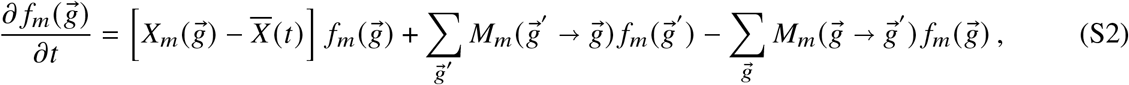

where 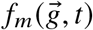 is the frequency of individuals with the mutant allele at the modifier locus and genotype 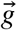 elsewhere, 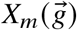 is genotype-to-fitness map for the modifier, 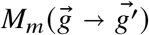 is the corresponding set of mutation rates, and the mean fitness in Eqs. (S1) and (S2) now sums over both the mutant and wildtype lineages,

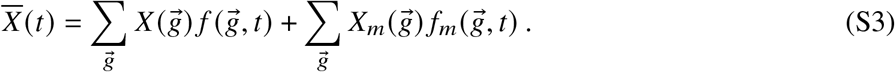

In principle, this model allows for arbitrary changes to arbitrary non-linear fitness landscapes, encapsulating all possible forms of epistasis (75).

The solutions to this coupled system of equations can be written in the general form

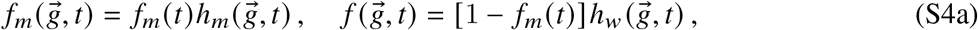

where *f*_*m*_(*t*), *h*_*m*_(*t*), and *h*_*w*_ (*t*) satisfy the related set of equations,

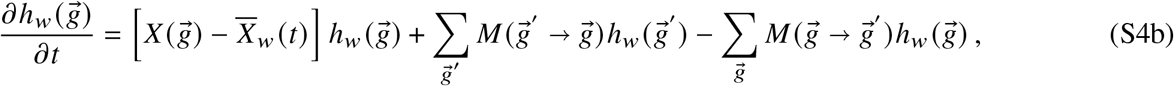

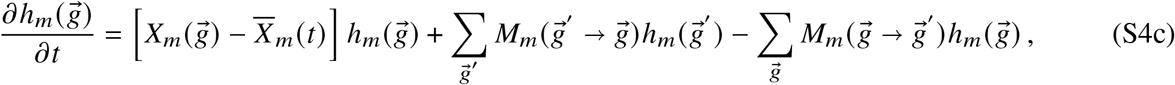

and

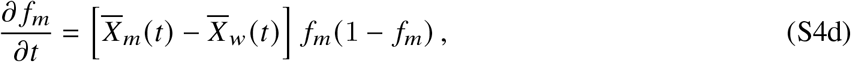

with

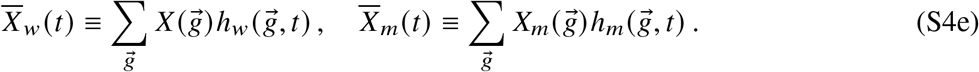

In this notation, 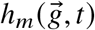 represents the re-normalized genotype distribution within the modifier lineage, 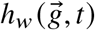 denotes the corresponding distribution within the wildtype, 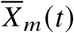 and 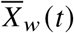 represent the mean fitnesses of each lineage, and *f*_*m*_ (*t*) denotes the total frequency of the modifier.

This change-of-variables shows that the total frequency of the modifier lineage only depends on the relative values of the mean fitnesses, 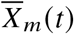 and 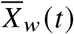, yielding the time-dependent solution,

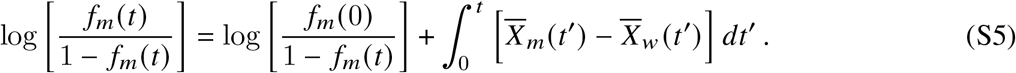

This shows that at long times (*t* → ∞) the lineage that will dominate the population is the one with the higher value of 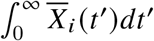. These mean fitnesses can be predicted from the dynamics of the intra-lineage frequencies, 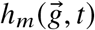 and 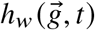, which decouple from each other and from the total size of *f*_*m*_(*t*). In most cases of interest, the mean fitness of each lineage will approach an equilibrium value, 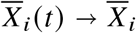, which may be a complicated function of 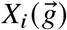 and 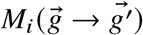, but is otherwise independent of the surrounding population. Equation (S5) then shows that the modifier will take over the population if and only if it increases the equilibrium mean fitness. This generalizes the “mean fitness principle” derived in previous work (20).

However, this deterministic calculation neglects two key factors that are relevant for any large but finite population. First, the mean-field dynamics in Eqs. (S1) and (S2) neglect the random occurrence of new mutations and the stochastic fluctuations they experience while rare. These fluctuations can dramatically influence the dynamics of the mean fitness – particularly when multiple beneficial mutations are available (47). In addition, the deterministic calculation neglects the possibility that the mutant or wildtype lineage may fix before their long-term benefits in Eq. (S5) are fully realized. As we will see below, both of these effects will become extremely important in the adapting populations that we analyze in this work. Interestingly, we will see that in some cases, it will be possible to account for these effects in an approximate manner by inserting a upper limit on the integral in Eq. (S5) (SI Section 4.1.7), providing a conceptual link between classical modifier theory and the more complex scenarios studied in this work. Identifying such cases and their appropriate time horizons is the goal of the next several sections.

## 2 Evolvability modifiers in the successive mutations regime

Another useful limit occurs in small populations, where the production of beneficial mutations is sufficiently rare that adaptation proceeds via a sequence of discrete selective sweeps. In this regime, the fate of a given modifier will strongly depend on its ability to generate the next beneficial mutation. We can formalize this idea by defining the local distribution of fitness effects (DFE),

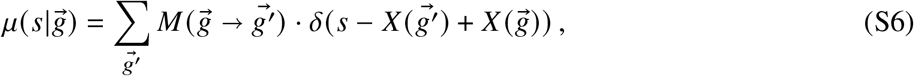

which tabulates the fitness effects of all the mutations that can be accessed from a given genotype 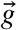. Modifier individuals will have their own corresponding set of DFEs,

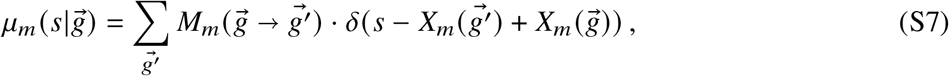

along with a direct cost or benefit

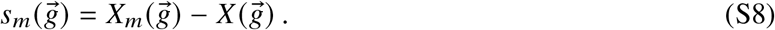

The distributions in Eqs. (S6) and (S7) can be computed for any epistatic fitness landscape, and will constitute the key input parameters in our analysis below.

In the absence of any modifier mutations, the evolutionary dynamics of the successive sweeps regime are well described by previous work (47). The wildtype population produces new beneficial mutations at a total rate 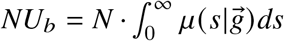 per generation. Most of these mutations will drift to extinction before they reach appreciable frequencies. However, with probability *p*_est_≈2*s*, a lucky mutant will fluctuate to a sufficiently large frequency (*f* ≈1/2*Ns*) that it begins to grow deterministically [*∂*_*t*_ *f* ≈ *s f* (1 − *f*)], and will sweep through the population on a timescale of order 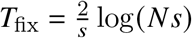. The population will produce these successful mutations at a total rate 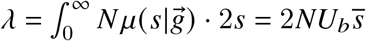 per generation, where 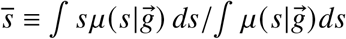 is the average beneficial fitness effect that is accessible to the current genotype. This implies that the typical waiting time for the next sweep event is 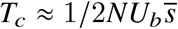 generations. The condition that successive sweeps will not interfere with each requires that *T*_fix_ ≪ *T*_*c*_, which requires that 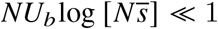 (47). This limit is also known as the “strong selection, weak mutation regime” (SSWM).

A general evolvability modifier (*μ* → *μ*_*m*_) will produce a change in the beneficial mutation rate 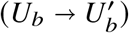 as well as the typical fitness benefit 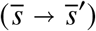. (It may also change the spectrum of deleterious mutations, which we neglect for the time being; see SI Section 6). The mutator version of this model was previously analyzed in Ref. (24), as well as a number of related studies (27, 73, 76–85). We reproduce these calculations below, while generalizing them to allow for changes in the average fitness benefits of mutations (similar to Ref. 27) as well as a broader range of direct costs and benefits. These classical results will provide a useful baseline for understanding the impact of larger populations, which we analyze in SI Section 3 below.

In the successive mutations regime, a newly arising modifier lineage will initially compete with the wildtype population according to the single-locus dynamics

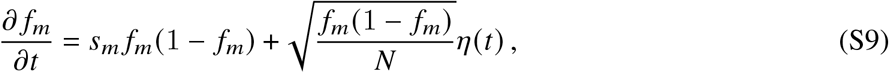

where *η*(*t*) is a Browian noise term (86) and *s*_*m*_ is the direct cost or benefit of the modifier. The next selective sweep will now be generated by a pair of competing Poisson processes with rates

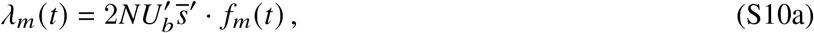

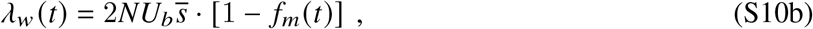

which correspond to the mutant and wildtype lineages, respectively. Since the mutation rate and fitness benefit both contribute linearly to *λ*_*i*_ (*t*), this process is formally equivalent to the mutator scenario analyzed in (24), with 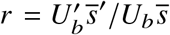 replacing 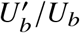. This allows us to conclude that in absence of a direct cost (*s*_*m*_ = 0), the fixation probability of the modifier scales as

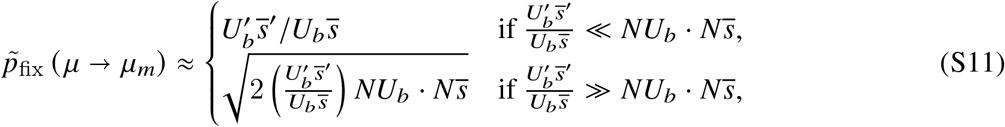

provided that *T*_*c*_ ≪ *N* (or 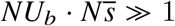). When 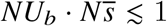, the modifier will either fix or go extinct neutrally (*p*_fix_ ≈ 1/*N*) before the next sweep occurs. This shows that second-order selection is more efficient in larger populations, which is reminiscent of our results in Fig. 2. In this case, however, the benefits of second-order selection are capped by the ratio 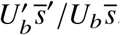, which implies that very large changes in 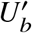 or 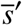 are required to produce an appreciable fixation probability.

A similar application of Ref. (24) shows that the fixation probability of a modifier with a direct cost (*s*_*m*_ *<* 0) scales as

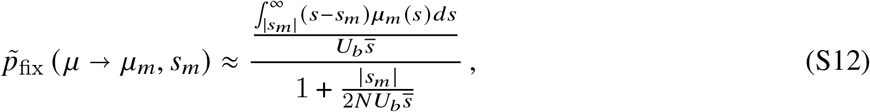

provided that 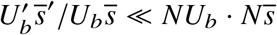. The integral in the numerator implies that modifiers with direct costs larger than 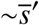 will have dramatically reduced fixation probabilities, since future beneficial mutations in these backgrounds will still be less fit than the current wildtype population. However, since *NU*_*b*_ ≪ 1, the fixation probability of the modifier will be significantly reduced by the denominator term well before these “wasted opportunities” start to become relevant. This illustrates how even small direct costs – much smaller than a single driver mutation – can overwhelm the evolvability benefits of mutations in the successive sweeps regime.

The fixation probability of a modifier with a direct benefit *s*_*m*_ *>* 0 can be computed using a similar procedure. When 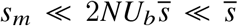, the next sweep will typically occur before the modifier lineage establishes, so an analogous version of Eq. (S12) still holds:

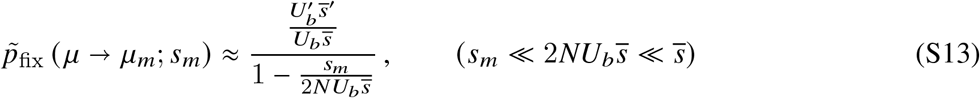

For stronger fitness benefits 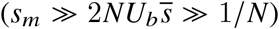, the modifier will have the opportunity establish and grow deterministically before the next sweep occurs:

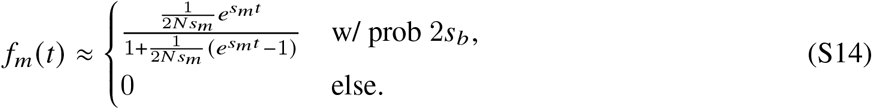

In this regime, the fixation process is similar to the “first-step” clonal interference analysis in Ref. (87). Provided that 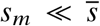, the additional fitness benefit of the modifier will have a negligible impact the establishment probability of the next sweep in either genetic background, so that

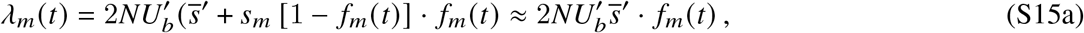

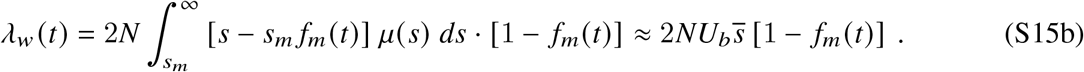

The fixation probability is determined by these competing Poisson processes, so that

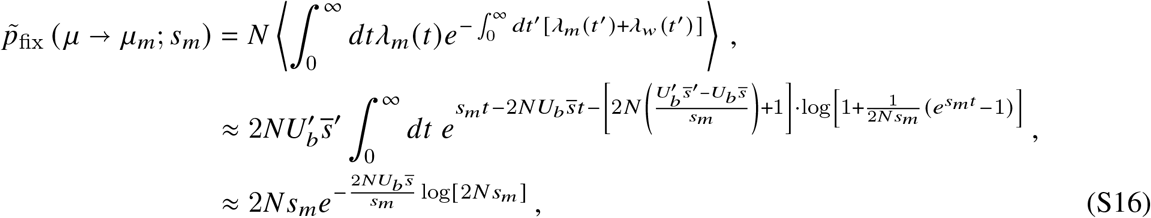

which will be valid in the limit that 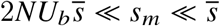. This result has a simple heuristic interpretation as the establishment probability of the modifier multiplied by the probability the wildtype population does not generate a sweep before the modifier fixes on its own (87). Since this fixation probability is independent of *μ*_*m*_(*s*), it implies that even small direct benefits — much smaller than the size of a typical driver mutation — can override the effects of second order selection, even if they reduce the long-term rate of adaptation to zero. Together with the direct cost results above, these calculations emphasize that natural selection can be extremely sensitive to the short-term costs or benefits of a mutation in the successive sweeps regime, in contrast to what we observe in larger populations like Fig. 1A.

## 3 Evolvability modifiers in the clonal interference regime

In larger populations (*NU*_*b*_ ≳ 1), the assumption of discrete selective sweeps will start to break down. Multiple beneficial lineages will segregate in the population at the same time, and will interfere with each other as they compete for dominance in the the population (47). In this *clonal interference regime*, the fate of a given mutation will sensitively depend on the genetic background that it arises on, and the future mutations that its descendants produce before they fix or are driven to extinction. This requires a stochastic generalization of the full multi-locus dynamics in Eqs. (S1) and (S2),

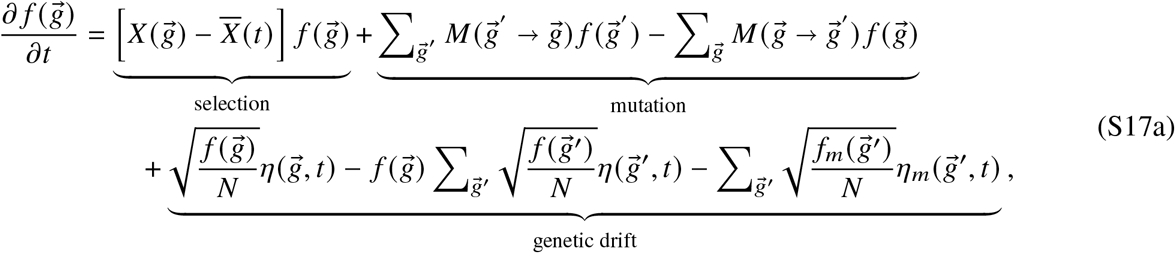

and

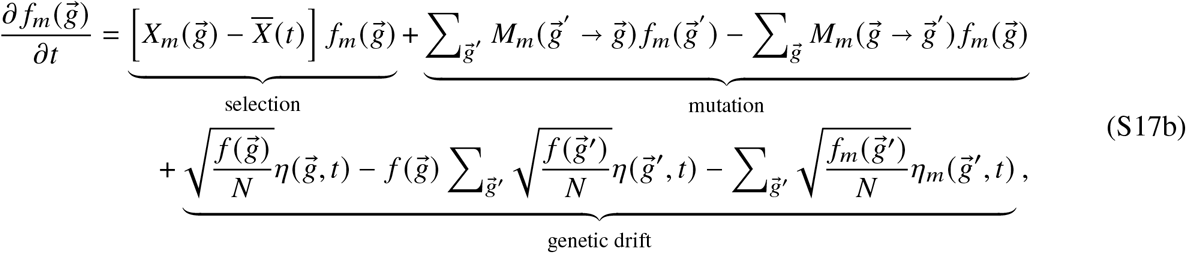

where 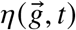 and 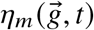 are uncorrelated Brownian noise terms (88). While this model is straightforward to write down, there are no known solutions for arbitrary choices of 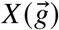 and 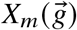. Further progress requires us to make specific assumptions about the shapes of these fitness landscapes, as well as the corresponding mutation kernels 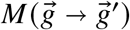 and 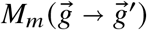. We introduce one particularly convenient parameterization in the next section, which is motivated by the concept of *macroscopic epistasis* explored in previous work (41, 89).

### 3.1 Macroscopic epistasis expansion

There are many possible fitness landscapes one could consider. These are often parameterized as a power series involving different combinations of loci,

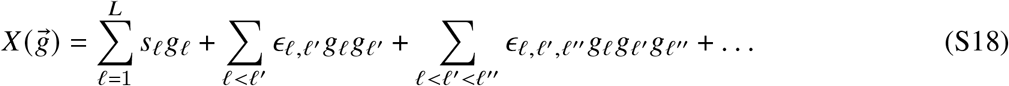

where *g*_*ℓ*_ = 1 if there is a mutation at site *ℓ* and 0 otherwise (75). This representation can be viewed as a Taylor expansion around a perfectly smooth fitness landscape, where all the {*ϵ*_*ℓ*,…,*ℓ*_′} coefficients vanish. Nonzero values of the *ϵ*_*ℓ*,…,*ℓ*_′ coefficients correspond to epistatic interactions between loci; following previous work (41), we will refer to these interactions as *microscopic epistasis*, since they can in principle vary across all possible combinations of sites.

Modifier alleles can be expressed in this framework by designating an arbitrary site as the modifier locus (e.g. *ℓ* = *m*), and recalculating the landscape for *g*_*m*_ = 1 (modifiers) and *g*_*m*_ = 0 (wildtype) separately. This notation makes it clear that a modifier that changes the fitness effects of other mutations must necessarily involve some microscopic epistasis, corresponding to terms like *ϵ*_*m,ℓ*_, *ϵ*_*m,ℓ,ℓ*_′, and so on.

The space of epistatic fitness landscapes is enormous, and their impact on the evolutionary dynamics of large populations is not well understood (64). Some studies have attempted to navigate this complexity by truncating Eq. (S18) after the pairwise terms (90); others have focused on smaller landscapes containing just a handful of interacting loci (91, 92). In our analysis below, we will show that it will be useful to consider an alternative limit of Eq. (S18), in which the local DFEs defined in Eqs. (S6) and (S7) are approximately constant for different genotypes:

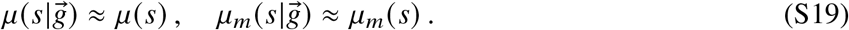

We can view this approximation as the lowest order term in an alternative expansion of the fitness landscape,

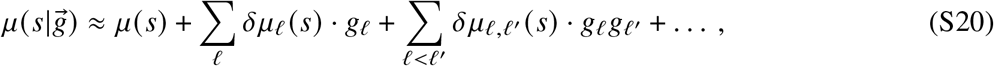

which works directly in the DFE basis. This genotype dependence of the DFE is sometimes known as *macroscopic epistasis* (41), since it aggregates over a large number of microscopic interactions in Eq. (S18). The approximation in Eq. (S19) can therefore be viewed as the simplest possible model that incorporates some amount of macroscopic epistasis, with a non-zero *δμ*_*ℓ*_ (*s*) term at the modifier locus (*ℓ* = *m*) and all other *δμ*_*ℓ*,…,*ℓ*_′ terms vanishing.

We note that such macroscopic epistasis can arise even in the absence of microscopic epistasis (*ϵ*_*ℓ*,…,*ℓ*_′ = 0) if the modifier alters the mutation rates at other loci. For example, mutator alleles are often modeled as a simple change of scale,

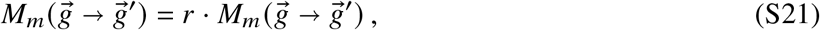

leading to a proportional change in the DFE,

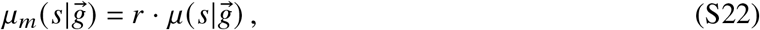

which has been the starting point for many previous studies (24, 26, 85). In practice, mutator strains are typically biased toward specific types of mutations, so that the simple proportional model in Eq. (S21) does not necessarily apply (4, 6, 27). Other mutagenesis mechanisms that target specific genomic regions (93, 94) lead to conceptually similar complications. Both require us to consider changes in the shape of the DFE in addition to its overall scale (4, 6, 27).

In addition to mutator alleles, Eq. (S19) can also emerge from epistatic interactions between loci. The simplest example is a pairwise epistasis model, with nonzero values for the coefficients involving *ℓ* = *m*, and zeros everywhere else. This is sufficient to recover the limit in Eq. (S19), but it is not the only possibility: any other landscape that gives rise to the same overall distributions in Eq. (S19) will exhibit similar dynamics, even if the individual fitness effects 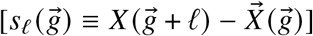 are undergoing more complicated rearrangements under the hood.

Moreover, our results will not require Eq. (S19) to hold across the entire fitness landscape, but only within a smaller region that is explored before the modifier either fixes or goes extinct. We determine the size of this local neighborhood in SI Section 7 and find that it is often modest, corresponding to just a handful of mutational steps for many empirically relevant parameter values (Fig. S3). We will also show that the assumption in Eq. (S19) is most sensitive to a narrow range of beneficial fitness effects, so that substantial deviations in other parts of the DFE can still have a negligible impact on the modifier lineage (SI Section 7). At present, it is difficult to enumerate all of the microscopic landscapes that are consistent with a given DFE function, 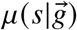. However, we can still identify several examples that satisfy Eq. (S19) – at least in the approximate sense required – that go beyond the mutator allele and pairwise epistasis examples above:

#### Branching epistatic landscapes

At a formal level, we can consider a “maximally epistatic” landscape of branching uphill paths of length *K*, where each step *k* of a given path can access *M* ≪ *L* other beneficial mutations (Fig. S1A). If the uphill paths do not contain any loops, the fitness function can be expressed in the form

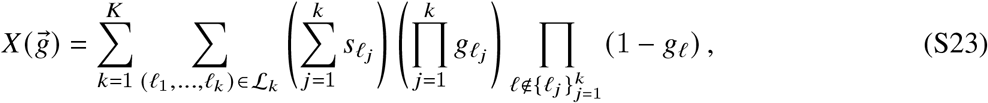

where ℒ_*k*_ denotes the set of all uphill sub-paths of length *k*, and the *s*_*ℓ*_ coefficients are independently drawn from the target DFE *μ*(*s*). This construction ensures that the DFEs calculated from Eq. (S23) will satisfy 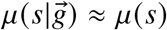 when *M* is large, even though the landscape contains large amounts of microscopic epistasis.

We can extend this construction to include a modifier allele by writing

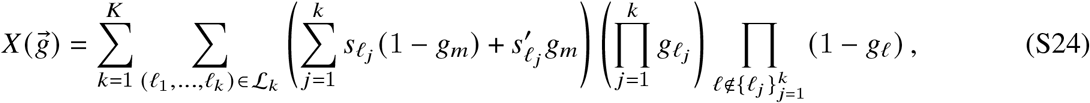

where the 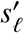 coefficients are independently drawn from the modifier DFE *μ*_*m*_(*s*). This microscopic landscape satisfies the macroscopic epistasis approximation in Eq. (S19), but includes many non-pairwise interactions by construction.

#### Non-linear global phenotypes (e.g. stability-activity tradeoffs)

The approximation in Eq. (S19) can also emerge in more concrete settings, when the fitness of the organism is a non-linear function of multiple global phenotypes. A prototypical example is the stability-activity tradeoff that is often observed in viral evolution (95, 96) and other protein-binding problems (97), where stabilizing mutations can potentiate the fitness benefits of mutations that would destabilize the protein on their own. The essential ingredients of this behavior can be captured in a simple model containing two global phenotypes: (i) activity, denoted by 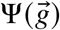, which contributes additively to the total fitness, and (ii) stability, denoted by 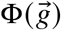, which has a Boltzmann-like contribution,

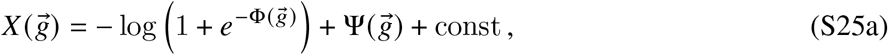

We will assume that both phenotypes can be expressed as additive functions of the genotype,

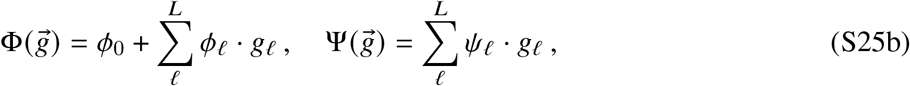

where *ϕ*_0_ denotes the stability of the reference strain. In this model, mutations that only increase the activity of the protein are always beneficial (*s*_*ℓ*_ ≈ *ψ*_*ℓ*_). However, mutations that increase activity while also decreasing stability can be either costly or beneficial depending on the stability of the background that they occur on.

Suppose that there are a large number (*M*) of such mutations with |*ϕ*_*ℓ*_| − *ϕ*_0_ ≫ 1, which implies that they will be deleterious on the wildtype background. In this scenario, any mutation that strongly increases stability will function like an evolvability modifier, by allowing these previously deleterious mutations to occur. In particular, if the stability enhancement *ϕ*_*m*_ is much larger than |*ϕ*_*ℓ*_| − *ϕ*_0_, then the fitness benefits of the unleashed mutations will be approximately constant (*s*_*ℓ*_ ≈ *ψ*_*ℓ*_) until roughly *K* ≈ *ϕ*_*m*_ /(|*ϕ*_*ℓ*_| − *ϕ*_0_ such variants have accumulated. If *M* ≫ *K* ≫ 1, this example will satisfy the approximation in Eq. (S19), while relying on the higher-order epistatic interactions in Eq. (S25).

While this particular example was motivated by protein stability, similar behavior can occur for other combinations of phenotypes, as long as they include the appropriate non-linearities. For example, a simple model of stabilizing selection involving a one nearly optimized phenotype and another non-optimized one can be expressed as

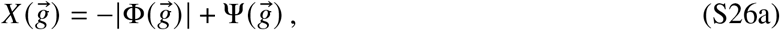

where

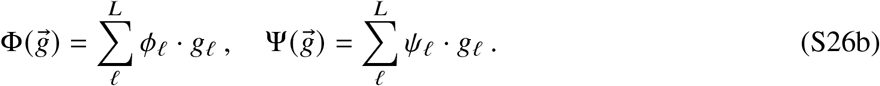

A mutation that increases 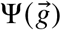 while displacing 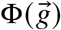 from its optimal value would enable *K* ≈ *ϕ*_*m*_/*ϕ*_*ℓ*_ previously deleterious mutations to accumulate — each providing a fitness benefit *s*_*ℓ*_ ≈ *ϕ*_*ℓ*_ — before the phenotypic optimum is reattained. If *K* is sufficiently large, this evolvability modifier would also satisfy the approximation in Eq. (S19), while involving a distinct form of non-pairwise epistasis.

#### Chromosomal duplications / aneuploidy

The approximation in Eq. (S19) can also apply to scenarios that are difficult to capture with a traditional fitness landscape, because they involve changes to the structure of the genome itself. Classical examples include chromosomal duplications and other copy-number changes, which are frequently observed in cancer evolution (14, 98) and laboratory evolution experiments in eukaryotes (99). These copy number variants involve changes in ploidy — in addition to changes in target size — so that dominance effects start to become important. For example, a whole-genome duplication of a haploid genome would lead to a modified DFE of the form

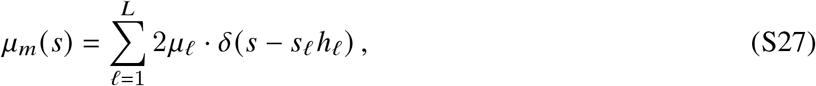

where *h*_*ℓ*_ denotes the dominance coefficient of the mutation at site *ℓ*. The successive sweeps picture in SI Section 2 predicts that the fixation probability of this copy-number variant is 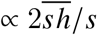, which is neutral in purely semi-dominant case 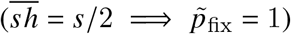 and only moderately beneficial for complete dominance 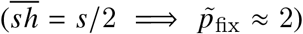. In contrast, our results below suggest that in larger populations, these variants can be both strongly favored or disfavored by second-order selection depending on the joint distribution of (*s*_*ℓ*_, *h*_*ℓ*_).

Together, these examples illustrate that the limiting behavior in Eq. (S19) — while still capturing just a subset of all possible fitness landscapes — can nevertheless approximate many biologically relevant scenarios where second-order selection is thought to play an important role. We will therefore use this model as a starting point for all of our mathematical derivations below. We will revisit this assumption in SI Section 7, where we discuss extensions to other possible fitness landscapes.

### 3.2 Fitness wave formalism

An advantage of the model in Eq. (S19) is that it allows us to exploit existing “fitness wave” methods for modeling clonal interference (24, 39, 40, 46, 47, 69, 100–103). This previous literature has shown that the multi-locus dynamics in Eq. (S17) can often simplified by considering a coarse-grained picture, which groups together individuals with the same overall fitness:

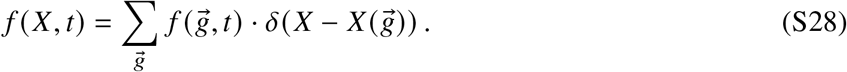

We can extend this idea to our present context, defining a corresponding fitness distribution for the modifier lineage as well:

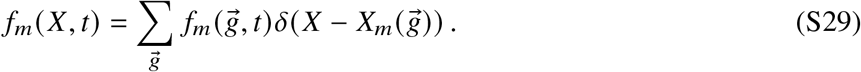

In the special case that 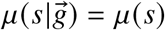 and 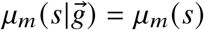 for all genotypes 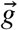, Eq. (S17) can be written in the coarse-grained form,

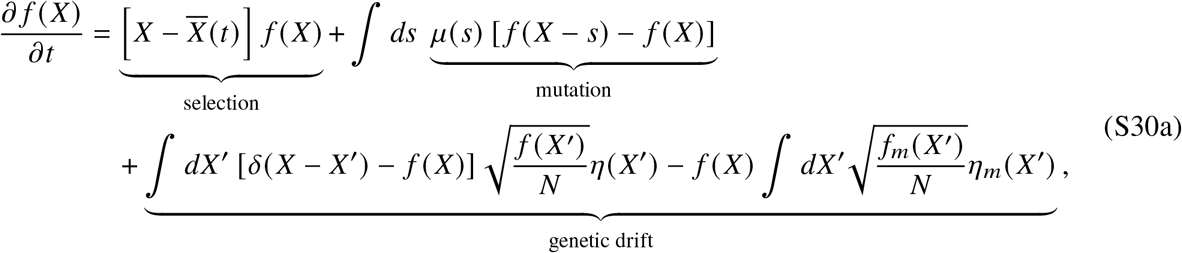

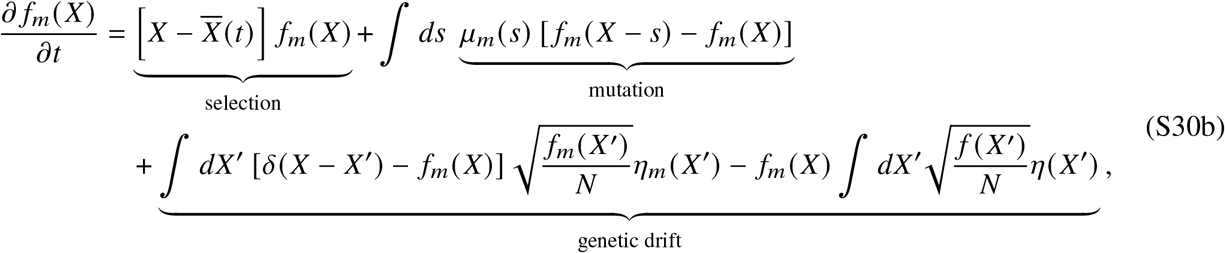

where 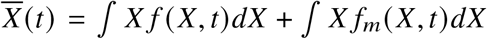 is the mean fitness of the population, and the *η*_*i*_ (*X*) are uncorrelated Brownian noise terms (88). This allows us to generalize the notion of a mutator allele to handle more general differences in evolvability, while still bypassing the enormous complexity of the underlying fitness landscape.

In the absence of the modifier (*f*_*m*_ = 0), the wildtype distribution *f* (*X, t*) approaches a traveling wave form that increases in fitness at an average rate 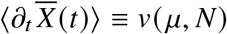 (39, 47). Previous work has shown that the typical profile *f* (*x*) is well-approximated by the deterministic equation,

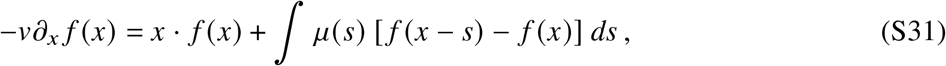

which is the expected value of Eq. (S30a) when 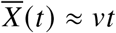 (39, 40).

*We assume that the modifier mutation will arise in this steady-state population, on a genetic background drawn from f*(*x*). Its descendants will found a second fitness wave, *f*_*m*_(*X, t*), which competes with the wildtype population *f*(*X, t*) as they continue to acquire additional mutations according to the dynamics in Eq. (S17) (Fig. 1).

#### Branching process approximation

The key approximation we will make in this work is that the fate of the modifier will often be determined while it is still at a low frequency in the population. Previous studies have shown that this is a good approximation for first-order mutations – including both neutral and deleterious mutations – in the clonal interference regime (39, 40, 43, 46, 103, 104). In this work, we use a combination of self-consistency arguments and comparisons with simulations to show that this same approximation holds for a broad range of modifier alleles as well. The main exceptions will occur for modifiers that dramatically reduce evolvability (e.g. the dead-end modifiers in Fig. 3D); we treat this case separately in SI Section 4.1.6.

When the frequency of the modifier is small, the mean fitness of the population can be approximated by the wildtype value 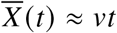, and the higher-order contributions in the drift term in Eq. (S30b) can be neglected. The dynamics of the modifier then reduce to the linear branching process,

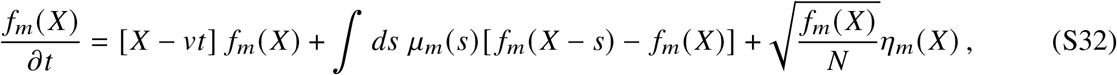

subject to the initial condition *f*_*m*_ (*X, t*) = *δ*(*X x s*_*m*_). The non-extinction probability of this process satisfies the standard branching process recursion,

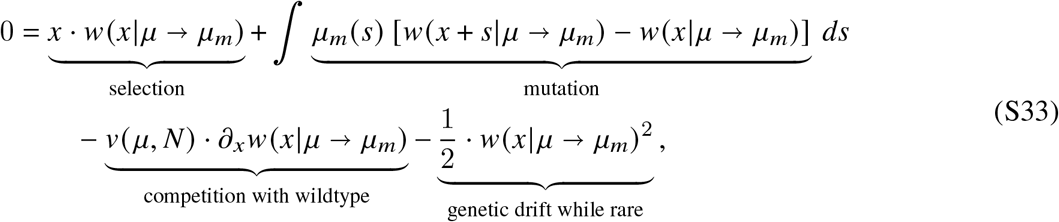

which is a straightforward generalization of the mutator version in Ref. (24). If the fate of the modifier is determined while it is rare, then this non-extinction probability must also coincide with the long-term probability of fixation. This implies that the overall fixation probability of the modifier can be obtained from Eq. (S33) by averaging over the random genetic background,

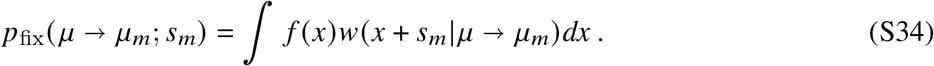

In the limit that *μ*_*m*_(*s*) → *μ*(*s*), Eq. (S34) reduces to the fixation probability of a first-order mutation which has been studied in previous work (39, 40, 46, 103). In particular, this previous work has shown that a self-consistency condition can be obtained from the fact that the overall fixation probability of a neutral mutation must always be equal to 1/*N* (this follows directly from the normalizability of the full dynamics in Eq. S30). Combining this fact with Eq. (S34) yields the self-consistency condition,

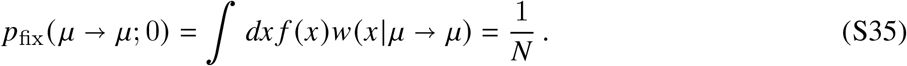

Together with Eqs. (S31) and (S33), this completely determines the wildtype rate of adaptation *v* (*μ, N*) as a function of *μ*(*s*) and *N* (39, 40, *46, 103*).

*When μ*_*m*_ (*s*) ≠ *μ*(*s*), the conditional fixation probability in Eq. (S33) will differ from that of a first-order mutation because there are different mutation spectra in the mean fitness and mutation terms. This is because the modifier lineage primarily competes against the wildtype population [whose mean fitness is controlled by *μ* (*s*)], but acquires additional mutations from its own distribution, *μ*_*m*_(*s*). We note, however, that since the competition with the wildtype population is completely mediated by *v*(*μ, N*) (a monotonic function of *N*), the conditional fixation probability of the modifier lineage can always be mapped to the conditional fixation probability of a first-order mutation in a population that is fixed for the modifier allele, but with a different population size *N*^*^. In other words,

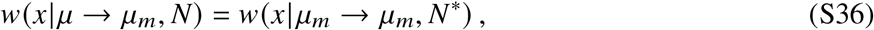

where *N*^*^ is defined by

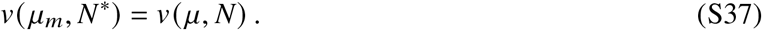

This implies that the space of solutions for *w*(*x*|*μ* → *μ*_*m*_) will have the same general form as the “first-order” *w*(*x*) function that has been studied in previous work (39, 40, 46, 102, 103). It also implies that the dynamics of non-extinction in Eq. (S32), which were previously described in Ref. (40), will be qualitatively similar as well. However, we will see below that actually using this result for evolvability modifiers will often require solutions to Eq. (S33) that go beyond the parameter regimes that have been examined in these earlier studies. We have therefore developed a new approach for deriving approximate analytical solutions for *w*(*x*|*μ* → *μ*_*m*_) that will apply across this broader range of parameters. We outline the general approach in SI Section 3.3 below, and apply it to different classes of distributions of fitness effects in SI Sections 4 and 5.

For notational convenience, we will suppress the explicit dependence on *μ*(*s*) and *μ*_*m*_(*s*) in the following sections, writing *w* (*x*) ≡ *w* (*x* | *μ* → *μ*) for the conditional fixation probability of a first-order mutation and *w*_*m*_ ≡ *w* (*x* | *μ* → *μ*_*m*_) for the conditional fixation probability of the modifier lineage. We will also let *v* ≡ *v* (*μ*) denote the rate of adaptation in the wildtype population and *v*_*m*_ ≡ *v*(*μ*_*m*_) denote the long-term rate of adaptation that is achieved if the modifier takes over.

### 3.3 Asymptotic solution in the “sharp shoulder” regime

There are many parameter regimes of clonal interference that one can consider (40, 103). Here, we will primarily focus on a regime that is relevant for a broad range of naturally and experimentally evolving populations, which roughly corresponds to the case where the fitness benefits of a typical “driver mutation” (*s*_*b*_) are much larger than the total rate at which they occur (*U*_*b*_) (40, 46). We will define these conditions more precisely below.

#### Relative fitness distribution

Previous work has shown that in our regime of interest, the solution to Eq. (S31) can be approximated by a truncated Gaussian distribution,

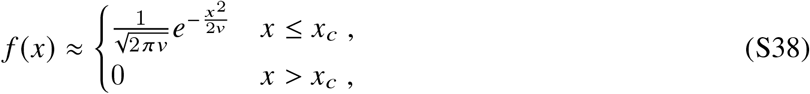

where *x*_*c*_ denotes the location of the fittest individuals that are likely to exist within the population (39, 40, 46). This solution is valid when 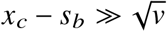 and 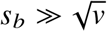, which constitutes the formal definition of the regime that we consider in this work. The first condition 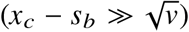 implies that the parents of the most fit individuals are substantially more fit than the majority of the population (i.e. clonal interference is common).

The second condition 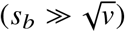 implies that the majority of the population is concentrated near the mean fitness. This is also known as the “moderate speeds” regime (40), since the second condition can be rewritten as 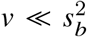. We will assume that these conditions hold for the wildtype population in which the modifier arises.

#### Lineage fixation probability

Previous work (39, 46) has shown that in our parameter regime of interest, the solutions to Eq. (S33) can be decomposed into a high-fitness region, where the mutation term is sub-dominant:

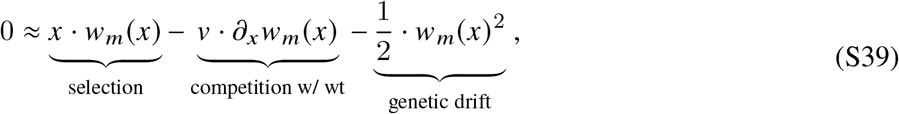

and a linearized region, where the mutation term is important but the drift term can be neglected:

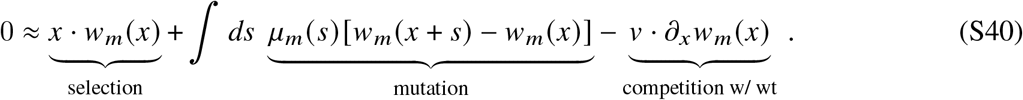

There is also a narrow region in the middle where both approximations are valid, so that asymptotic matching can be used to obtain a full global solution. This asymptotic decomposition will continue to be valid for our modifier case as well.

In the high fitness region, the solution to Eq. (S39) is given by the *shoulder solution*,

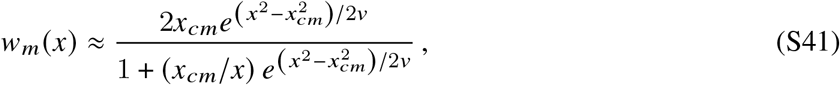

where *x*_*cm*_ is a constant of integration that will be determined self-consistently below. When 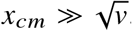, this shoulder solution develops a narrow boundary layer around *x* = *x*_*cm*_ ± 𝒪 (*v*/*x*_*cm*_), such that the fixation probability can be well-approximated by the piecewise form,

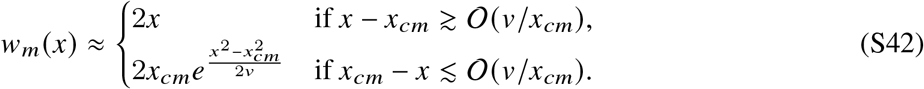

This piecewise function has a simple interpretation in terms of the dominant balances in Eq. (S39) (39). The linear scaling above *x*_*cm*_ emerges from a balance between the selection and genetic drift terms; this implies that lineages with *x* ≳ *x*_*cm*_ will fix provided that they survive genetic drift. Conversely, the exponential scaling at lower *x* emerges from a balance between the selection and mean fitness terms; this implies that lineages with *x* ≲ *x*_*cm*_ are strongly influenced by competition with the surrounding population (i.e. clonal interference). In the special case where *μ*_*m*_(*s*) = *μ*(*s*), previous work has shown that *x*_*cm*_ approximately coincides with the nose of the fitness distribution in Eq. (S38) (39, 40, 46). This makes some intuitive sense: lineages with relative fitness ≳ *x*_*c*_ do not experience clonal interference, and will fix if they survive genetic drift; this suggests that the steady-state fitness distribution should not contain such individuals, since they should already have fixed. We will therefore also refer to *x*_*c*_ as the *interference threshold* for the wildtype population. When *μ*_*m*_(*s*) ≠ *μ*(*s*) the interference threshold of the modifier (*x*_*cm*_) will generally differ from the location of the nose (*x*_*c*_) — this will be critically important for our analysis in SI Sections 4 and 5 below.

The mutation terms must eventually become important for smaller values of *x*, since the shoulder solution in Eq. (S42) starts to increase for *x <* 0. To avoid this unphysical behavior, and ensure *w*_*m*_(*x*) decreases as *x* → ∞, the shoulder solution must eventually map on to the correct branch of the solution to Eq. (S33). Previous work has used saddle point methods to identify the relevant solutions to Eq. (S40) (40, 103), while other studies have utilized thresholding approximations that set *w*_*m*_(*x*) ≈ 0 below a critical fitness value (39, 46). Here we take a slightly different approach, by recasting Eq. (S33) as an integral equation.

Multiplying both sides of Eq. (S33) by 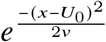 and integrating from −∞ to *x*, we can rewrite Eq. (S33) in the recursive form,

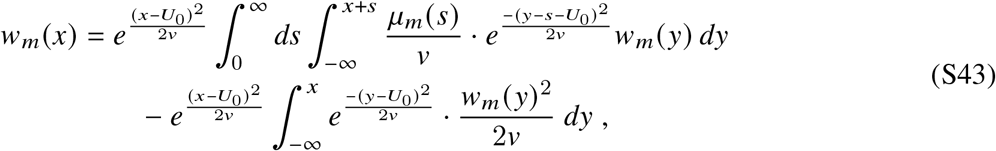

where *U*_0_ ≡ ∫ *μ*_*m*_(*s*)*ds* is the total mutation rate. Since *w*_*m*_(*x*) rapidly declines below the interference threshold, the contributions from the second term will become negligible when *x* ≲ *x*_*cm*_ − 𝒪 (*v*/*x*_*cm*_), and Eq. (S43) will reduce to the simpler form

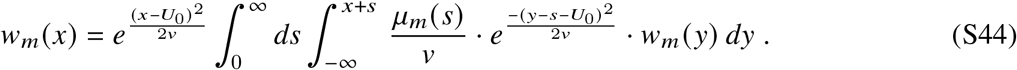

This recursive formula has a simple intuitive interpretation illustrated by the schematic in Fig. 1. A modifier lineage founded at a relative fitness *x < x*_*cm*_ will start as a single clone, whose average size will evolve as

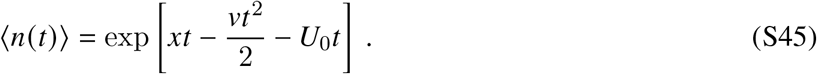

These growth dynamics account for the steady increase in the wildtype mean fitness, as well as the outflow of individuals that acquire further mutations as the clone is growing. The total production rate of new mutations is ⟨*n*(*t*)⟩ · *μ*(*s*)*ds*, and each of these events will found a new modifier lineage with relative fitness *x* + *s* − *vt*; the original lineage will fix if one of these descendant lineages is ultimately successful, yielding the recursive formula,

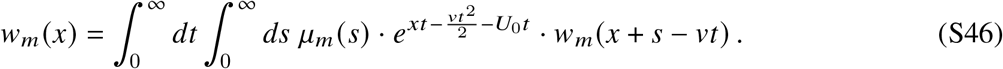

The linearity of this expression implies that successful clones are highly unlikely to give rise to multiple successful mutants, which is consistent with the assumption that clonal interference is very strong (*w*_*m*_ (*x*) ≪ 2*x*) when *x < x*_*cm*_. Equation (S44) can be recovered from Eq. (S46) by changing variables from *t* to *y* ≡ *x* − *v* · *t* + *s*, which represents the time-dependent landing fitness of the clone’s mutant offspring.

Note that while the recursion in Eq. (S44) is also a solution to the linearized version of Eq. (S33), our present derivation shows that the interpretation is slightly different here. In particular, since the upper limit of the *y*-integral in Eq. (S44) is larger than *x*, the mutation term can in principle depend on the behavior of *w*_*m*_(*x*) for *x > x*_*cm*_, where the effects of the nonlinear *w*_*m*_(*x*) ^2^ term start to become important. We can account for this non-locality using the shoulder solution in Eq. (S42), by rewriting Eq. (S44) as a piecewise integral,

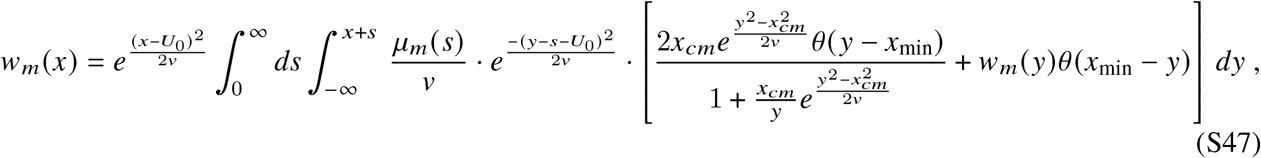

where *θ*(·) is the unit step function, and *x*_min_ represents the point a which the shoulder solution starts to break down. When *x*_min_ ≲ *x*_*cm*_ − 𝒪 (*v x*_*cm*_), we can use this expression to determine *x*_*cm*_, by noting that Eq. (S47) must also match the shoulder solution in Eq. (S42) in the overlap region where both approximations our valid [*x*_min_ ≲ *x* ≲ *x*_*c*_ − 𝒪 (*x*_*c*_)].

While this integral formulation is similar to the transform methods (40, 103) that have previously been used to analyze *w*_*m*_(*x*), it offers practical advantages that will become important for our analysis of modifier mutations below. In particular, we will see that in our parameter regime of interest, we can use Eq. (S47) to analytically extend the shoulder solution to progressively lower fitness values. These analytical approximations will be critically important for treating modifier mutations with large direct costs (Fig. 3).

In the following sections, we use this framework to derive explicit solutions for *w*_*m*_(*x*) different choices of *μ*_*m*_(*s*). We begin by considering a simple model, where the driver mutations all share the same characteristic fitness benefit (SI Section 4). This will allow us to verify the self-consistency conditions assumed above, and to obtain explicit predictions for the overall fixation probabilities of different modifier mutations. We then extend these calculations to continuous distributions of fitness effects in SI Section 5, and show that more general distributions can often be understood using the simple model in SI Section 4.

## 4 Solution for a simple model of the distribution of fitness effects

To make analytical progress, we first consider a simple scenario in which the wildtype population produces mutations with a single fitness benefit *s*_*b*_ at a total rate *U*_*b*_. The distribution of fitness effects can then be written as a point mass,

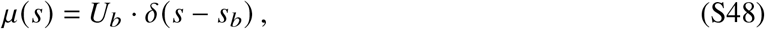

where *δ*(*z*) is the Dirac delta function. Previous work has shown that in our parameter regime of interest, the rate of adaptation (*v*) and interference threshold (*x*_*c*_) follow the approximate scaling,

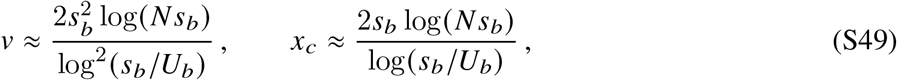

which are valid in the limit that *x*_*c*_ ≫ *s*_*b*_ (39, 40, 47). In terms of these parameters, the conditions we assumed for our solution in SI Section 3.3 (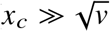 and 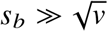) become

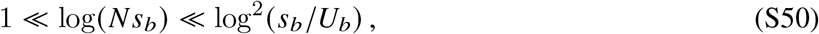

which require that *N s*_*b*_ ≫ 1 and *s*_*b*_ ≫ *U*_*b*_.

For distributions of fitness effects in the same class as Eq. (S48), the most general evolvability modifier is one that produces mutations with a different benefit 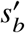 and different rate 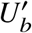, so that

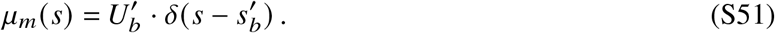

In the analysis below, we will assume that 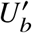 and 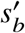 are chosen such that 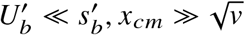, and 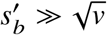. In this regime, the outflow of mutations can be neglected in Eq. (S47). We will also assume that the fold change in the mutation rate may be 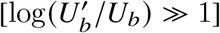, but the fold change in the selection strength will always be comparatively modest 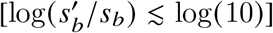. This will be sufficient to derive all of the results in the main text.

Substituting Eq. (S51) into Eq. (S47) yields an integral relation for *w*_*m*_(*x*),

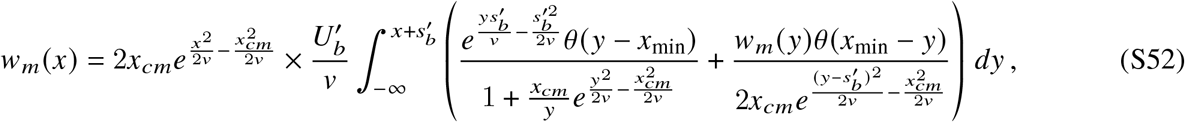

where we have neglected the outflow due to new mutations since 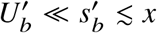. The interference threshold is determined by matching Eq. (S52) to the shoulder solution when *x* ≲ *x*_*cm*_ − 𝒪 (*v*/*x*_*cm*_), which yields a related integral,

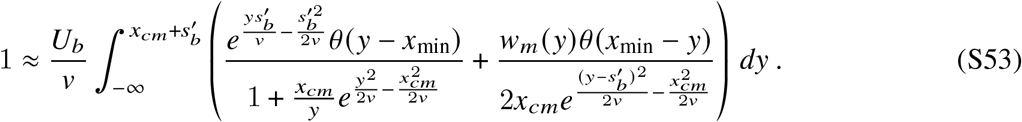

The integrals in Eqs. (S52) and (S53) will sensitively depend on the relative values of *x*_*cm*_ and 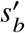. When 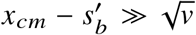, the integrand in Eq. (S53) will be maximized for relative fitnesses near *x*_*cm*_. We will refer to this limit as the *multiple mutations regime*, since it implies that successful lineages must always possess anomalously high relative fitnesses 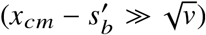. In the opposite extreme 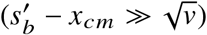, the integrand in Eq. (S53) will be maximized for relative fitnesses near 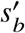, where the shoulder solution has already transitioned to the Haldane limit [*w*(*x*) ≈ 2*x*]. We will refer to this limit as the *quasi-sweep regime*, since it implies that lineages will be guaranteed to fix if they produce a single mutation that survives genetic drift. Our assumptions imply that the wildtype population will always fall in the multiple mutations regime, but the modifier lineage may differ depending on the relative values of 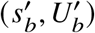 and 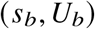. We consider each case separately below.

### 4.1 Multiple mutations regime

#### 4.1.1 Location of the interference threshold

When 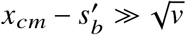, the integral in Eq. (S52) will be dominated by fitnesses close to *x*_*cm*_. In particular, the contributions from *x < x*_*cm*_ will be dominated by fitnesses within 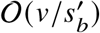 of *x*_*cm*_, while the contributions from will be dominated by fitness within 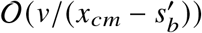 of *x*_*cm*_. The dominant contribution will therefore depend on the relative magnitudes of 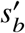 and 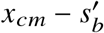. When 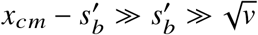, the dominant contribution will come from the exponential region of Eq. (S42), so that the auxilliary condition becomes

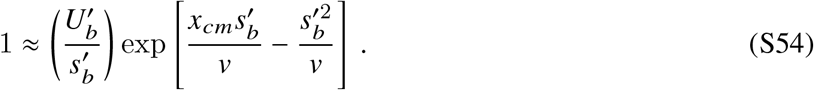

Note that this derivation implicitly assumes that the shoulder solution extends at least 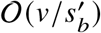 below *x*_*cm*_; we will validate this assumption below. In the opposite case where 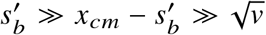, the dominant contribution to the integral in Eq. (S53) will come from the linear region of Eq. (S42), so that the auxilliary condition becomes

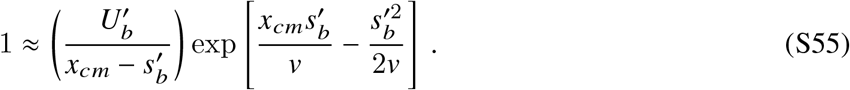

For convenience, we will summarize these two equations with the common expression,

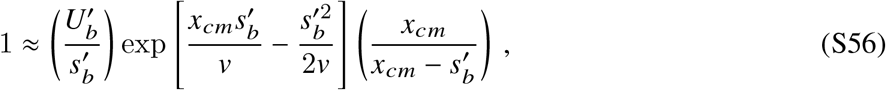

which reduces to the correct scaling in the corresponding limits. Since we have assumed that the wildtype population always lies in the multiple mutations regime 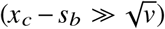, the background interference threshold must satisfy an analogous condition,

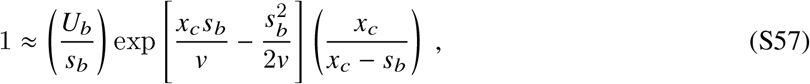

which matches the expression derived in Ref. (46). These expressions allow us to solve for *x*_*cm*_ and *x*_*c*_ as a function of 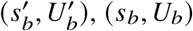, and *v*. In particular, by dividing Eq. (S56) by Eq. (S57), we can solve for *x*_*cm*_ as a function of the fold changes in *U*_*b*_ and *s*_*b*_:

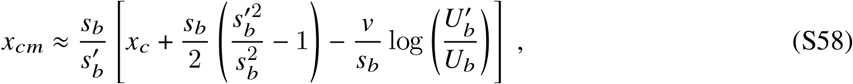

where we have neglected logarithmic corrections in *x*_*cm*_/*x*_*c*_ and 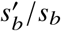. This shows that the 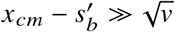 assumption will be valid provided that

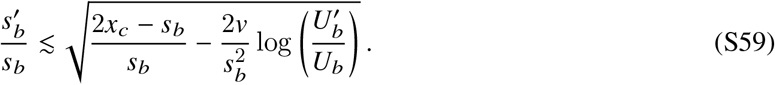

In particular, Eq. (S59) shows that the multiple mutations regime will apply for pure mutation rate modifiers as large as 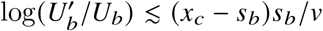, while pure selection strength modifiers require the more stringent condition, 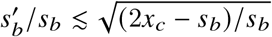. Violations of these conditions are considered in SI Section 4.2 below.

#### 4.1.2 Extending the shoulder solution to lower fitness values

With the location of *x*_*cm*_ fixed by Eq. (S56), we will now use the integral recursion in Eq. (S52) to extend the shoulder solution for *w*_*m*_(*x*) to progressively lower fitness values. When 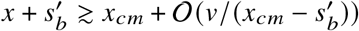, the dominant contribution to Eq. (S53) will also be contained within the region of integration in Eq. (S52).

We can therefore substitute Eq. (S53) to obtain

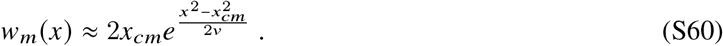

This derivation makes it clear that the shoulder solution will continue to be valid for fitnesses as low as

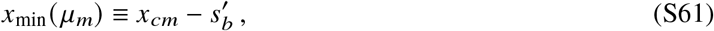

which will serve as our definition of *x*_min_ in Eqs. (S53) and (S52) above. This validates our assumption that the mutation term in Eq. (S33) is negligible for 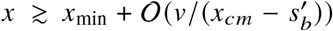. Moreover, since 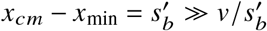, it also validates our assumption that the location of *x*_*cm*_ in Eq. (S56) is completely determined by regions where the shoulder solution is valid (*x* > *x*_min_). For fitnesses below *x*_min_, the finite upper limit in Eq. (S52) will start to become important. However, as long as 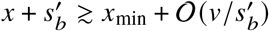, the integral will continue to be dominated by regions where the shoulder solution is valid. When *x* ≲ *x*_min_ − 𝒪 (*v/x*_*cm*_), the upper limit of integration will fall within the exponential region of the shoulder solution, yielding

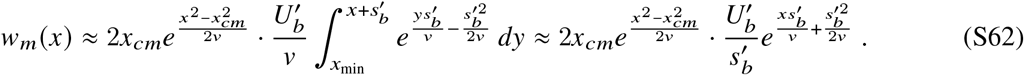

After dividing this expression by the auxiliary condition for *x*_*cm*_ in Eq. (S56), we obtain

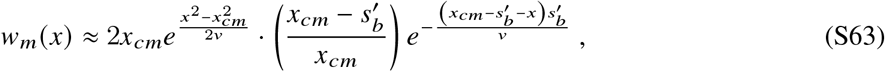

which will be valid for relative fitnesses in the range

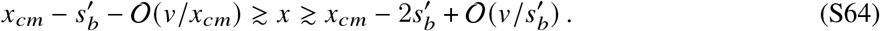

By comparing Eq. (S60) and Eq. (S63), we can see that *w*_*m*_(*x*) declines by a factor of 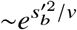 across this region. Since we have assumed that 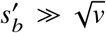, this provides a natural justification for the thresholding approximation employed in Ref. (46), which assumed that *w x* was negligible for fitnesses below *x*_min_. Equation (S63) constitutes a more quantitative version of this approximation, which will be useful for the analysis below.

We can continue these calculations to recursively extend *w*_*m*_(*x*) to progressively lower fitness values. When 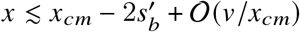, the shoulder solution will not contribute at all to Eq. (S52), but the region in Eq. (S63) will dominate, yielding

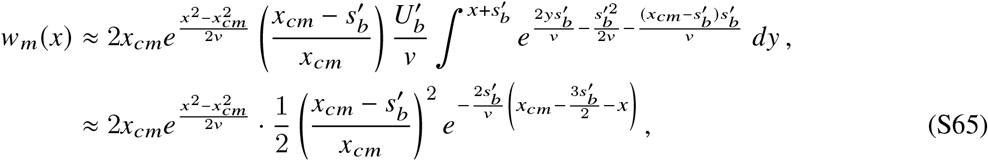

for 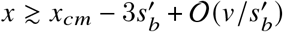. More generally, for relative fitnesses in the range

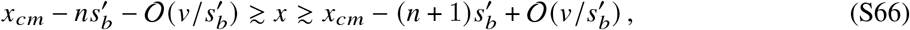

one can continue this argument to show that

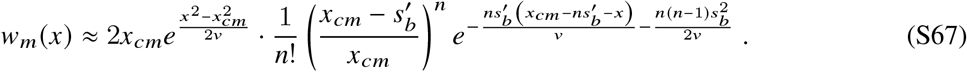

This shows that each additional “step” reduces the original shoulder solution by factor of 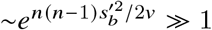. For fitness values that are many multiples of 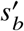 below *x*_*cm*_, we can substitute 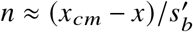 to show that the leading order contribution to *w*_*m*_(*x*) scales as

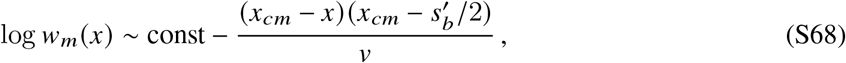

which obeys the required boundary condition that *w*_*m*_(*x*) → 0 as *x* → −∞. Thus, Eq. (S67) provides an asymptotic solution for *w*_*m*_(*x*) that is valid across the full range of relative fitnesses below *x*_*cm*_.

#### 4.1.3 First-order mutations and the rate of adaptation

When *μ*_*m*_(*s*) → *μ*(*s*), our solution for *w*_*m*_(*x*) can be used to derive the fixation probabilities of first-order mutations, which have been studied in previous work (39, 40, 46). We reproduce these results here for completeness, using the new expressions for *w*(*x*) that we have derived above. This will allow us to make comparisons to the second-order selection pressures analyzed in SI Section 4.1.4 below. (Readers who are familiar with this material may skip directly to SI Section 4.1.4.)

For a completely neutral mutation (*s*_*m*_ = 0), we can substitute our solution for *f*(*x*) in Eq. (S38) into the self-consistency condition in Eq. (S35) to rewrite it in the convenient form:

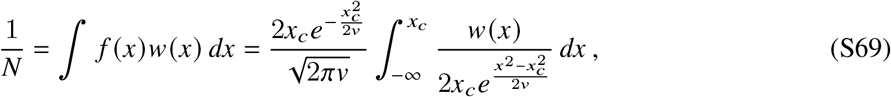

which depends on the ratio between the true value of *w* (*x*) and the exponential shoulder solution in Eq. (S42). Substituting our solution for *w* (*x*) in Eq. (S67) then yields

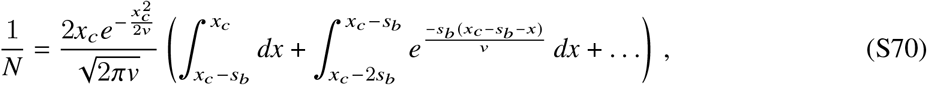

where we have only included the *n* = 0 and *n* = 1 regions from Eq. (S67); all of the other terms are smaller by additional factors of 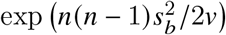, and will therefore provide a negligible contribution when 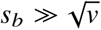. Note that the first term in Eq. (S70) contributes equally over the whole range of *x* ∈ (*x*_*c*_ − *s*_*b*_, *s*_*b*_), while the second term is dominated by fitnesses within 𝒪(*v*/*s*_*b*_) of *x*_*c*_ − *s*_*b*_. Since *s*_*b*_ ≫ *v*/*s*_*b*_, the first term dominates, yielding

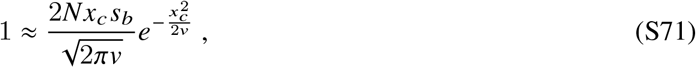

which matches the condition previously derived in Ref. (46). When combined with the auxiliary condition for *x*_*c*_ in Eq. (S57), this allows us to solve for *v* and *x*_*c*_ as a function of the underlying parameters *N, s*_*b*_, and *U*_*b*_. An iterative solution of Eqs. (S71) and (S57) assuming that 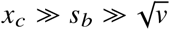 yields the asymptotic expressions for *v* and *x*_*c*_ in Eq. (S49). The fixation probabilities of non-neutral mutations can be calculated using a similar procedure. We first use the self-consistency condition in Eq. (S71) to rewrite the scaled fixation probability in a similar form as Eq. (S69),

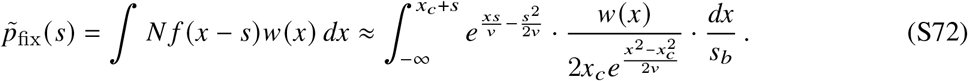

This differs from the previous integral in Eq. (S69) by the presence of the exponential factor *e*^*xs*/*v*^ and the shifting of the upper limit of integration from *x*_*c*_ to *x*_*c*_ + *s*. The behavior of this integral will therefore depend on the sign of *s*.

##### Beneficial mutations

When *s* > 0, the *e*^*xs*/*v*^ term will enhance the contributions from higher fitness values, so the *n* ≥ 1 terms in Eq. (S67) will continue to be negligible. Moreover, since the upper limit of integration is now larger than *x*_*c*_, there will also be a new contribution from the Haldane region of the shoulder solution, so that

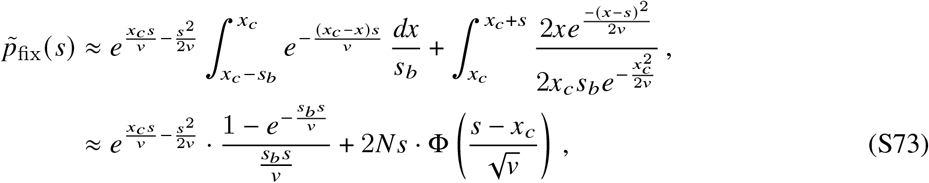

where 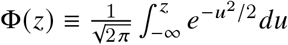 is the Gaussian cumulative distribution function. The leading order scaling is given by

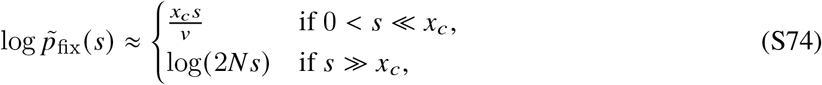

which transitions between a regime of strong clonal interference for *s* ≪ *x*_*c*_ and the Haldane limit when *s* ≫ *x*_*c*_ (39).

The fixation probabilities in Eq. (S74) have a simple heuristic interpretation. To be a successful, a moderately beneficial mutation (*s* ≪ *x*_*c*_) will typically arise on a background near *x* = *x*_*c*_ − *s*. This mutation will found a new lineage which experiences the effects of clonal interference and must generate multiple more fit descendants to take over and fix. Over this fixation time, *T*_*sw*_ = *x*_*c*_/*v*, the beneficial mutation will provide an exponential advantage to its founding lineage. On the other hand, successful mutations that are extremely beneficial (*s* ≫ *x*_*c*_) will typically land above the interference threshold (*x > x*_*c*_ − *s*) and fix if they survive genetic drift.

##### Deleterious mutations

The fixation probabilities of deleterious mutations (*s <* 0) can be calculated using a similar procedure. In this case, however, the *e*^−*x*|*s*|/*v*^ term becomes increasingly large at lower fitness values, and will need to be cut off by one of the *n* ≥ 1 terms in Eq. (S67). To build intuition, let us first consider the case where |*s*| *< s*_*b*_, so that

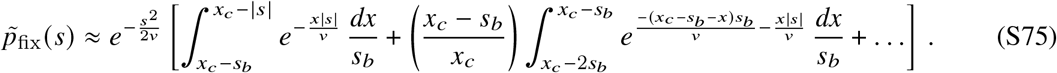

The contribution from the *n* = 0 term is now peaked at its lower limit of integration (*x* ≈ *x*_*c*_ − *s*_*b*_). However, when |*s*| is smaller than *s*_*b*_, the contribution from the *n* = 1 term is still peaked at its upper limit of integration (*x* ≈ *x*_*c*_ − *s*_*b*_), which implies that the total contribution to the fixation probability will also be peaked at *x* ≈ *x*_*c*_ − *s*_*b*_. [The contributions from the terms will be smaller by additional factors of 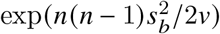, and will therefore be negligible in the limit that 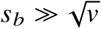.] Evaluating the two integrals then yields

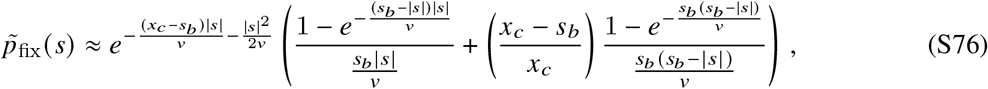

which is valid for fitness costs *s* ∈ (−*s*_*b*_ + 𝒪(*v*/*s*_*b*_), 0). When |*s*| ≪ *s*_*b*_, this matches the fixation probability derived using the thresholding approximation in Ref. (46), but it starts to deviate from Ref. (46) for |*s*| ∼ 𝒪(*s*_*b*_). We expect that the expression in Eq. (S76) will be more accurate in this case, since it better captures the behavior of *w*(*x*) below *x* ≈ *x*_*c*_ − *s*_*b*_.

The fixation probabilities of more strongly deleterious mutations can be calculated in a similar manner. In this case, it will be convenient to write

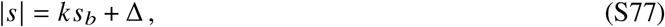

where *k* is the largest integer such that Δ *>* 0. The upper limit of integration at *x*_*c*_ − *s* ≡ *x*_*c*_ − *k s*_*b*_ − Δ implies that the first term with nonzero contribution to *p*_fix_ will be the *n* = *k* term, so that

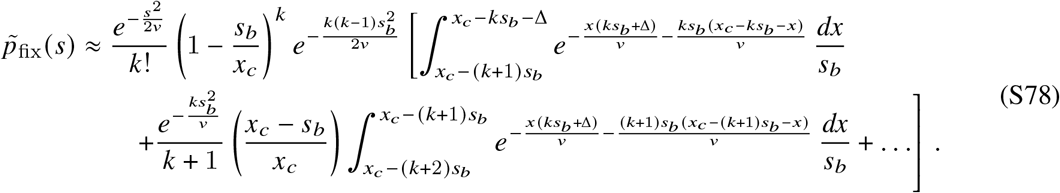

As above, the contribution from the *n* = *k* term is peaked at the lower limit of integration (*x* ≈ *x*_*c*_ − (*k* + 1)*s*_*b*_), while the contribution from the *n* = *k* + 1 term is peaked at its upper limit (*x* ≈ *x*_*c*_ − (*k* + 1)*s*_*b*_). The terms with *n* ≥ *k* + 2 are smaller by exponential factors of 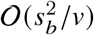, and will therefore provide a negligible contribution to the fixation probability when 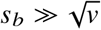. Evaluating the two integrals then yields,

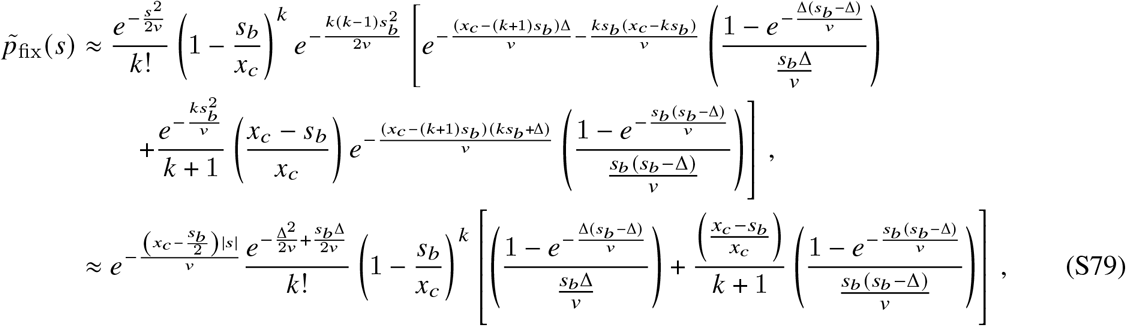

whose leading order behavior simplifies to

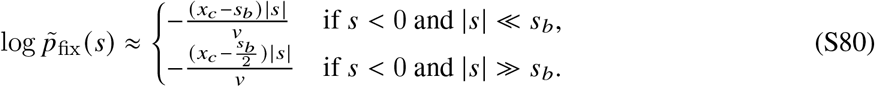

For the purposes of numerical evaluation, it is useful to employ a modified version of Eq. (S79),

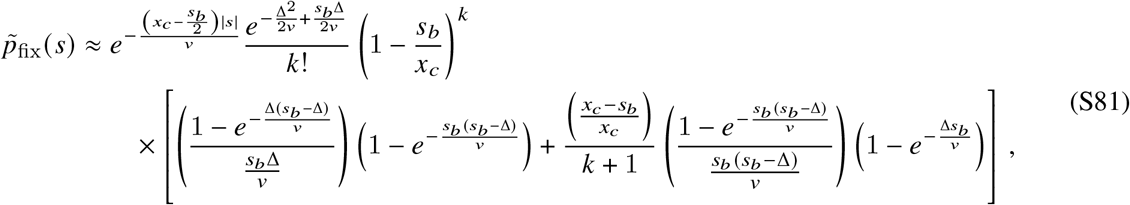

which has the same asymptotic limit when 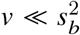, but enforces strict continuity at Δ = *s*_*b*_.

#### 4.1.4 Modifiers without direct costs or benefits

We now are in a position to calculate the fixation probabilities of modifier mutations (*μ*_*m*_ (*s*) ≠ *μ*(*s*)). In the absence of a direct cost or benefit (*s*_*m*_ = 0), this will only be a slight generalization of the neutral fixation probability calculation in SI Section 4.1.3. In the case of a modifier mutation, Eq. (S69) becomes

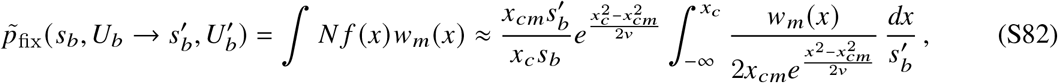

where the primary difference is the presence of the prefactor term, and the fact that the upper limit of integration is now equal to *x*_*c*_ ≠ *x*_*cm*_. The behavior of this integral will sensitively depend on the relative magnitudes of *x*_*cm*_ and *x*_*c*_.

##### Positively selected modifiers

If the modifier interference threshold is less than that of the wildtype (*x*_*cm*_ *< x*_*c*_), then the integral in Eq. (S82) will contain both the *n* ≥ 0 terms as well as a portion of the Haldane region,

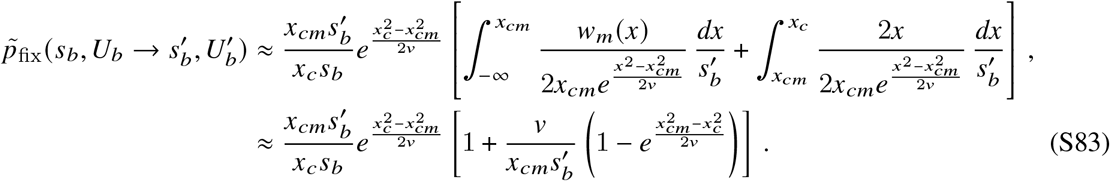

The second term is negligible in our regime of interest where 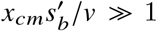, so the fixation probability reduces to

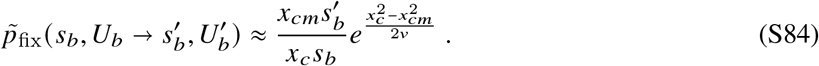

Since *x*_*cm*_ *< x*_*c*_, Eq. (S84) implies that 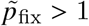 (i.e., the modifier is favored by natural selection).

##### Negatively selected modifiers

In the opposite case, where the interference threshold of the modifier is greater than that of the wildtype (*x*_*cm*_ *> x*_*c*_), the upper limit of the integral in Eq. (S82) will occur somewhere in the interference region of *w*_*m*_(*x*). The precise behavior will depend on how far *x*_*c*_ extends into this region. If 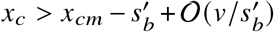, then the dominant contribution will still come from the *n* = 0 region of Eq. (S67), so that

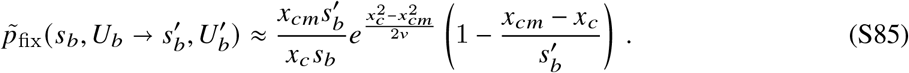

Since *x*_*cm*_ *> x*_*c*_, this implies that *p*_fix/_*p*_0_ *<* 1 (i.e., the modifier is disfavored by natural selection). More generally, if

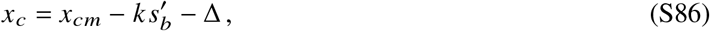

for some 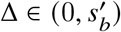, then the primary contribution will come from the *n* = *k* region of Eq. (S67). This yields

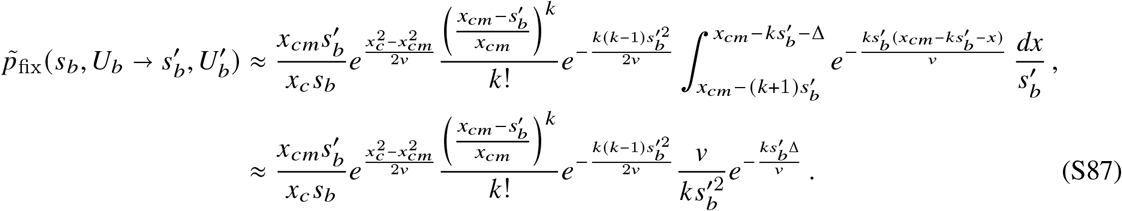

##### ***Predictions for specific values of*** 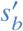 ***and*** 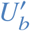

We can express the fixation probability of a modifier in terms of the underlying parameters, 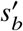, 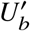, and *N* by substituting Eq. (S58) into Eq. (S84) to obtain,

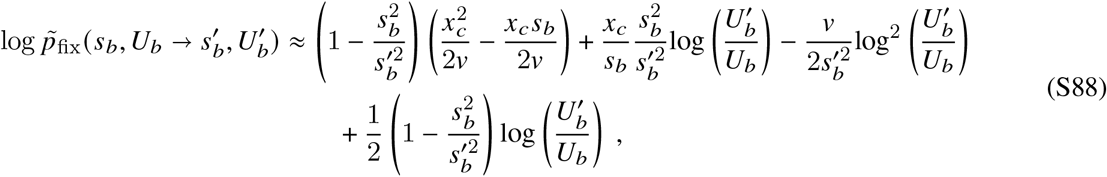

where we have assumed that 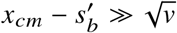. In the case of a selection strength modifier 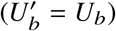, the leading order contributions simplify to

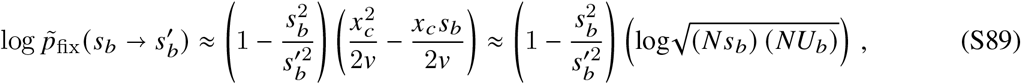

where we have substituted 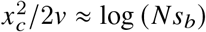 and *x*_*c*_*s*_*b*_/*v* ≈ log (*s*_*b*_/*U*_*b*_) on the right hand side.

The fixation probability of a selection strength modifier has a simple heuristic explanation. For small changes in the selection coefficient 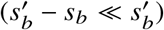, successful mutations arise in the high-fitness “nose” of *f* (*x*) (*x* ≈ *x*_*c*_ ≫ *s*_*b*_) and must acquire ∼*x*_*c*_/*s*_*b*_ additional mutations before they reach 𝒪(1) frequencies. In each of these steps, a selection-strength modifier produces 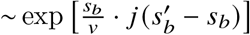 more mutations than a wildtype individual with the same fitness, leading to the exponential scaling observed in Eq. (3).

Similarly, the fixation probability of a mutation rate modifier is given by setting 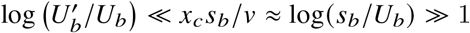 in Eq. (S88), which yields the leading order contribution

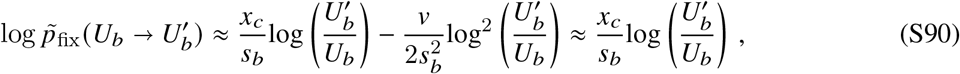

where we have assumed that *x*_*c*_/*s*_*b*_ ≈ 2log (*N s*_*b*_)/log(*s*_*b*_/*U*_*b*_) and 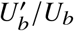. This matches the result previously derived in Ref. (24). The heuristic picture is similar to the one described in that work: a mutation rate modifier must also arise in the high-fitness nose of the fitness distribution (*x* ≈ *x*_*c*_) and accrue *x*_*c*_/*s*_*b*_ additional mutations to fix. In each of these *x*_*c*_/*s*_*b*_ steps before a coalescent event, the mutation rate modifier produces 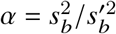 more offspring, leading to the observed scaling in Eq. (S90).

More generally, we see that the fixation probability of a joint selection strength and mutation rate modifier can be expressed in terms of the fixation probabilities of these stand-alone modifiers,

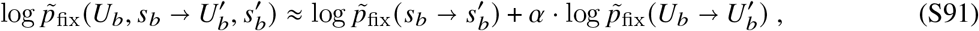

where 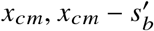. The presence of this factor *α* indicates that a change to the selection strength of mutations can modulate the effect of a linked mutation rate change. The effect of this interplay can be extremely important to determining the fate a modifier mutation, causing modifiers that would be disfavored by natural selection without this modulation to be favored. Interestingly, the modulation effect only works in this direction: modifiers that would be favored without the effect cannot actually be deleterious.

#### 4.1.5 Modifiers with direct costs or benefits

Finally, we can extend these calculations to modifiers with direct costs or benefits by combining our results from SI Sections 4.1.3 and 4.1.4. Generalizing Eqs. (S72) or (S82) to this case yields

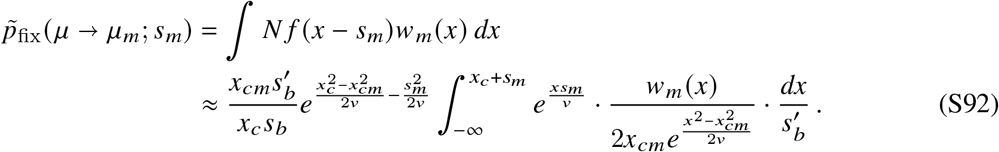

Our analysis above suggests that the two key considerations will be (i) the sign of *s*_*m*_, and (ii) where *x*_*c*_ + *s*_*m*_ falls relative to the internal scales of *w*_*m*_(*x*) (e.g. 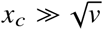, etc.).

##### Modifiers with direct fitness benefits

For a modifier with a direct fitness benefit (*s*_*m*_ *>* 0), the exponential term will once again amplify the contributions from higher fitness values. The behavior of Eq. (S92) will therefore strongly depend on the location of the upper limit of integration. If *s*_*m*_ − (*x*_*cm*_ − *x*_*c*_) *>* 𝒪 (*v*/*x*_*cm*_), there will be a new contribution from the Haldane region of *w*_*m*_(*x*), so that

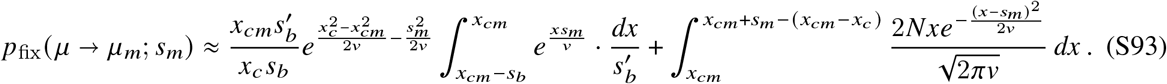

Since 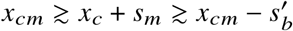, the upper limit from the Haldane region can be taken to infinity, so that Eq. (S93) reduces to

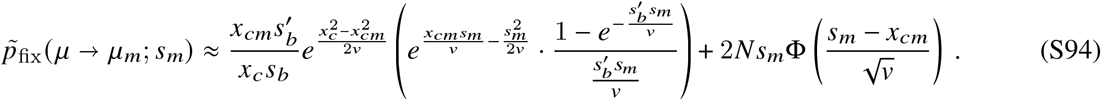

In the opposite case, where *s*_*m*_ ≲ *x*_*cm*_ −*x*_*c*_ − 𝒪 *(v* /*x*_*cm*_), the upper limit of integration in Eq. (S92) will fall below the interference threshold of *w*_*m*_ *x*. When *s*_*m*_ *>* 0, this will only occur when *x*_*cm*_ *> x*_*c*_ (i.e., when the modifier would be disfavored on its own). Similar to the pure modifier case in Eqs. (S85) and (S87), the behavior of Eq. (S92) will depend on where *x*_*c*_ + *s*_*m*_ falls relative to the internal scales of *w*_*m*_(*x*). If 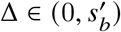, then the dominant contribution will come from the *n* = 0 region of Eq. (S67), so that

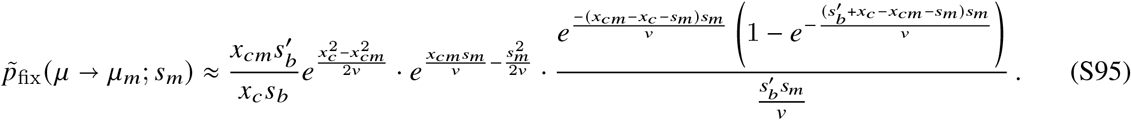

More generally, if we have

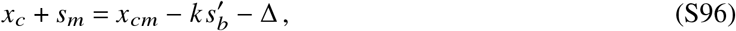

for some 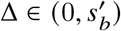, then the primary contribution will come from the *n* = *k* region of Eq. (S67). This yields

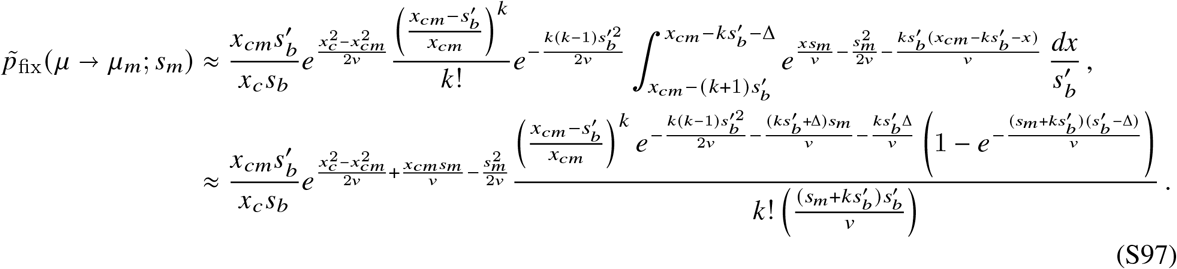

##### Modifiers with direct fitness costs

For a modifier with a direct fitness cost (*s*_*m*_ *<* 0), the exponential term will once again amplify contributions from lower fitness values, and must eventually be cut off by the *n* = *k* + 1 term in Eq. (S67), where *k* is defined by

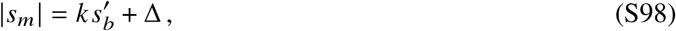

for some 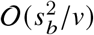. The major difference from the first-order selection scenario in Eq. (S81) is that the upper limit of integration includes an additional term, *x*_*c*_ − *x*_*cm*_. If *x*_*cm*_ *< x*_*c*_ (i.e. a positively selected modifier), then the upper limit of integration is larger than in Eq. (S81). This can in principle lead to contributions from the *n < k* terms in Eq. (S67). However, the *n < k* terms will all be smaller by exponential factors of 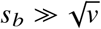 so they will provide a negligible contribution when 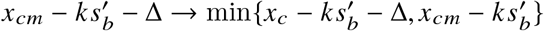. The larger upper limit of integration will therefore only alter the *n* = *k* integral in Eq. (S78), shifting it from 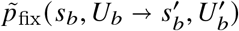. This implies that the modifier fixation probability will be given by a slight generalization of Eq. (S81),

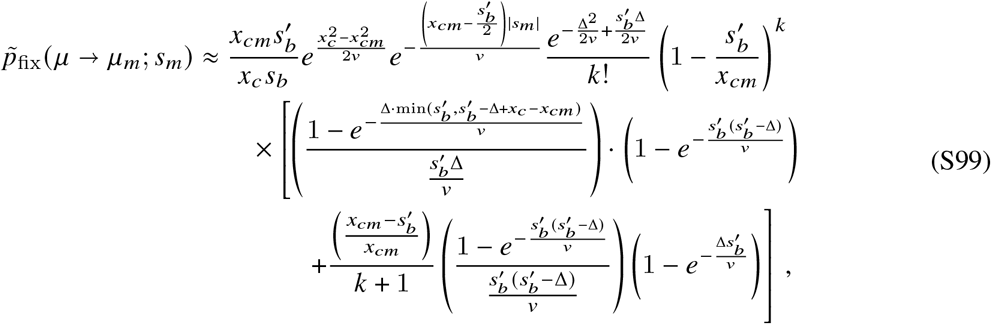

which will be valid for *x*_*cm*_ *< x*_*c*_. The opposite scenario (*x*_*cm*_ *> x*_*c*_) corresponds to a negatively selected modifier that has an additional a direct cost. We will not consider such mutations here, as they will have a negligible chance of fixing.

##### Leading order scaling

Combining these expressions, we see that the leading-order solution for the fixation probability of a modifier with a direct cost or benefit can be summarized as,

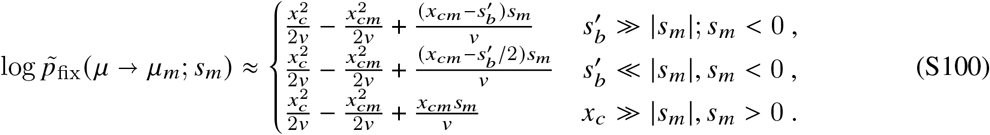

Interestingly, we see that this fixation probability decomposes into contributions from first- and second-order selection,

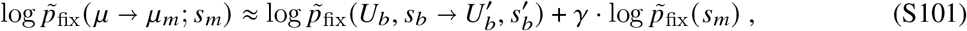

where 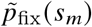 is the fixation probability of a modifier without direct costs or benefits, 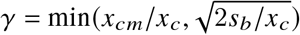 is the fixation probability of a first order mutation, and the weighting factor *γ* satisfies 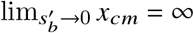. The presence of this factor *γ* demonstrates that second order selection modulates the effects of first order selection; the fixation probability of a deleterious or beneficial mutation arising in the background population scales with *x*_*c*_ /*v*, while the contribution of a direct cost or benefit scales with *x*_*cm*_ /*v*. This means that a modifier without direct costs or benefits that is favored by natural selection (*x*_*cm*_ *< x*_*c*_) suppresses the contribution of a direct cost, while a modifier without direct costs or benefits that is disfavored by natural selection (*x*_*cm*_ *> x*_*c*_) amplifies the contribution of a direct benefit. This effect can be extremely large, affecting the contribution of a direct cost or benefit and overall favorability of a modifier mutation by orders of magnitude.

For sufficiently large direct benefits (| *s*_*m*_| ≫ *x*_*c*_), we expect that even the most deleterious modifiers must fix deterministically upon surviving genetic drift, irrespective of the background on which they arise (*p*_fix_ /*p*_0_ ≈2*N s*_*m*_). This expectation follows from the fact that a modifier with a large direct benefit will jump far out ahead of the ‘traveling wave’ and take over the population long before the background population is able to ‘catch-up’. This is true even in the extreme case of a dead-end modifier, which would drive the rate of adaptation to zero upon fixing. Our current theoretical approach, however, does not enable us to predict the fixation probability of a modifier with direct benefits *s*_*m*_ ∼ 𝒪 (*x*_*c*_). In particular, to derive Eq. (S33) we assumed that the fate of a modifier mutation is determined at small frequency while competing against the mean fitness set by the rate of adaptation of the background population. However, if the rate of adaptation does not change, our theory predicts the background population will always “catch up” and pass a deleterious modifier in fitness, no mater how large the direct benefit. This pathology can be seen in Eq. (S56), where 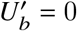.

This limitation prevents us from understanding a key aspect of the trade-off between short-term fitness and long-term evolvability. How large a direct benefit is sufficient to drive a deleterious modifier to be favored? In the next section, we focus on the extreme case of an evolutionary dead-end (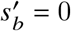or 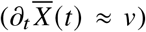), showing that even modest direct benefits can drive these modifiers to be favored by selection.

#### 4.1.6 Fixation of a dead-end modifier

When a dead-end modifier arises with an extremely large direct benefit (| *s*_*m*_| ≫ *x*_*c*_), it will sweep long before the background population is able to catch-up in fitness. On the other hand, if the direct benefit is not sufficiently large, the background population will surpass the modifier lineage in fitness. We can understand this cross-over by analyzing the dynamics of a modifier lineage arising with landing fitness, *x*, as it transitions from small to large frequency.

A newly established modifier clone with relative fitness *x* will initially start to grow deterministically as

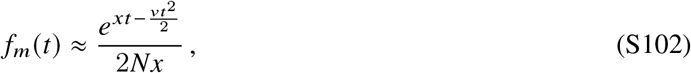

where 1/2*Nx* corresponds to the size of the clone immediately after it establishes. This will be a good approximation as long as the modifier frequency remains small (*f*_*m*_ ≪ 1), so that the mean fitness of the population is still primarily determined by the wildtype 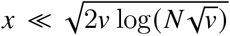. If 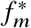), then the modifier clone will remain small throughout its entire lifetime, and the wildtype population will eventually pass it by. However, for larger initial fitnesses, the modifier clone will eventually grow to 𝒪 (1) frequencies, and will start to exert its own effect on the population mean fitness. At this point, the dynamics in Eq. (S102) will start to break down.

The ultimate fate of a dead-end modifier will depend on how much fitness the wildtype population has gained during this time. To understand this process, let *t*^∗^ denote the time required for the modifier clone to reach a reference frequency 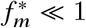. If 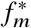, then *t*^∗^ can be calculated from Eq. (S102):

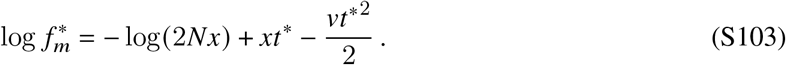

During this time, the wildtype population will have increased in fitness by a total amount Δ*x* = *vt*^∗^, so that the current relative fitness of the modifier is *x* − *vt*^∗^. If the nose of the wildtype population has surpassed the modifier in this time (*x*_*c*_ ≳ *x* − *vt*^∗^), then regardless of how much the modifier grows in the short-term, it will be destined for extinction in the long-term, since it is unable to produce additional mutations.

On the other hand, if *x* − *vt*^∗^ *> x*_*c*_, then it is possible for the modifier to sweep through the rest of population extremely rapidly, and “freeze” the wildtype population in its place. If further adaptation of the wildtype can be neglected, then the modifier will transition from from frequency 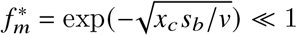 to 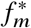 according to the logistic dynamics,

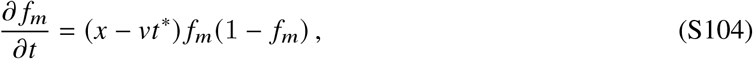

which requires an additional time interval

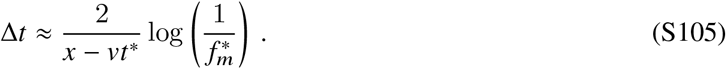

Since we have assumed that *x* − *vt*^∗^ ≥ *x*_*c*_, it is always possible to choose a reference frequency 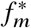 that is≪ 1, but large enough that Eq. (S105) is much smaller than the time required for the wildtype population to acquire one additional mutation (*s*_*b*_/*v*). For example, choosing 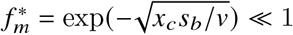 yields

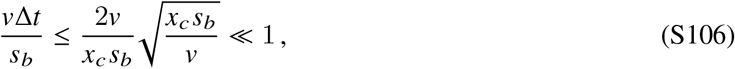

since *x*_*c*_*s*_*b*_ *v* 1. This shows that the wildtype population is effectively frozen in place while the modifier transitions from 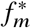 to 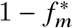. At this point, the mean fitness of the population is dominated by the modifier 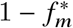. Since we have assumed that *x* − *vt*^∗^ *> x*_*c*_, this implies that even the fittest individuals in the wildtype population now have a negative relative fitness, so their further adaptation will be effectively halted. The modifier clone will therefore continue to sweep through the population, and will fix in a time of order 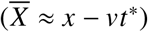.

Based on this reasoning, we conclude that a newly established modifier will fix if its initial fitness exceeds a critical threshold *x*^∗^ defined by

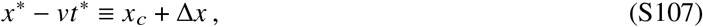

where Δ*x* is a small correction term [≲𝒪 (*s*_*b*_)] that accounts for potential ambiguities in the definition of the nose, as well as the possibility that a leapfrogged nose may produce one additional mutation via stochastic tunneling (91) before it goes extinct. It is also accounts for the fact that a small subclass of individuals in the background population will need to have fitness greater than *x*^∗^ to out-compete the modifier after the mean fitness changes. The size of this subclass can be determined by requiring that the sum of the establishment probabilities of the individuals that have crossed *x*^∗^ be equal to 1,

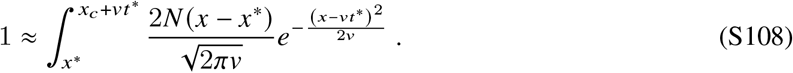

Assuming *v* /*x*_*c*_≪ *x*^∗^ − *vt*^∗^ − *x*_*c*_ and 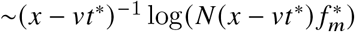, we can evaluate this integral with a Laplace approximation to obtain the self-consistent solution,

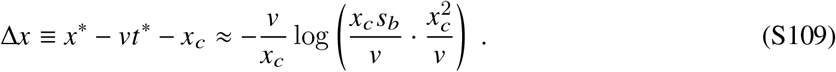

This constitutes a small correction to *x*^∗^ (as expected), but we will see that it provides an important contribution to the fixation probability below. To solve for *x*^∗^, we can substitute Eq. (S109) into Eq. (S103) to obtain the condition,

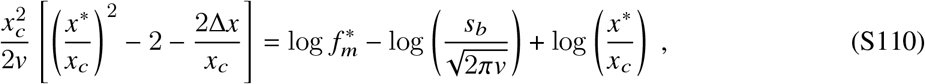

where we have also used Eq. (S71) to substitute for *N*. In our regime of interest where 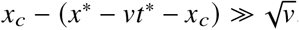 the terms on the left will be asymptotically larger than those on the right, and the dominant balance will be given by the terms in the square brackets, yielding

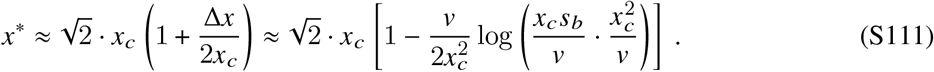

This result implies that a modifier lineage with relative fitness greater than 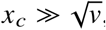 will be guaranteed to fix as long as it survives genetic drift while rare — regardless of its ability to produce additional mutations. This will always be consistent with the wildtype *w x* function from the branching process approximation in SI Section 4.1.3, since *x*^∗^ *> x*_*c*_. However, it suggests that the interference threshold for the modifier lineage must be capped at a maximum value,

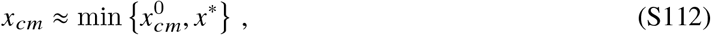

where 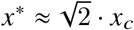 is our original solution for the interference threshold derived in SI Section 4.1.1 above. This constitutes a simple modification of the original branching process model in Eq. (1) that can capture some of these non-linear feedback effects.

In the extreme case of a dead-end modifier (*μ* → ∅), this ansatz leads to a conditional fixation probability of the form

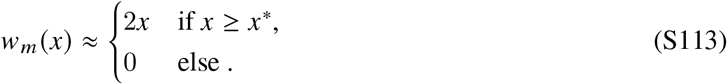

Substituting this result into Eq. (S92) yields a prediction for the total fixation probability of a dead-end modifier with a direct benefit *s*_*m*_:

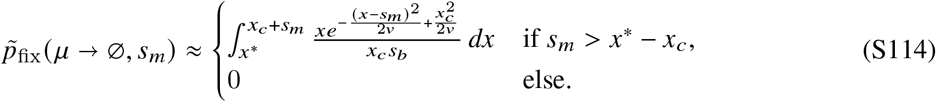

Evaluating the integral in the upper branch yields,

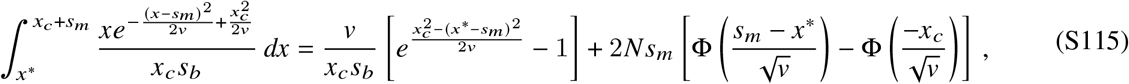

where 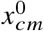 is the Gaussian cumulative distribution function. The leading-order scaling is given by

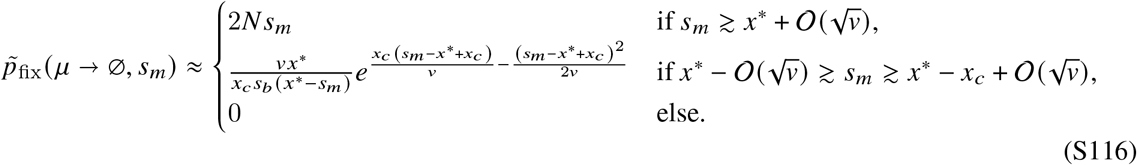

These results suggests that modest direct benefits of size 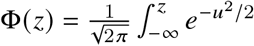 are sufficient to cause an evolutionary dead-end to be favored by natural selection. While this critical size is larger than the advantage of a single mutation (*s*_*b*_), it is smaller than the total fitness variation in the population (*x*_*c*_), and is only weakly dependent on *NU*_*b*_. This contrasts with the traditional linear scaling observed in models with small numbers of competing loci (92), highlighting the unique features that arise in genome-wide models of adaptation.

Together, this analysis shows that the fate of a dead-end modifier is still determined by chance events that occur while the modifier lineage is at low frequency (e.g. its random genetic background and whether it survives genetic drift while rare) even though its ability to take over critically relies on non-linear feedbacks that occur while it is common.

#### 4.1.7 Relation between the fixation probability and the long-term rate of adaptation

It is useful to compare these results with the deterministic modifier theory in SI Section 1, which predicts that modifiers will be favored if they increase the long-term mean fitness of the population. While this result was originally derived for non-adapting populations at mutation-selection balance, a natural extension of this idea to adapting populations might suggest that natural selection would favor modifiers that increase the long-term rate of adaptation (i.e. *v*_*m*_ *> v*).

Our results above show that this simple heuristic clearly breaks down for modifiers with direct costs or benefits (Fig. 3). This discrepancy is most dramatic for the “dead-end” modifiers in SI Section 4.1.6, which can be strongly favored by selection even though they lower the rate of adaptation to zero. Our analysis in SI Section 4.1.6 showed that the origin of this effect could be traced to the early fixation of the modifier lineage before its long-term costs are fully realized. These fixation events are completely neglected by the deterministic theory in SI Section 1: beneficial variants can grow to arbitrarily large frequencies in the short-term, but their long-term fitness gains will always override their initial costs or benefits. This highlights how finite population sizes can be critically important, even when the population size is very large (47).

To understand this relationship in more detail, we can combine our results in Eqs. (S58), (S71), and (S100) to derive an approximate formula connecting *p*_fix_, *v*_*m*_, and *s*_*m*_. In the limit that *x*_*c*_ ≫ *s*_*b*_ and 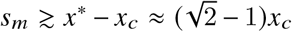, we find that the leading-order contributions satisfy the approximate scaling,

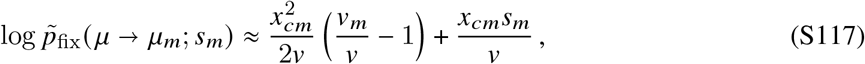

to leading order in the logarithm of 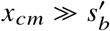. This result shows that in the absence of direct costs or benefits (*s*_*m*_ = 0), the sign of selection in our simple model is directly related to the modifier’s effect on the long-term rate of adaptation (*v*_*m*_ − *v*). This is reminiscent of the mean fitness principle in SI Section 1. Equation (S117) allows us to generalize this result to modifiers with direct costs or benefits (*s*_*m*_ ≠ 0). Interestingly, we find that Eq. (S117) has the same form as the deterministic prediction in Eq. (S5) if we take 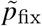 and 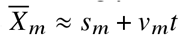, as expected, but impose a finite integration time *T*_max_ = *x*_*cm*_ *v*. This time limit roughly coincides with the fixation time of a successful mutation. This allows us to recover a version of the mean fitness principle in SI Section 1: within our simple clonal interference model, natural selection will favor modifiers that produce a higher mean fitness within a typical fixation time.

We note, however, that Eq. (S117) is only an approximate formula, which captures the leading-order scaling of the logarithm of *p*_fix_ (rather than *p*_fix_ itself). The sub-leading contributions to the logarithm are often important in practice, since they can be large in an absolute sense (≳ 1) and must be exponentiated to obtain *p*_fix_. Thus, while Eq. (S117) can provide a useful rule of thumb, we use the full expressions in SI Section 4.1.5 when calculating our theoretical predictions in Figs. 2 and 3 in the main text.

### 4.2 Quasi-sweep regime

#### 4.2.1 Location of the interference threshold

As we consider larger values of 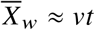, we will eventually reach a regime where a single established driver mutation will be sufficient to drive a modifier lineage to fixation 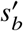. To solve for the shape of *w*_*m*_ (*x*) in this regime, we first revisit the auxilliary condition *x*_*cm*_ for in Eq. (S53). When 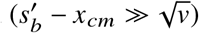, this integral is dominated by fitnesses within 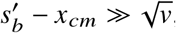 of 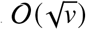, where the shoulder solution has already transitioned to the linear Haldane limit. Performing a Gaussian Laplace approximation around this maximum yields a new auxiliary condition for *x*_*cm*_,

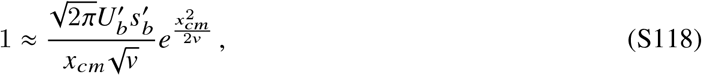

which is valid when 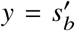 and 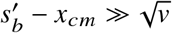. Solving for *x*_*cm*_, we obtain the leading order solution,

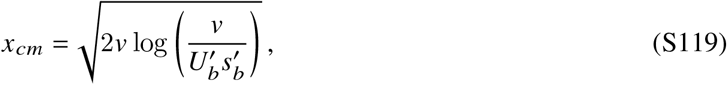

which is valid when

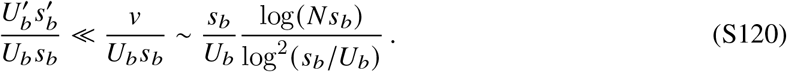

This is a broad regime when *U*_*b*_ ≪ *s*_*b*_, but violations of this condition can become important when we consider continuous distributions of fitness effects below (SI Section 5). It will also be useful to derive an alternative expression for *x*_*cm*_ by substituting the auxilliary condition for *x*_*c*_ in Eq. (S57) into Eq. (S118), yielding

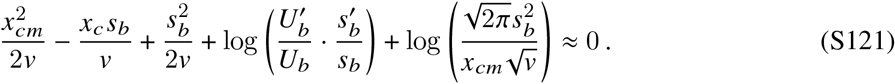

The leading order solution is given by

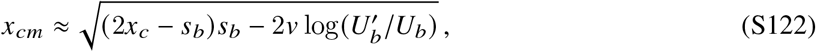

where we have included the potential contribution from 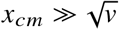, but assumed that the corresponding 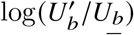 term is subdominant. The condition that 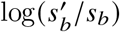 will be valid provided that

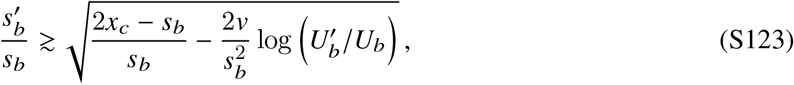

which complements the analogous condition for the mulitple mutations regime in Eq. (S59). Thus, for a pure selection strength modifier, the quasi-sweep regime will occur for 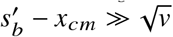, while a pure mutation rate modifier requires the more stringent condition, 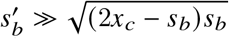. This suggests that relatively modest changes to the selection strength will require the analysis described below.

#### 4.2.2 Extending the shoulder solution to lower fitness values

Having fixed the location of *x*_*cm*_, we can repeat the procedure in SI Section 4.1.2 to compute the shape of *w*_*m*_ *(x*) for fitness values below *x*_*cm*_. If 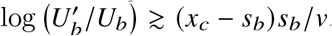, then the dominant contribution to Eq. (S53) will be contained within the region of integration, so that we can substitute the auxilliary condition to obtain

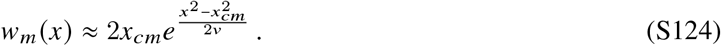

This once again coincides with the shoulder solution in Eq. (S42) in the region where *x < x*_*cm*_ − 𝒪 (*v x*_*cm*_). This derivation shows that the shoulder solution will continue to be valid for fitnesses as low as

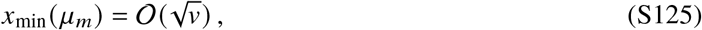

which is significantly smaller than both 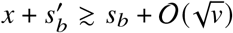 and *x*_*cm*_.

For fitnesses that are less than 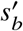, the integral in Eq. (S52) will start to be dominated by the upper limit of integration at 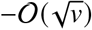. Provided that 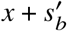, then the dominant contribution to the integral will still come from the Haldane region of the shoulder solution. An exponential Laplace approximation then yields

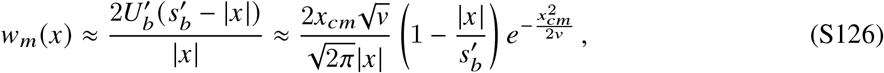

where we have substituted the auxilliary condition in Eq. (S118). To capture the behavior in the intermediate region around 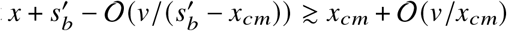, we can turn to the full Gaussian integral,

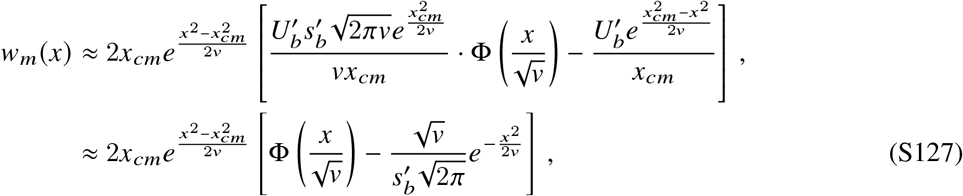

where 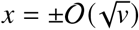 is the Gaussian cumulative distribution function. This reduces to Eqs. (S124) and (S126) in the appropriate limits, but captures the intermediate region where 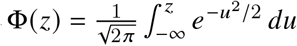.

For fitness below 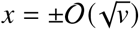, the upper limit of the integral in Eq. (S52) will start to fall within the exponential region of the shoulder solution in Eq. (S42). However, since the latter smoothly maps on to the Gaussian integral in Eq. (S127), we can use this solution to extend *w*_*m*_(*x*) down to *x* ≈ *x*_*cm*_ − 2*s*_*b*_:

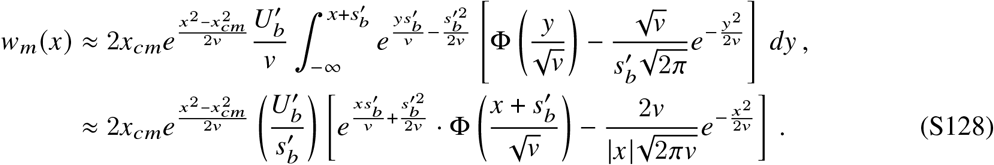

Iterating this procedure again yields

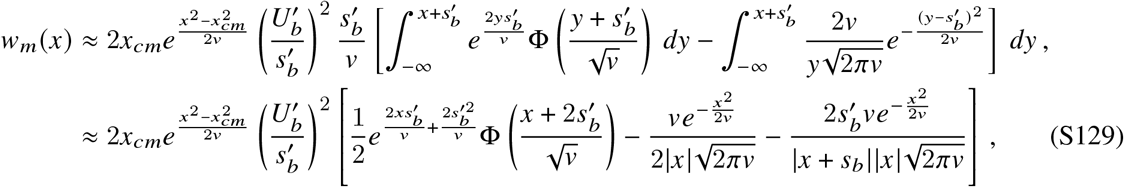

which will be valid for 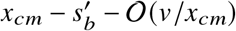. More generally, for 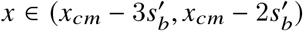, one can show that the solution for *w*_*m*_(*x*) is given by:

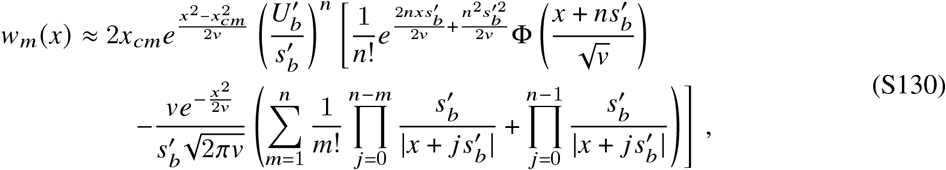

which simplifies to,

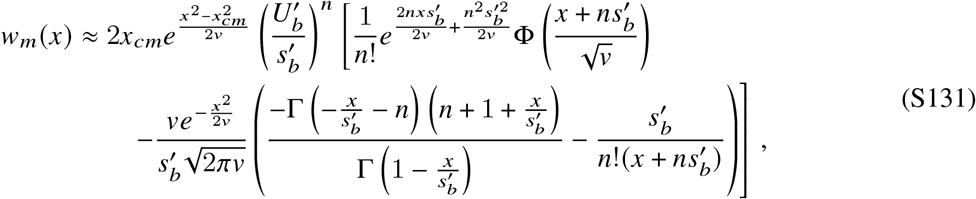

and approaches the asymptotic limit,

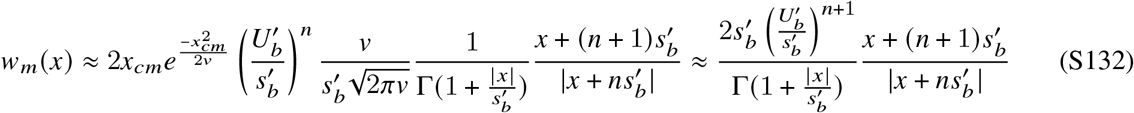

when 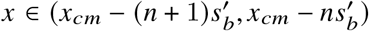. Each additional “step” reduces the original shoulder by a factor of 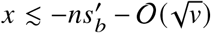. For fitness values that are many multiples of 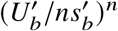 below *x*_*cm*_, we can substitute 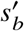to show that the leading order contribution to *w*_*m*_(*x*) scales as

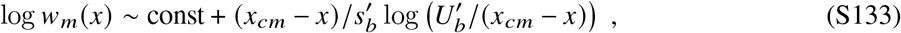

which obeys the required boundary condition that *w*_*m*_(*x*) →0 as *x* →∞. Thus Eq. (S131) provides an asymptotic solution for *w*_*m*_(*x*) that is valid across the full range of relative fitness below *x*_*cm*_ when 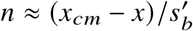.

#### 4.2.3 Fixation probabilities of modifiers

Using the solution for *w*_*m*_ *(x*) in Eq. (S131), we can repeat our calculations from SI Sections 4.1.4 and 4.1.5 to predict the fixation probabilities of modifier mutations in the quasi-sweep regime.

##### Modifiers without direct costs or benefits

In the absence of direct costs and benefits, the primary contribution to the fixation probability in Eq. (S69) will come from the *n* = 0 region of Eq. (S131). This yields

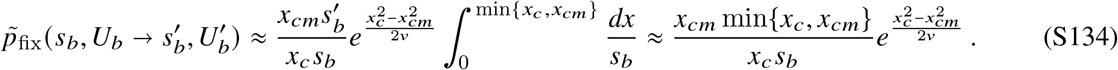

Using Eqs. (S71) and (S118), we can rewrite this expression in the compact form,

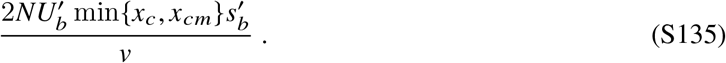

This expression has a simple heuristic explanation. A modifier lineage founded at relative fitness *x* will produce a total of 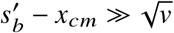 offspring before it is purged from the population. The modifier lineage will produce the bulk of these offspring within 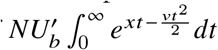 generations of time *t* = *x*/*v*, when it will have reached size 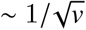 and be near the mean fitness of the population. These offspring will fix provided they survive genetic drift, which occurs with probability 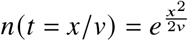. Multiplying these terms together, we see that the advantage of arising on a more fit background and hitchhiking to exponentially larger frequency is exactly balanced by the exponentially smaller probability of arising on one of these privileged backgrounds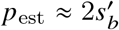. Consequently, a successful modifier is equally likely to have arisen on a background with relative fitness in the range 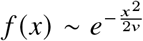.

By comparing Eq. (S90), Eq. (S89), and Eq. (S135) and Eq. (S88), we can see that there is a limit to the degree in which changes to the selection strength of mutations can modulate the effect of linked changes to mutation rate 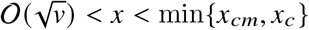. When 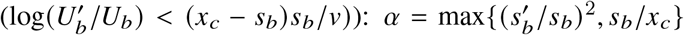, a mutation rate modification influences only a single subsequent mutation compared to multiple mutations when 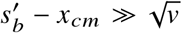. We also note that Eq. (S135) implies that large changes to the mutation rate 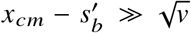 can modulate the effect of a selection-strength modifier, limiting its ability to compound.

##### Modifiers with direct fitness benefits

For a modifier with a direct fitness benefit (*s*_*m*_ *>* 0), the fixation probability in Eq. (S92) will once again depend on how the upper limit of integration (*x*_*c*_ + *s*_*m*_) compares to the interference threshold *x*_*cm*_. If *s*_*m*_ ≳ *x*_*cm*_ − *x*_*c*_ + 𝒪 (*v*/*x*_*cm*_), there will again be a contribution from the Haldane region of *w*_*m*_(*x*), so that

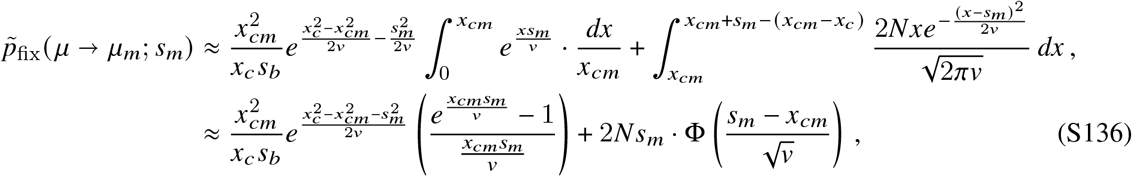

where 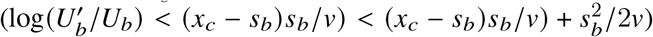 is the Gaussian cumulative distribution function.

In the opposite case, where *s*_*m*_ ≲ *x*_*cm*_ − *x*_*c*_ − 𝒪 (*v*/*x*_*cm*_), the upper limit of integration in Eq. (S92) will fall below the interference threshold of *w*_*m*_(*x*). Repeating the calculation in Eq. (S136) in this case yields

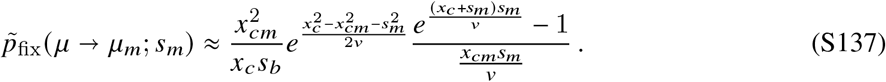

##### Modifiers with direct fitness costs

For a modifier with a beneficial effect on evolvability (*x*_*c*_ *> x*_*cm*_) but a direct fitness cost (*s*_*m*_ *<* 0), the derivation will be similar to Eq. (S99) above. The dominant contribution to the fixation probability in Eq. (S92) will again come from the *n* = *k* term in Eq. (S131) (as well as the first portion of the *k* + 1 term), where *k* is defined by

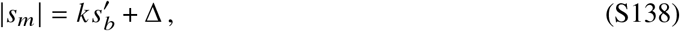

for some 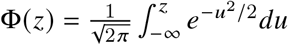. The fixation probability of the modifier can therefore be approximated as

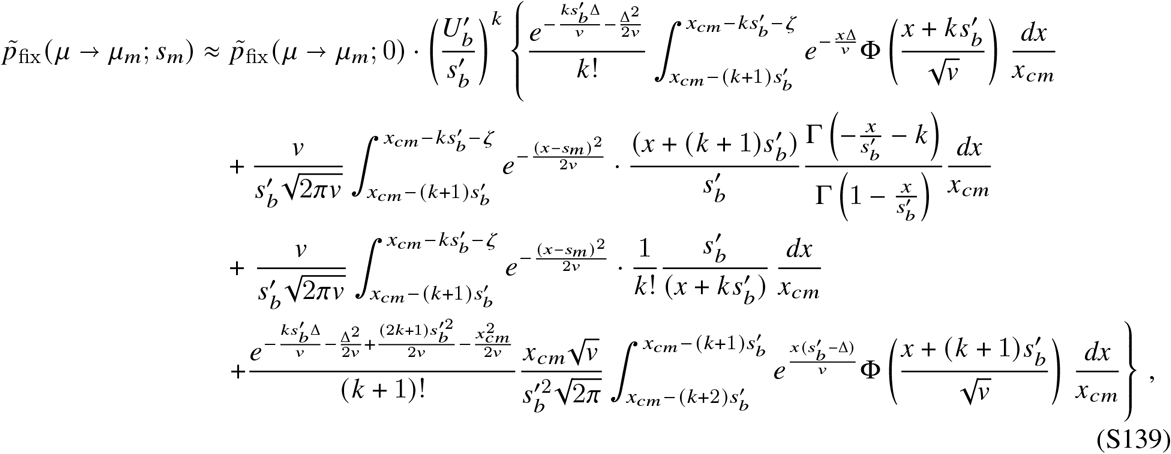

where *ζ* ≡ max{0, Δ − (*x*_*c*_ − *x*_*cm*_)}. When *k* = 0 and 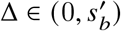, the fixation probability is dominated by the portion of the first term where Φ(*z*) ≈ 1. This yields the asymptotic approximation

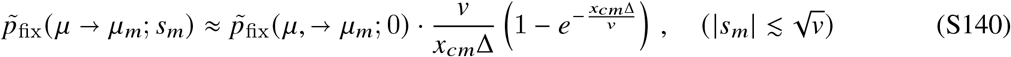

On the other hand, when 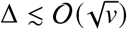, the fixation probability will be dominated by the sum of the first three terms. These contributions will be peaked around a narrow interval near 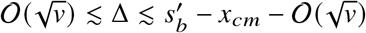, where the sum of the first three terms transitions to the asymptotic expression in Eq. (S132):

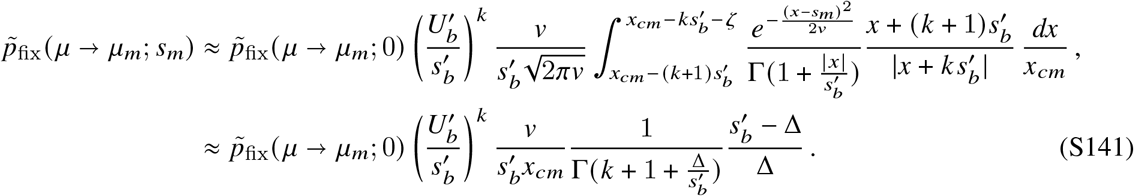

which is valid for 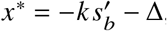. Finally, in the last region where when 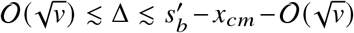, the fixation probability will be dominated by the sum of all four terms, but the first three will be peaked near the lower boundary. This yields

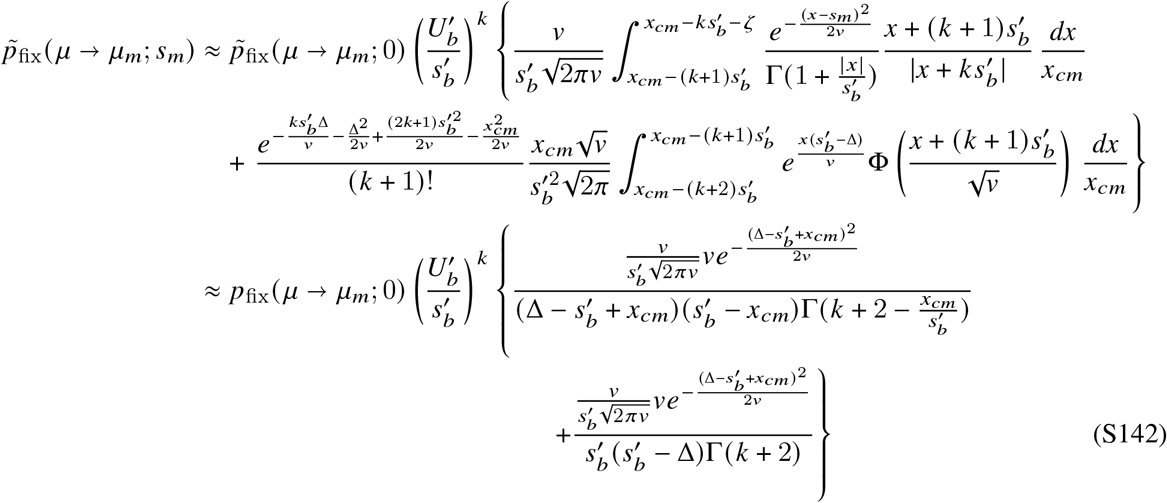

which is valid for 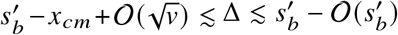. The leading order solution for the fixation probability can therefore be summarized as

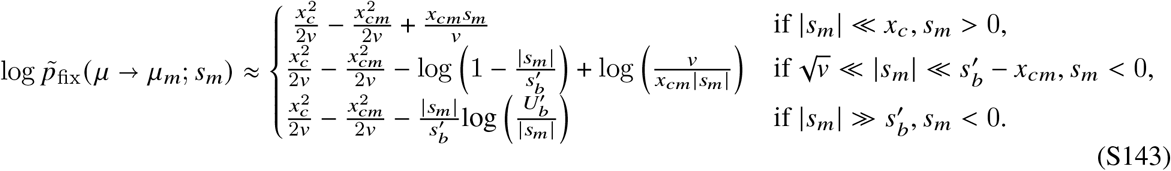

## 5 Extension to continuous distributions of fitness effects

So far, we have focused on a simplified model of the mutation spectrum, where all new mutations confer the same characteristic fitness benefit. In reality, beneficial mutations can have a range of different fitness benefits, so a general evolvability modifier will involve an arbitrary perturbation to the distribution of fitness effects,

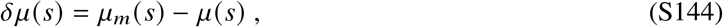

which could include the addition or subtraction of mutations with a range of different fitness benefits. This forces us to return to the full solution in Eq. (S43) of SI Section 3,

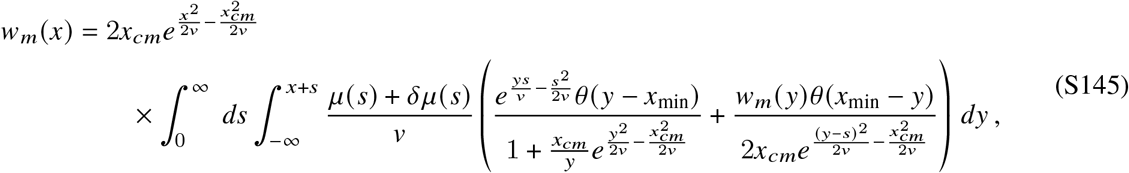

along with the corresponding condition for *x*_*cm*_,

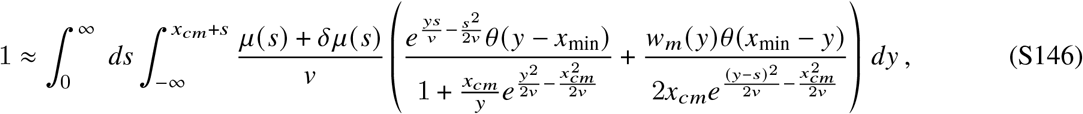

both of which now include an integral over the fitness benefit of the next mutation (*s*) in addition to its landing fitness (*y*). The solutions to this equation are more complicated than the single-effect model in SI Section 4, since the *y* values that contribute the most to each integral will depend on the corresponding value of *s* (and vice versa).

However, previous work (39, 40) has shown that for a broad class of distributions with exponentially bounded tails, the integrals over *s* are dominated by a characteristic fitness effect 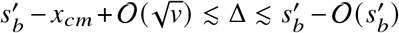 when 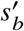. This is a crucial simplification, as it implies that for this range of fitnesses, the shape of *w*_*m*_ *(x*) and the location of *x*_*cm*_ can be well approximated by the single-s solutions in SI Section 4 for some appropriately chosen values of 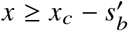 and 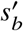 (39, 40, 47, 48). These effective parameters will depend on the underlying values of *μ*_*m*_ *(s*), *x*_*cm*_, and *v*, and must be determined self-consistently using the procedures described below.

In this section, we ask whether this same idea can be extended to the full range of *x* values in Eq. (S145), which is necessary for predicting the modifier fixation probability, *p*_fix_ (*μ* →*μ*_*m*_; *s*_m_). We will see that the single-s approximation holds for certain classes of modifiers, but not others, and we will develop methods for approximating *p*_fix_ (*μ* →*μ*_*m*_; *s*_*m*_) in both cases.

To carry out this analysis, we note that the integral formulation in Eqs. (S145) and (S146) naturally decomposes the fixation probability into contributions from different fitness benefits. By splitting *μ*_*m*_ *(s*) into its components, *μ*_*m*_ *(s*) = *μ (s*) *+δμ (s*), it also allows us to infer the relative contributions of *μ (s*) and *δμ (s*). We can make this connection even more explicit by rewriting Eq. (S146) in the form,

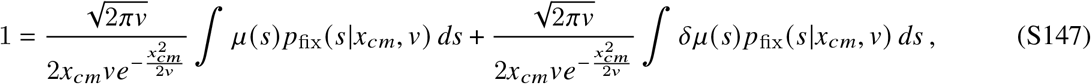

where

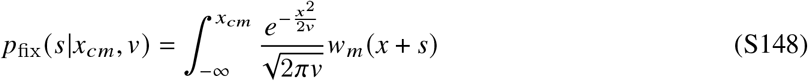

is the fixation probability of a first-order mutation in the “dual” population defined by *v (μ*_*m*_, *N*^∗^)= *v (μ, N*) (see Eq. S37 in SI Section 3).

The decomposition in Eq. (S147) allows us to distinguish between two broad regimes depending on the relative contributions of *μ*(*s*) and *δμ*(*s*). When the fixation probability of the modifier is dominated by mutations from *μ (s*), then 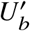 and *x*_*cm*_ will remain close to their wildtype values. This allows us to calculate *x*_*cm*_ and *p*_fix_ (*μ* →*μ*_*m*_; *s*_*m*_) perturbatively, by treating the *δμ (s*) term as a small correction. We will refer to this case as the *perturbative regime*, since it will only apply for modifiers with weak-to-moderate fixation probabilities. For stronger second-order selection pressures, the fixation probability of the modifier will be dominated by mutations from the *δμ*(*s*) portion of *μ*_*m*_(*s*). In this *modifier-dominated regime*, we will need to turn to the full solution of *w*_*m*_(*x*), with *μ*_*m*_(*s*) ≈ *δμ*(*s*). We will discuss each of these cases in turn, after quickly reviewing the wildtype dynamics when *δμ*(*s*) = 0.

### 5.1 Location of the interference threshold for the wildtype population

When *δμ*(*s*) = 0, Eq. (S146) determines the location of interference threshold for the wildtype population,

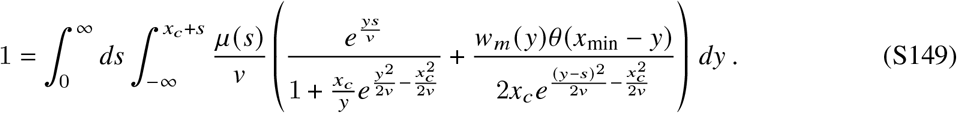

The solution to this equation was previously described by Ref. (39). We reproduce it here for completeness, since we will build on this result in the following sections.

Ref. (39) showed that in our parameter regime of interest, the *s* integral in Eq. (S149) will be strongly peaked at a characteristic value, *s*_*b*_, which will be determined self-consistently below. This observation allows us to evaluate the inner integral over *y* for *s* close to *s*_*b*_. When 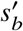*v*, the contributions from *y < x*_*c*_ will be dominated by *y* within 𝒪 (*v*/*s*) of *x*_*c*_, while the contributions from *y > x*_*c*_ will be dominated by fitnesses within 𝒪 (*v*/(*x*_*c*_ − *s*)) of *x*_*c*_. Equation (S149) then reduces to a single integral over *s*,

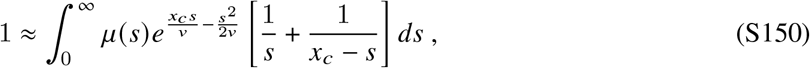

where the rapidly increasing exponential term is eventually counteracted by the rapidly decreasing *μ (s*)term. When 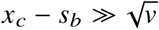, the competition between these two factors produces a sharp peak around *s* ≈ *s*_*b*_ ± Δ*s*_*b*_, where *s*_*b*_ is the value of *s* that maximizes the rapidly varying portion of the integrand,

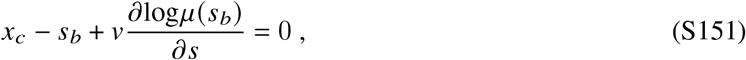

and Δ*s*_*b*_ is the characteristic width,

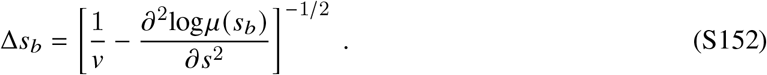

When *s*_*b*_ ≫ Δ*s*_*b*_, we can evaluate Eq. (S150) with a Gaussian Laplace approximation to obtain

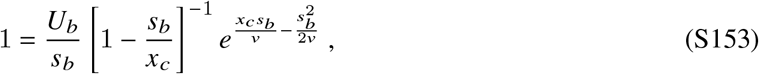

where the effective mutation rate,

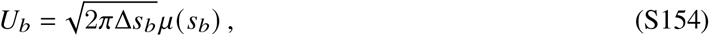

denotes the total rate of producing “driver” mutations within Δ*s*_*b*_ of *s*_*b*_. This matches the corresponding expression for the single-effect model in SI Section 4 if the effective selection strength and mutation rate are defined by Eqs. (S151) and (S154). Unlike SI Section 4, these effective parameters will now depend on the underlying values of *μ s* and *N*, and can shift if one or the other is varied (39, 40, 47, 48). This behavior will be crucial for understanding the fates of modifier mutations below.

#### Application to stretched exponential distributions

Following previous work (39, 40, 47), we can gain some intuition for these expressions by considering the class of stretched exponential distributions,

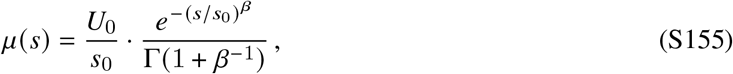

as a function of the tail parameter *β*≥ 1. Ref. (39) showed that the effective parameters for the exponential case (*β* = 1) are given by

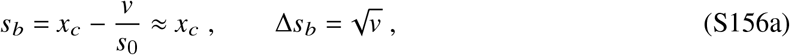

leading to the approximate scaling,

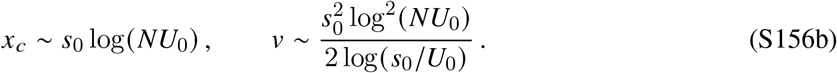

Similarly, the solution for the *β* ≫ 1 case is given by

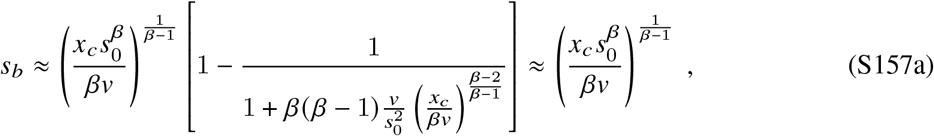

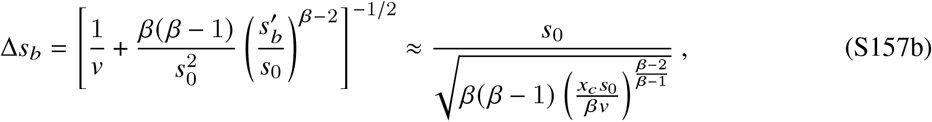

with

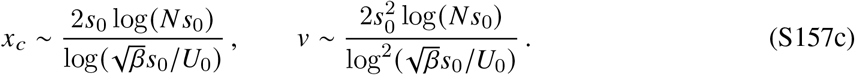

In our analysis below, we will find that the *β* = 2 case (corresponding to a half-Gaussian distribution) will serve as an important boundary case, so it will be useful to derive an explicit expression for this case as well. Equations (S151) and (S152) imply that

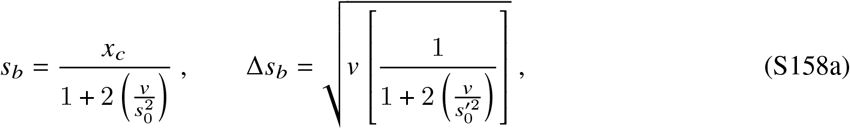

which yields the asymptotic approximations

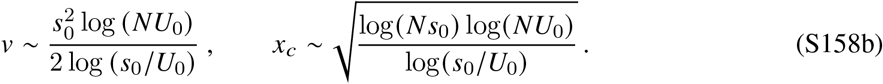

When 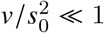, this solution behaves like the exponential case above (with *s*_*b*_ ≈*x*_*c*_). This regime will be valid provided that

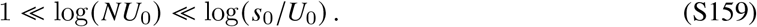

On the other hand, when 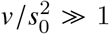, the solution will behave more like the *β* ≫ 1 case (*s*_*b*_ ≪ *x*_*c*_). This regime will be valid provided that

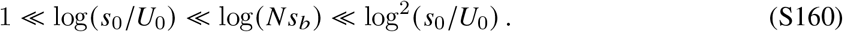

which mirrors the conditions of validity for the single-effect model in SI Section 4.

### 5.2 Perturbative regime

#### 5.2.1 Location of the interference threshold

In the perturbative regime, we assume that the dominant contribution to Eq. (S146) still comes from the *μ (s*) term. This suggests that *x*_*cm*_ will remain close to the wildtype interference threshold,

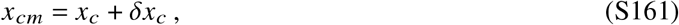

where *δx*_*c*_ is a small correction whose magnitude will be determined self-consistently below.

If *δx*_*c*_ ≪ *v /s*_*b*_, then the integral over *s* will remain strongly peaked around the same value as the wildtype integral in Eq. (S150). This implies that

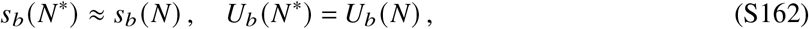

where *s*_*b*_ and *U*_*b*_ are defined as in Eqs. (S151) and (S154), and that

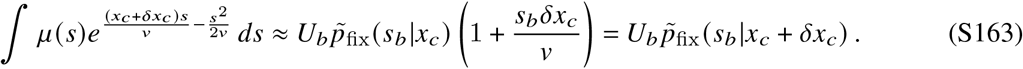

From the self-consistency condition for *N* in Eq. (S71), we also have

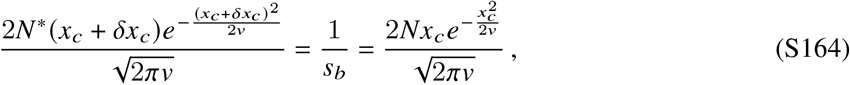

and

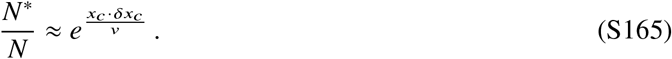

Substituting these results into Eq. (S147), we obtain

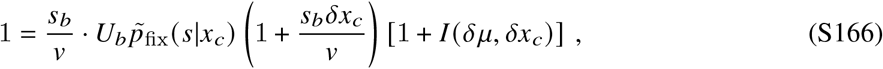

Where

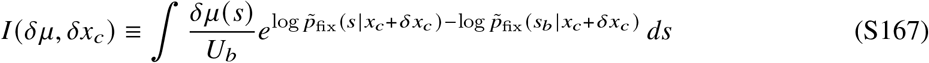

is the integral from Eq. (8) in the main text. Equation (S166) shows that *I* (*δμ, δx*_*c*_) can be interpreted as the relative contribution of mutations from *δμ*(*s*) vs *μ*(*s*) when *δx*_*c*_ ≪ *v*/*s*_*b*_. Substituting for the wildtype *x*_*c*_ condition 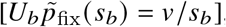, we can rearrange Eq. (S166) to obtain

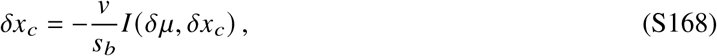

which is valid when *δx*_*c*_ ≪ *v /s*_*b*_. This shows that *δx*_*c*_ ≪ *v /s*_*b*_ is equivalent to the condition that *I (δμ, δx*_*c*_) ≪ 1. If *δx*_*c*_ ≪ *v*/ *s* for the fitness effects that dominate Eq. (S167), then we can perform the same Taylor expansion obtain an analytical expression for *δx*_*c*_,

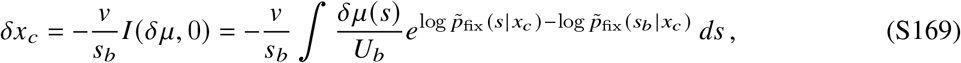

which can be evaluated in terms of the wildtype interference threshold *x*_*c*_. For stronger fitness benefits, *δx*_*c*_ must be obtained by solving the implicit relation in Eq. (S168).

#### 5.2.2 Fixation probabilities of modifiers

When the above assumptions hold, the fixation probability of the modifier can be obtained from our previous calculations in Eqs. (S82) and (S92). Since 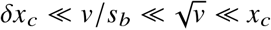, the fixation probability of a modifier without direct costs or benefits will be dominated by exceptionally fit genetic backgrounds (*x*_*c*_ − *s*_*b*_ ≲ *x* ≲ *x*_*c*_), so that Eq. (S82) reduces to,

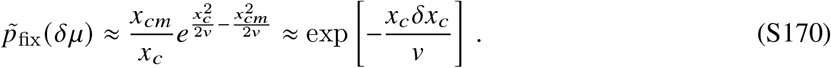

Substituting our expression for *δx*_*c*_ in Eq. (S168) then yields

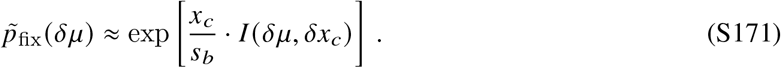

This shows that the conditions of validity for the perturbative regime,

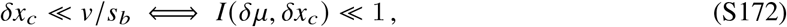

are equivalent to the assumption that

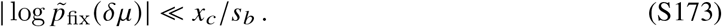

Since *x*_*c*_ can be asymptotically larger than *s*_*b*_, this implies that a small change to the distribution of fitness effects can be strongly selected 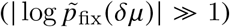 even when it leads to a negligible change in *s*_*b*_ and *U*_*b*_. The perturbative regime is therefore an example of a modifier mutation that cannot be reduced to the single-effect model in SI Section 4.

It is straightforward extend this calculation to consider modifiers that also include direct costs or benefits. Since the second-order selection pressures are bounded by Eq. (S173), most of the interesting behavior will occur when |*s*_*m*_| ∼ 𝒪 (*x*_*c*_/*s*_*b*_) ≪ *s*_*b*_. The fixation probability of these modifiers will continue to be dominated by exceptionally fit genetic backgrounds (*x*_*c*_ − *s*_*b*_ ≲ *x* ≲ *x*_*c*_), so that Eq. (S92) reduces to

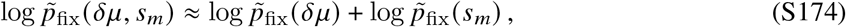

where 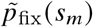 is the fixation probability of a first-order mutation from SI Section 4.1.3.

### 5.3 Modifier-dominated regime (multiple mutations)

For stronger second-order selection pressures 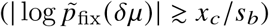, the fixation probability in Eq. (S145) will start to depend more sensitively on the mutations that are added (or removed) by *δμ (s*). For a strongly beneficial modifier, the fixation probability will be dominated by the new mutations that are added by *δμ (s*). In contrast, a strongly deleterious modifier will be dominated by the mutations that remain once *δμ (s)* is subtracted from *μ (s*). In both cases, we will need to develop a general solution for *w*_*m*_ *(x*) for arbitrary distributions *μ*_*m*_ *(s*) ≠ *μ (s*).

Following our analysis in SI Section 4, it will be useful to distinguish between two characteristic regimes, depending on whether the dominant fitness benefits are smaller or larger than *x*_*cm*_. We will continue to refer to these as the *multiple mutations* and *quasi-sweeps* regimes, respectively, since they will qualitatively resemble their analogues in SI Section 4. We will start by analyzing the multiple mutations regime, while the quasi-sweeps case will be considered in SI Section 5.4 below.

#### 5.3.1 Location of the interference threshold

When the relevant fitness benefits in *μ*_*m*_ *(s*) are small compared to *x*_*cm*_, we can repeat the calculation in SI Section 5.1 to obtain an analogous condition for *x*_*cm*_,

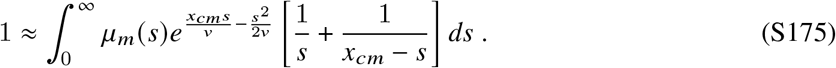

This integral will be peaked around a new value 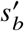 that satisfies

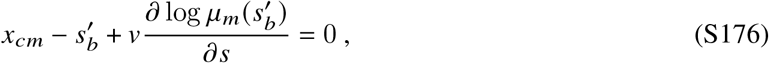

along with a new characteristic width,

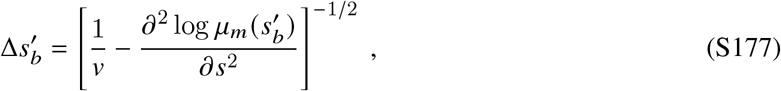

which will generally differ from the wildtype values of *s*_*b*_ and Δ*s*_*b*_ above. When 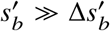 an analogous Gaussian Laplace approximation yields a simple expression for *x*_*cm*_,

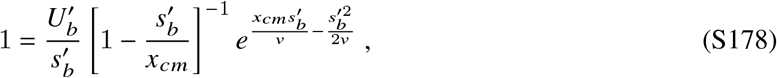

where we have defined the effective mutation rate

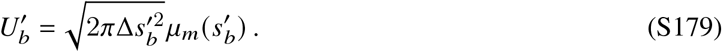

Together, Eqs. (S176-S178) allow us to solve for 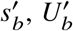, and as a function of *μ*_*m*_ *(s*) and *v*. These expressions will be self-consistently valid when 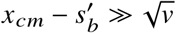.

#### 5.3.2 Extending the shoulder solution to lower fitness values

With the location of *x*_*cm*_ fixed by Eq. (S178), we can repeat our calculation in SI Section 4.1.2 to extend the shoulder solution for *w*_*m*_ *(x*) to progressively lower fitness values. Provided that 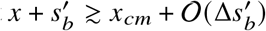, the dominant contribution to Eq. (S145) will also be contained within the region of integration. This demonstrates that the shoulder solution will continue to be valid for fitnesses as low as

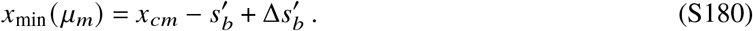

In particular, it shows that for *x > x*_min_(*μ*_*m*_), the shape of *w*_*m*_(*x*) can be approximated by a single-effect model with 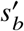 and 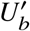 defined by Eqs. (S176) and (S179) above.

##### Hopping vs leapfrogging

When *x* ≲ *x*_min_ (*μ*_*m*_), the upper limit in the landing fitness integral in Eq. (S145) will again become important. However, the crucial difference from our previous analysis in SI Section 4.1 is that this upper limit is not a fixed parameter, but can vary as a function of *s*. In other words, it is theoretically possible for the mutant offspring to land at *any* relative fitness value if it acquires a sufficiently large beneficial mutation from *μ*_*m*_(*s*). This is a new effect that is only present when we allow for a continuous distribution of fitness effects. This will lead to two distinct classes of solutions for *w*_*m*_(*x*) depending on the tail of *μ*_*m*_(*s*).

If the tail of *μ*_*m*_ *(s*) decays sufficiently rapidly, then a modifier lineage that arises below 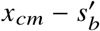 will typically fix by “hopping” to higher relative fitnesses through a sequence of smaller mutations, similar to the single-effect models in SI Section 4. However, if the tail of *μ*_*m*_(*s*) decays more slowly, then a modifier arising below 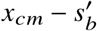 will typically fix by generating a *single* large driver mutation that bypasses these intermediate fitness values, and lands at or above interference threshold at *x*_*cm*_. This “leapfrogging” regime is potentially empirically relevant, since it will apply when *μ*_*m*_ *(s*) decays like an exponential distribution.

By splitting the integral in Eq. (S145) into contributions from above and below the interference threshold, we can obtain a corresponding expression for the fixation probability,

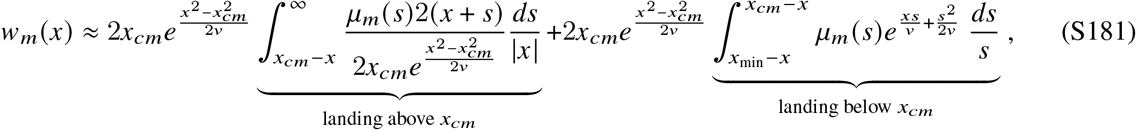

which will be valid for 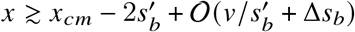. It will be helpful to evaluate these expressions by defining a new variable,

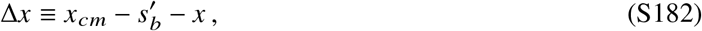

such that 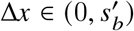.

When Δ*x* ≳ 𝒪 (Δ*s*_*b*_), then the lower limit of the integral in the first term in Eq. (S181) will lie in the high fitness tail of *μ*_*m*_(*s*). An exponential Laplace approximation around 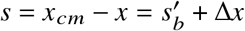 then yields

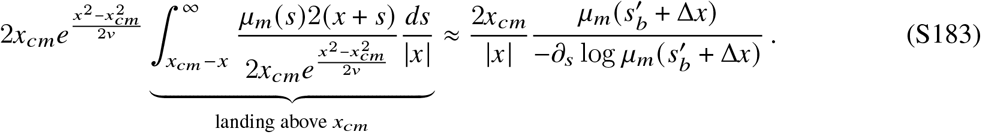

The integrand in the second term in Eq. (S181) has a critical point at *s* = *s*^∗^ (Δ*x*), which satisfies,

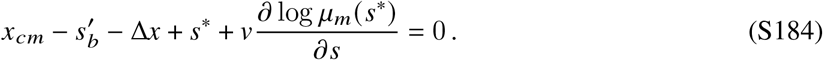

This critical point will correspond to a local maximum if the second derivative is negative,

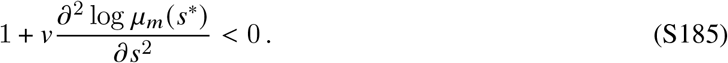

The value of this integral will therefore crucially depend on the tail of *μ*_*m*_ *(s*).

If the curvature of *μ*_*m*_ *(s*) is sufficiently small 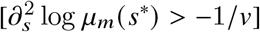, then *s*^∗^ will correspond to a local minimum of the integrand, and the integral will therefore be dominated by the upper limit of integration. An exponential Laplace approximation then yields,

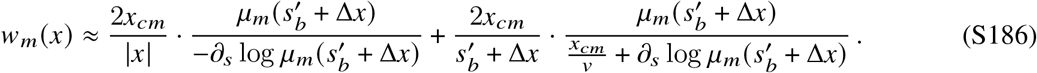

Intuitively speaking, this says that the fixation probability is dominated by mutations of size 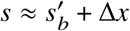 that result in a landing fitness very close to the interference threshold *x*_*cm*_. Since Δ*x* can be as large as 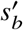 (which is 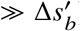), this constitutes an example of the “leapfrogging” behavior above, since it will involve a jump much larger than 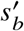. From the form of the condition in Eq. (S185), we can see that this leapfrogging behavior will always occur for the exponential distribution in Eq. (S155), as well as the Gaussian case (*β* = 2) when 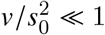. In these cases, we the single-s equivalence will break down for initial fitness below 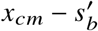, and we must use Eq. (S186) instead.

On the other hand, if the curvature of *μ*_*m*_(*s*) is large and negative 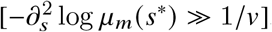, then *s*^∗^ will be a local maximum, and a Gaussian Laplace approximation yields

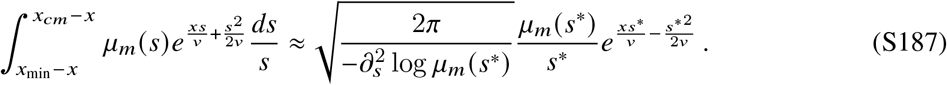

When 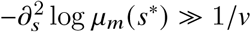, we can also solve Eq. (S184) perturbatively to show that *s*^∗^ (Δ*x*) is close to 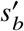:

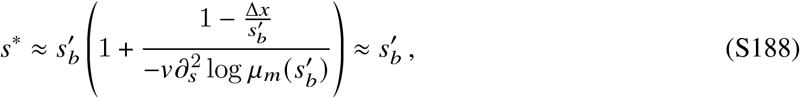

so that Eq. (S181) reduces to

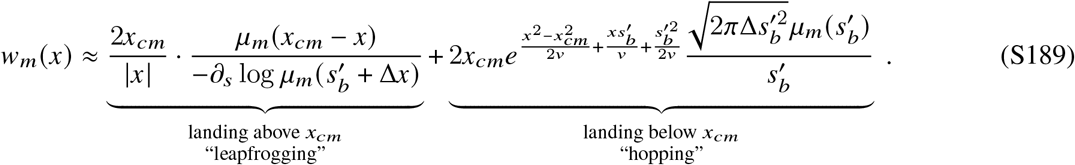

The balance between these two terms will determine whether the typical successful lineage “hops” or “leapfrogs” to fixation. Comparing the magnitudes of these terms, we find that the hopping term in Eq. (S189) will dominate if ′

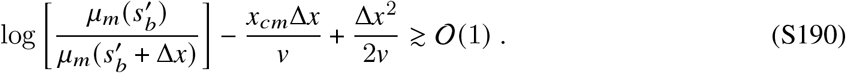

This condition is more stringent than the one defining 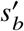 in Eq. (S175), which has an additional 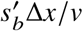 term on the left hand side. This means that even if mutations above 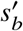 are negligible for 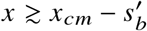, they may become relevant for larger direct costs. When *β* ≥ 2, the decay of *μ*_*m*_ *(s*) will still be bounded by its Gaussian approximation,

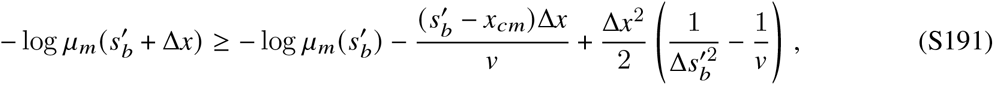

so Eq. (S190) reduces to

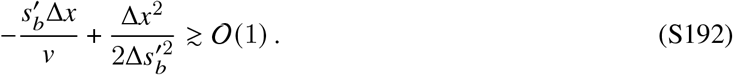

The leapfrogging term will therefore be negligible for

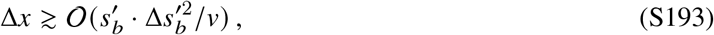

which will constitute the majority of the interval 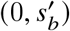when 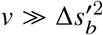 — this is the same condition used to evaluate the hopping term above. For more general choices of *μ*_*m*_(*s*), the Gaussian approximation may no longer bound the value of 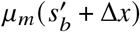, and full condition in Eq. (S190) must be used instead. When these conditions are met, the fixation probability in Eq. (S189) will be well-approximated by the hopping term,

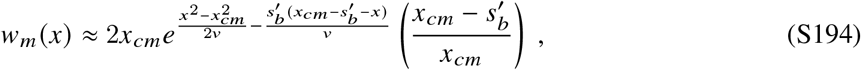

where we have used the auxilliary condition in Eq. (S178) to eliminate the explicit dependence on 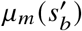. This expression is identical to the result in SI Section 4.1.2, which shows that the single-*s* mapping continues to apply for 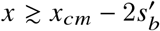.

We can continue these calculations to extend *w*_*m*_(*x*) to lower fitness values. At the next rung 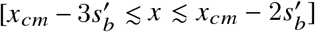, we can once again write *x* as

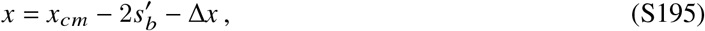

for some 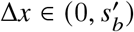, so that Eq. (S145) reduces to

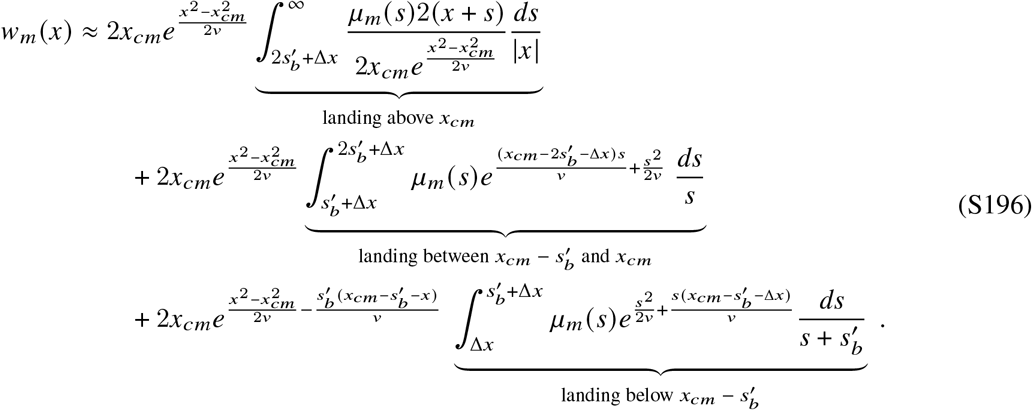

The first term in Eq. (S196) can be evaluated with a similar Laplace approximation as Eq. (S183), yielding

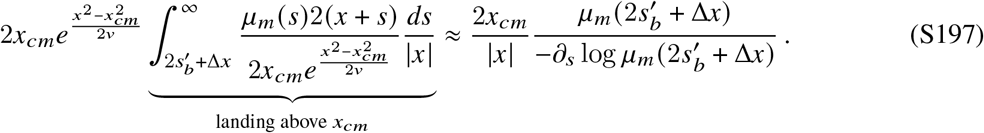

The integral in the third term in Eq. (S196) is similar to the hopping term in Eq. (S181) (up to a prefactor). This implies that it will have a critical point at *s* = *s*^∗^ (Δ*x*) with the same location and curvature as Eq. (S184). Finally, the second term in Eq. (S196) corresponds to intermediate leapfrogging events of size 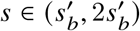. The integrand in this term will have a critical point at a new value, 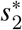, defined by

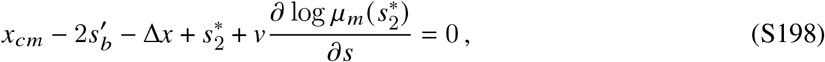

which will generally be less than *s*^∗^. If the curvature of *μ*_*m*_ *s* is small 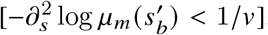, then the critical points in the second and third terms will once again correspond to local minima, and the fixation probability will be dominated by the first term,

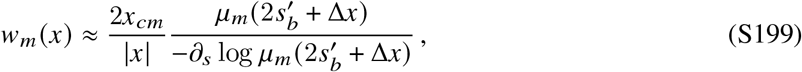

similar to Eq. (S186) above. On the other hand, if 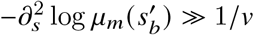, then 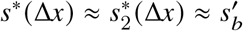 will both correspond to local maxima, and the fixation probability will be dominated by the first and third terms:

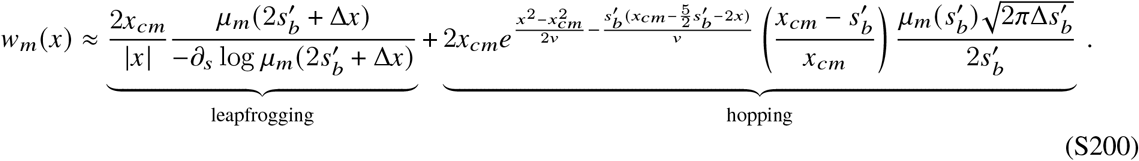

By comparing the magnitudes of these terms, we can see that the hopping term will dominate if

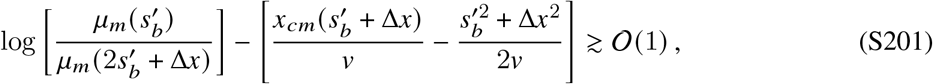

and the corresponding fixation probability will be given by

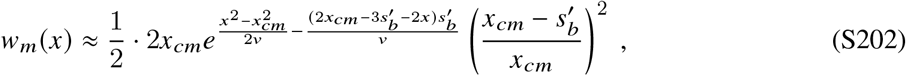

which matches the single-effect result from SI Section 4.1. For the class of stretched exponential distributions in Eq. (S155) with *β* ≥ 2, the mutation spectrum term will again be bounded by its Gaussian approximation,

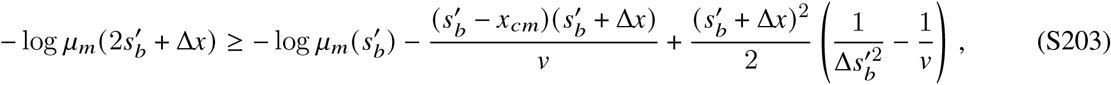

so that Eq. (S201) reduces to

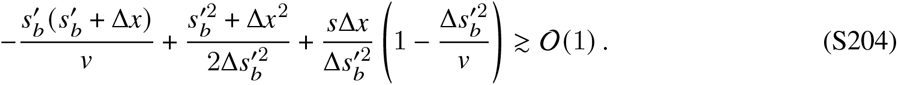

This shows that the hopping approximation will be valid whenever 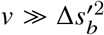, which is the same condition that we obtained for the previous interval above. One can continue this argument to show that for relative fitnesses of the form,

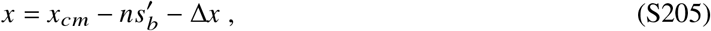

with 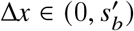), the hopping term will be given by

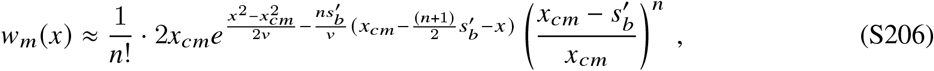

and will constitute the dominant contribution whenever

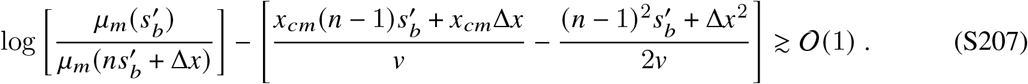

This shows that for the stretched exponential distributions in Eq. (S155) with *β* ≥ 2, the single-*s* approximation will continue to apply for arbitrarily large fitness costs as long as 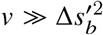.

#### 5.3.3 Fixation probabilities of modifiers

We can now use our solution for *w*_*m*_ *(x*) to evaluate the fixation probability of a modifier mutation. When the dynamics are dominated by hopping, we have seen that the functional form of *w*_*m*_ *(x*) is well-approximated by the single-*s* model in SI Section 4.1. This implies that the fixation probability of the modifier can be predicted using our earlier expressions in SI Sections 4.1.4 and 4.1.5

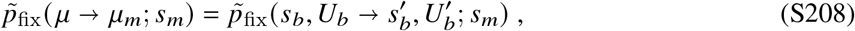

if 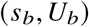 and 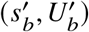 are chosen to coincide with the effective parameters defined in SI Sections 5.1 and 5.3.1 above. This approximation will be valid when 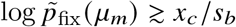and 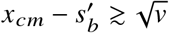. We used this expression to obtain the theoretical predictions for the exponential distribution in Fig. 4B,C.

This single-s approximation will also be valid for *μ*_*m*_(*s*) that allow for leapfrogging, as long as the direct or indirect costs of the modifier are not too strong 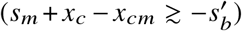. For larger costs 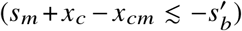, we will need to modify Eq. (S208) to include the additional contributions from the leapfrogging term. Substituting the solution for *w*_*m*_(*x*) in Eq. (S186) into Eq. (S92), we obtain

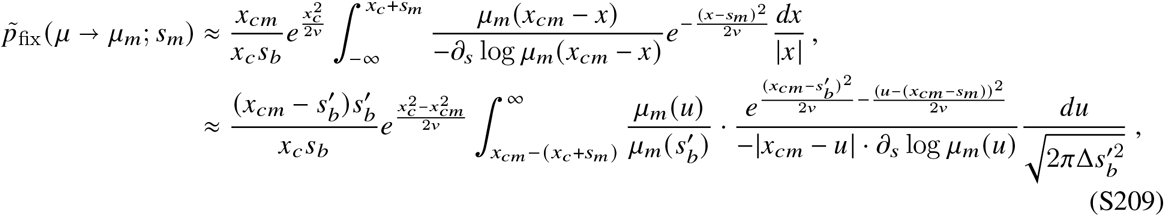

where we have substituted the auxiliary condition for *x*_*cm*_. This integral will have a local maximum at *u*^∗^, where *u*^∗^ satisfies

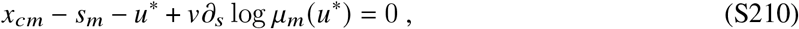

and a characteristic width Δ_*u*_,

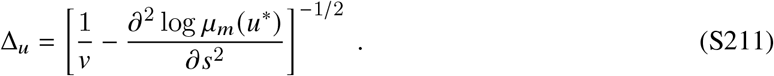

We can therefore evaluate it with a Gaussian Laplace approximation to obtain,

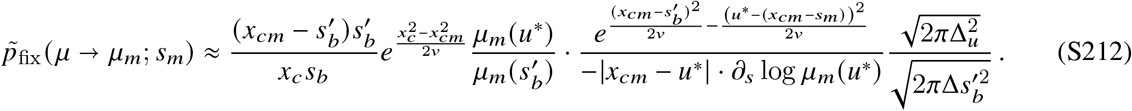

In the case of an exponential distribution, 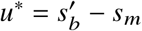 and 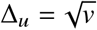, so Eq. (S212) reduces to,

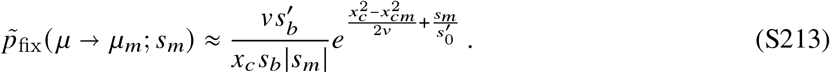

For a half-Gaussian distribution, 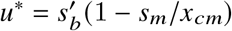 and 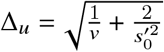, so that

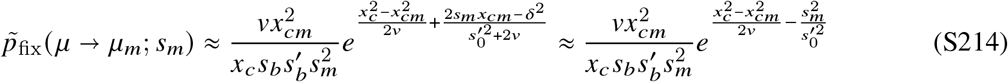

where the last approximation follows when 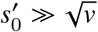. These results imply that the leading order scaling for large direct costs when leapfrogging is valid is given by

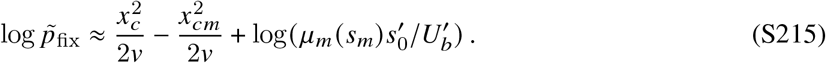

### 5.4 Modifier-dominated regime (quasi-sweeps)

The results in the previous section apply when 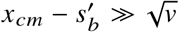. As the gap between *x*_*cm*_ and 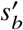 shrinks, we will eventually reach a point where a single established mutation will be sufficient to drive the modifier lineage to fixation. To solve for the shape of *w*_*m*_(*x*) in this “quasi-sweeps” regime, we must revisit the equations for *w*_*m*_(*x*) and *x*_*cm*_ in Eqs. (S145) and (S146).

#### 5.4.1 Location of the interference threshold

Motivated by our solution for the quasi-sweeps regime in SI Section 4.2, we anticipate that the dominant contributions to the y-integral in Eq. (S146) come from the Haldane region of the shoulder solution (*y > x*_*cm*_). When 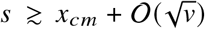, the integrand will be peaked around a characteristic value 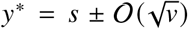. Performing a Gaussian Laplace approximation around this maximum yields,

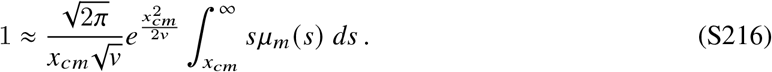

This is similar to the single-effect model in SI Section 4.2.1 if we take

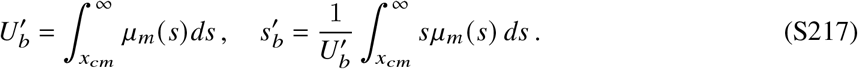

The decomposition in Eqs. (S145) and (S146) shows that this 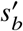 integral coincides with the distribution of fixed mutations in the modifier lineage. In contrast to the multiple mutations regime in SI Section 5.3, the distribution in Eq. (S217) may have a typical scale but will generally not be strongly peaked.

This quasi-sweeps regime will be self-consistently valid if Eq. (S216) is much larger than the remaining contributions from *y* ≤ *x*_*cm*_:

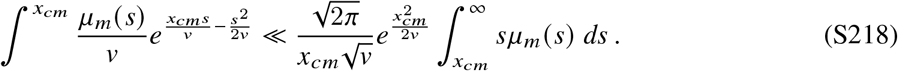

For the stretched exponential distributions in Eq. (S155), this will only be true if 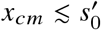, so that 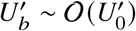and 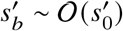. This shows that the quasi-sweeps regime will be valid when

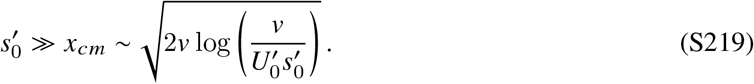

#### 5.4.2 Extending the shoulder solution to lower fitness values

We can continue this line of reasoning to compute the shape of for fitness values below. For mutations with fitness effects 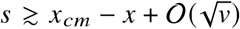, the y-integral in Eq. (S145) will continue to have a large contribution from the Haldane region of the shoulder solution. For these values of *s*, we can repeat the Gaussian integration in Eq. (S127) to obtain

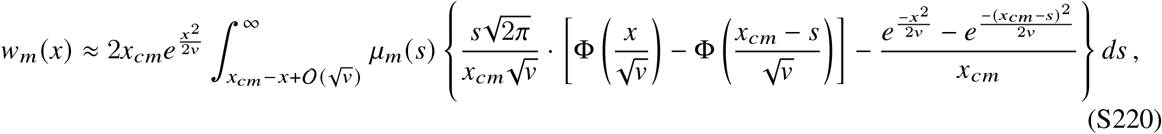

where 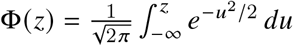 is the Gaussian cumulative distribution function. When 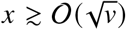, the Gaussian cumulative function is close to one, and Eq. (S220) reduces to

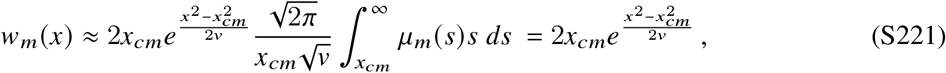

where the last line follows from substituting the auxiliary condition for *x*_*cm*_ in Eq. (S216). This shows that the shoulder solution will be valid down to 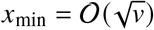, similar to the single-effect model in SI Section 4.2.2.

When 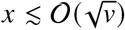, the shape of Φ(*x*) will start to become important. At this point, the boundary terms at *x*_*cm*_ − *s* will become negligible, so that Eq. (S220) reduces to

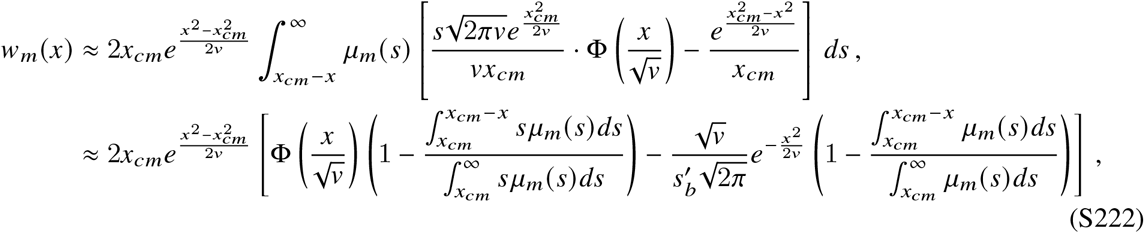

where the second line follows from substituting the auxiliary condition for *x*_*cm*_ in Eq. (S216). This will coincide with the analogous expression for the single-*s* model in 4.1.2 if

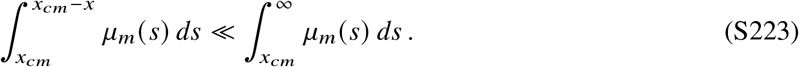

This condition will be satisfied if 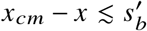, or alternatively, if 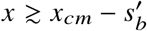.

For lower initial fitnesses, the contributions from mutations that land below the interference threshold (*x* + *s < x*_*cm*_) can begin to become important. We can estimate the onset of these effects by considering initial fitnesses in the range 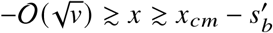. In this limit, Eq. (S222) reduces to

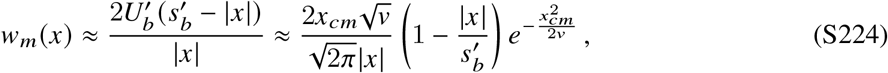

where the second line follows from substituting the auxiliary condition for *x*_*cm*_ in Eq. (S216). The corresponding contributions from mutations with *x*+ *s < x*_*cm*_ in Eq. (S145) are dominated by the upper limit of the *y*-integral, which yields an additional correction of order

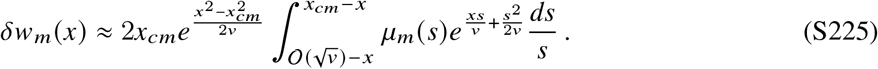

If the curvature of *μ*_*m*_ *(s*) is not too strong 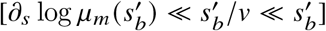, this *s*-integral will be dominated by the upper limit of integration, so that

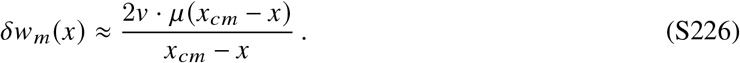

This will be small compared to Eq. (S224) as long as 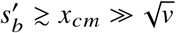, which shows that Eq. (S222) will be a good approximation for *w*_*m*_(*x*) as long as 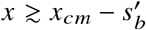.

For lower initial fitnesses 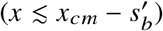, these additional contributions will start to become important.

In principle, we can generalize the recursive approach in SI Section 4.2.2 to extend our solution for *w*_*m*_ *(x*) below this point. However, these corrections will no longer be universal, and will depend on other features the mutation spectrum beyond the first two moments in Eq. (S217). Since our results in the main text will only require the portion of the solution for 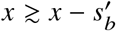, we will leave these calculations for future work.

## 6 Incorporating deleterious mutations

While the previous sections have focused on changes to the beneficial mutation spectrum, the vast majority of new mutations are neutral or deleterious (45). The most general evolvability modifiers could alter the rates and fitness costs of these deleterious mutations as well.

Previous work (46, 103, 105) has shown in our parameter regime of interest, deleterious mutations can be divided into two broad categories: (i) quasi-neutral mutations (|*s*| ≪ /*v x*_*c*_), which can frequently hitchhike to fixation with beneficial mutations, and (ii) “purgeable” deleterious mutations (|*s*| ≫*v* /*x*_*c*_), which rarely reach high frequencies, but can collectively reduce the effective population size if their mutation rate is sufficiently high. Previous estimates suggest that the overall mutation rates in many natural and laboratory microbial populations are sufficiently low (*U* ≲ 10^−3^ (106)) that both categories have a negligible impact on the overall the rate of adaptation (46, 103). However, differences in the deleterious distribution of fitness effects can still have a strong influence on the fixation probability of a modifier mutation (24).

We can incorporate purgeable deleterious mutations into our framework using the argument outlined in Ref. (24), which we reproduce here for completeness. Ref. (24) showed that it is useful to rewrite the mutation term in Eq. (S33) as separate beneficial and deleterious contributions,

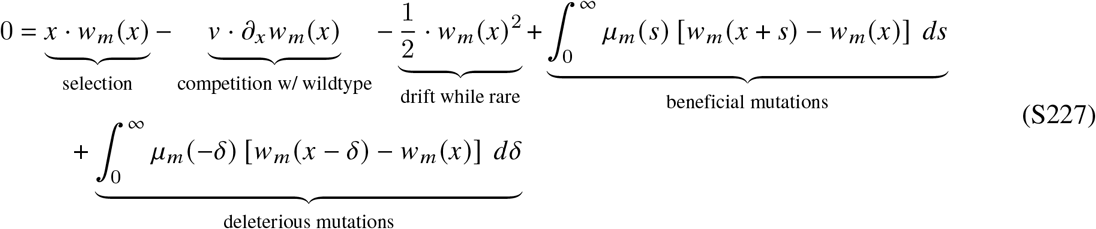

where we are indexing the deleterious mutations by their overall magnitude *δ*≡− |*s*|. The definition of a purgeable mutation is that they rarely reach high frequencies, which implies that the fixation probability after acquiring a purgeable mutation must be close to zero. If all of the deleterious mutations in *μ*_*m*_ fall in this purgeable category, we can therefore neglect the *w*_*m*_ *(x* −*δ*) term in Eq. (S227). This allows us to rewrite Eq. (S227) in the simpler form,

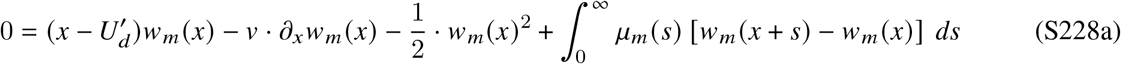

where 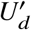 denotes the total mutation rate for purgeable mutations:

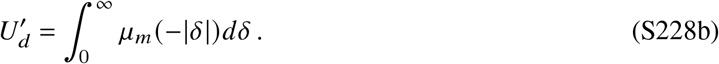

This equation has the same form as the purely beneficial case analyzed above, but with the *x* coordinate shifted by 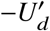. The solutions can therefore be expressed in the simple form

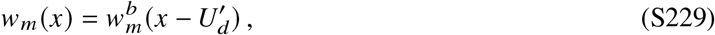

where 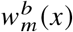 denotes our existing results for the purely beneficial case. Likewise, previous work (24, 46) has shown that the background fitness distribution satisfies an analogous formula,

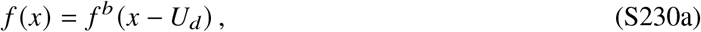

where *U*_*d*_ is the rate of producing purgeable mutations in the wildtype background,

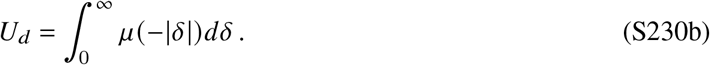

Substituting these expressions into Eq. (2), we find that the fixation probability of the modifier can be written as

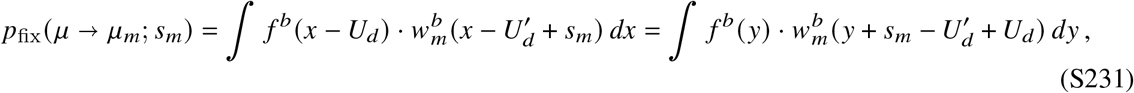

which implies that differences in the purgeable mutation rate behave like effective direct cost,

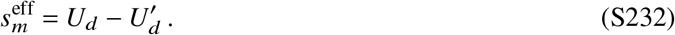

We validated this approximation using Wright-Fisher simulations (Fig. S2), and found that it encompasses a broad range of deleterious fitness effects, including those much smaller than a typical beneficial driver mutation (Fig. S2B). Deviations eventually occur for more weakly selected mutations, which have a lower impact on the fixation probability than Eq. (S232) would predict. This suggests that purgeable mutations represent an upper bound on the strength of second-order selection.

Interestingly, the mapping in Eq. (S232) implies that the same modulation effect that occurs between first- and second-order selection will also apply for selection on simultaneous changes to robustness 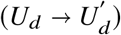 and evolvability (*μ* →*μ*_*m*_). In particular, it implies that larger populations are more likely to trade reduced robustness for long-term gains in evolvability. Since empirical deleterious mutation rates are often comparatively low 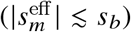, even the maximum possible reduction in robustness will be overpowered by marginal increases in evolvability (Fig. S2C). The opposite scenario – greedily selecting for increased robustness despite long-term reductions in evolvability – is also theoretically possible, but we expect that it will be less relevant in practice due to the low deleterious mutation rates in many organisms.

Finally, we note that robustness and evolvability do not have to trade off with one other – several recent studies have shown that they can sometimes interact synergistically as well (61, 63). Our analytical framework also provides new predictions for these synergistic mutations, allowing us to determine how each phenotype contributes to the lineage’s long-term evolutionary fate (Fig. S2E). Our results show that larger populations will tend to weigh proportional enhancements in evolvability (Δ*v*/ *v*) more heavily than comparable increases in robustness (Δ*U*_*d*_ /*U*_*d*_). This is qualitatively different from the successive sweeps picture (SI Section 2), which predicts that the opposite ordering should occur. These examples illustrate how clonal interference can reshape our intuition about second-order selection for evolvability.

## 7 Extensions to more general fitness landscapes

Our analysis above focused on the simplest possible model of an evolvability modifier, where the local distribution of fitness effects could be approximated by a pair of fixed distributions,

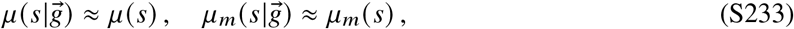

for all genotypes 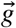(SI Section 3.1). In this section, we consider several extensions of our model that relax this assumption in different ways. In each case, we find that our existing theory provides a useful baseline for incorporating these new effects.

### 7.1 Weak Macroscopic Epistasis

It is clear that some deviations from Eq. (S233) will be too small to be relevant to natural selection. We can formalize this idea by considering subtle deviations of the form,

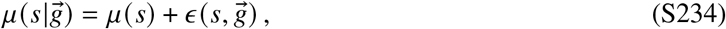

where the function 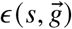 represents a small perturbation. In this case, our existing results in SI Section 5 allow us to precisely define what we mean by “small”. In particular, if we let 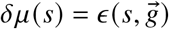, then we can conclude that any perturbation for which the integral Eq. (8) is small [*I (δμ*) ≲ *s*_*b*_ *x*_*c*_] will be essentially invisible to natural selection. Any deviations from Eq. (S233) that fall below this minimum resolution will therefore not affect our main results.

We note that this space of “negligible” perturbations can be quite large from the perspective of the original DFE. Fig. 4 shows that even large fluctuations in *μ s* can be tolerated for fitness effects ≲ *s*_*b*_ *μ, N*, even if they cause in large shifts in the overall mean and height of the DFE. Conversely, much smaller perturbations at fitness effects ≳ *s*_*b*_ *μ, N* can lead to important deviations from our original model, even if they nominally appear to satisfy Eq. (S233).

We also note that this argument constitutes an upper bound on the impact of a given deviation from Eq. (S233), since our derivation of Eq. (8) assumed that the evolvability differences were permanent. This means that some values of *ϵ s, g* that exceed the resolution limit above could still have a small influence on the results because their evolvability differences are only transient. We consider such cases in more detail below.

### 7.2 Transient differences in evolvability

In many cases of interest, the evolvability benefits of a modifier will not persist indefinitely (as in Eq. S233) but will only apply within some local region of genotype space. For example, in the stability-activity landscape in Fig. S1C, the stability enhancing modifier creates an opportunity for *K* additional mutations to accumulate. At a formal level, this constitutes a large deviation from Eq. (S233).

However, it is also clear that such permanent shifts are not truly necessary. Second-order selection can only take place while the modifier is competing with the wildtype; once one of these lineages fixes, future changes in 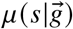 can no longer contribute to the fixation probability. Our results above allow us to estimate the size of this critical window. In particular, our heuristic analysis in SI Section 4.1.4 shows that the benefits of an evolvability modifier accumulate over 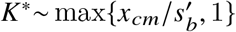 mutational steps. This suggests that changes to the DFE that occur outside this horizon will have a negligible impact on our results.

We tested this prediction by considering a simple model where the modifier reverts to the wildtype DFE after *K* mutational steps:

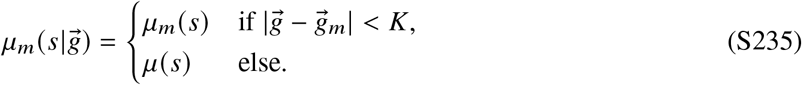

Simulations of this model show that our existing theory continues to provide a good approximation even when *K* is as low as 2 or 3 (Fig. S3A), consistent with the moderate values of *q* ≡ *x*_*c*_ */s*_*b*_ that are attained for many empirically relevant parameter values (39, 47). Similar results are also observed if the modifier reverts to a dead-end rather than the wildtype (Fig. S3B), with differences arising only for the smallest values of *K*. In both cases, we find that the evolvability benefits still confer exponentially large advantages over the classical SSWM expectation (Fig. S3A), even when they last for just a single mutation (*K* = 1).

### 7.3 Global diminishing returns epistasis

Many evolving populations exhibit a form of diminishing returns epistasis, where the fitness benefits of new mutations systematically decline with a lineage’s absolute fitness (9, 10, 41, 54, 55, 89). To understand how this phenomenon might impact our results, we considered a simple model of global epistasis inspired by the long-term evolution experiment in Fig. 1A (55), where the typical fitness benefits of new mutations decline exponentially with the total fitness,

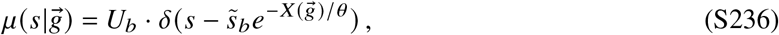

where *θ* controls the strength of the diminishing returns effect. In the simplest scenario, we can assume that the modifier distribution exhibits a similar decline,

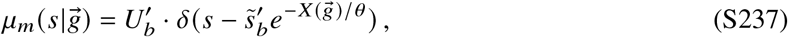

but with different values of 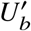 and 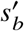. This generalizes the simple toy model in Fig. 2 to allow for steadily declining fitness effects.

#### Adiabatic approximation

The simplest behavior occurs when *θ* is sufficiently large. Suppose that the modifier arises at time *t*_0_ when the mean fitness of the population is 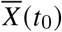. If we could neglect the additional deceleration that occurs over the lifetime of a single mutation, we could simply apply our existing results with 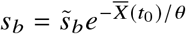 and 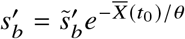. We can use our heuristic picture in SI Section 4.1.4 to determine when this “adiabatic approximation” will be valid. The fate of a mutation is determined over *T*_*c*_ *∼x*_*c*_ */v* generations, during which time the total fitness of the population increases by 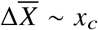. If *θ ≫ x*_*c*_, then diminishing returns epistasis will have a negligible effect over the lifetime of a single mutation, even if it had a large effect in setting the initial scale of selection 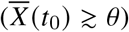. This provides a self-consistent justification for our adiabatic approximation above.

This simple picture leads to novel predictions when combined with our existing results in Eqs. (3) and (4). In particular, it suggests that a selection-strength modifier will become less strongly selected over time as the population becomes better adapted, while the benefits of a mutator allele will remain roughly constant over the same time window. Since *x*_*c*_ is roughly proportional to *s*_*b*_ (Eq. S49), we expect that *θ /x*_*c*_ will grow increasingly large at long times as *s*_*b*_ becomes progressively smaller (Fig. S4). This suggests that the adiabatic approximation will often be useful for understanding the long-term dynamics of a population (41).

#### Beyond the adiabatic approximation

The situation becomes more complicated when *x*_*c*_ is comparable to *θ*, since the fitness benefits of new mutations will now decline within the scale of the population fitness distribution. The dynamics in this regime are poorly understood even in the absence of second-order selection (41). However, we can still account for these effects in a crude way by leveraging our heuristic picture in SI Section 4.1.4, and focusing on the leading-order corrections when *x*_*c*_ */θ* is small but finite. We briefly review the results for our original model below, and then show how they can be extended to calculate the leading-order corrections from diminishing returns epistasis.

Recall that for small changes in the selection coefficient 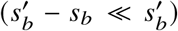, successful modifiers will typically arise in the high-fitness nose of the population (*x* ≈ *x*_*c*_) and acquire *x*_*c*_/*s*_*b*_ mutations before reaching appreciable frequencies (SI Section 4.1.4). In each of these steps (*j* = 1, …, *x*_*c*_/*s*_*b*_), a selection-strength modifier will grow for *τ*_est_ ≈ *s*_*b*_/*v* gen erations while t he next nose establishes. During this establishment time, the modifier will produce 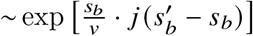 more mutations than a wildtype individual with the same fitness, leading to the scaling

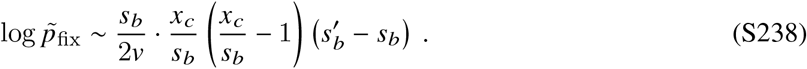

This picture becomes more complicated with diminishing returns epistasis since the wildtype and modifier selection coefficients will decline with each subsequent mutation,

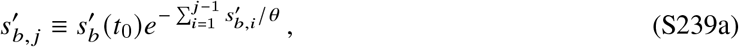

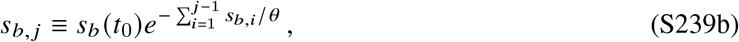

where 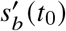 and *s*_*b*_ *(t*_0_) denote the effective selection coefficients at the time that the modifier arises. As a result, the modifier’s growth advantage, and the time it spends growing, will also vary with each mutation. We can straightforwardly extend our heuristic prediction to account for these differences,

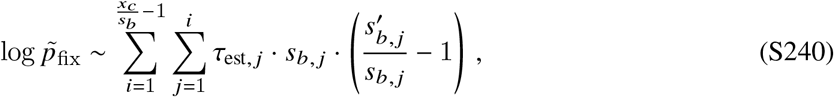

where *τ*_est, *j*_ denotes the establishment time of the *j* ^th^ mutational step. These establishment times will emerge from the non-equilibrium dynamics imposed by diminishing returns epistasis, including the declining rate of fitness increase and width of the fitness distribution. While these dynamics remain poorly characterized even in the absence of second order selection, we find that we can obtain accurate predictions by assuming that the product *τ*_est, *j*_ · *s*_*b, j*_ remains approximately constant during the lifetime of the modifier:

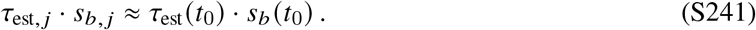

This assumption is motivated by the logarithmic dependence of *τ*_est_ · *s*_*b*_ on *s*_*b*_ in the absence of diminishing returns epistasis (47; Eq. S49), which suggests that treating it as constant over fixation timescales will often be a good approximation. With this assumption, Eq. (S240) reduces to the simpler form

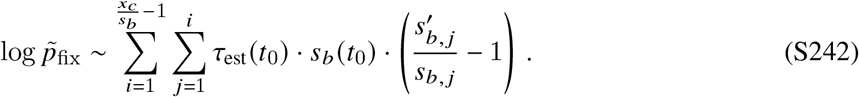

Under the model in Eq. (S239), the diminishing returns effects on 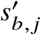 and *s*_*b, j*_ exactly cancel each other, so that we end up with the same adiabatic approximation above. This suggests that the next-order corrections remain small when the modifier and wildtype follow similar diminishing returns schedules.

We can also consider modifiers that alter the form of the diminishing returns epistasis itself. The simplest example is one that changes the diminshing returns parameter *θ* to a new value *θ*_*m*_ once the modifier mutation arises:

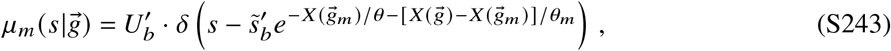

where 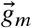 denotes the founding genotype of the modifier lineage. The difference between *θ* and *θ*_*m*_ is invisible in the adiabatic approximation, but the next order corrections can be obtained from Eq. (S242), by substituting

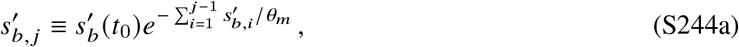

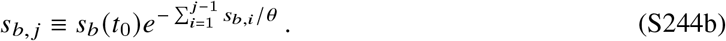

The diminishing returns contributions no longer cancel, leading to modest corrections to our existing theory (blue lines, Fig. S4D). In particular, when *θ*_*m*_ ≫ *θ*, these diminishing returns modifiers can be positively selected even when their initial DFEs are the same 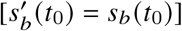, since the modifier experiences fewer diminishing returns effects at later times. These differences disappear once 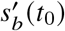 is sufficiently large that we enter the quasi-sweeps regime. In this case, we have seen that the success of the modifier is driven by its ability to acquire its first additional mutation (SI Section 4.2.3), where the new value of *θ*_*m*_ has not yet taken effect. These extensions of our heuristic analysis suggest that the simple model we have studied in this work may be a powerful tool to understand selection for evolvability on more general fitness landscapes.

## 8 Forward-Time Simulations and Numerical Methods

### 8.1 Forward-time simulations

We validated our theoretical predictions by comparing them to forward-time, Wright-Fisher-like simulations similar to those employed by Ref. (24). We begin with a population of *N*_*t*=0_ ≡ *N* individuals at generation 0. In each generation, every individual *i* in the population produces a Poisson number of identical offspring with mean

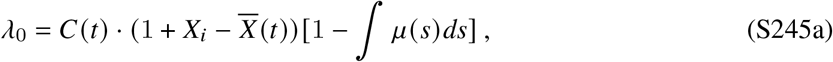

and a Poisson number of mutant offspring with mean

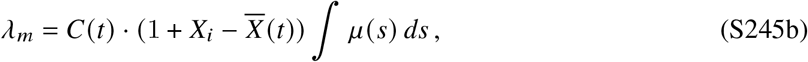

where *X*_*i*_ is the absolute fitness of individual *i*, 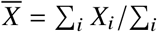 is the mean fitness of the population, and *C (t*) = *N /N*_*t*_ is a normalization constant that ensures that the expected population size in each generation is equal to *N*. Each mutant offspring is assigned a new fitness value *X*_*i*_ + *s*, where *s* is randomly sampled from the normalized version of *μ (s*). After initialization, each simulation is allowed to “burn in” for Δ*t* = 2 10^4^ generations.

To measure the fixation probability of a modifier lineage, we continue this basic algorithm while allowing individuals in the background population to produce new modifier offspring at a per capita rate *U*_*m*_. These modifier individuals produce offspring according to an analogous version of Eq. (S245), with *μ (s*) → *μ*_*m*_ *(s*). Reversions from modifier to wildtype genotypes are not allowed. Following the burn in period, we record the number of generations *T* that elapse until a modifier lineage fixes in the population. In the limit that *U*_*m*_→ 0, the fixation time *T* is related to the fixation probability through the relation,

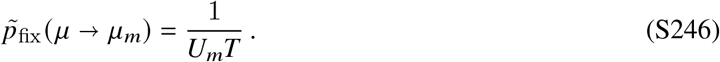

For the estimator in Eq. (S246) to apply, *U*_*m*_ must be small enough that the total time to fixation time *T* is much larger than the predicted sweep time of a successful modifier (*x*_*cm*_ */v*). To ensure that this is true, we repeat this process of measuring *T* for *M* = 60 replicate populations with a sequence of modifier mutation rates *U*_*m*,1_, *U*_*m*,1_, …*U*_*m,M*_, yielding a sequence of fixation times *T*_1_, *T*_2_, … *T*_*m*_; the mutation rate at step *i* is chosen based on the previous *T*_*i*−1_,

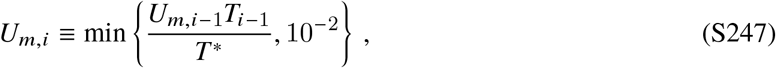

so that the predicted value of *T*_*i*_ is *T* ^∗^ 250· *x*_*cm*_ */v* generations. The mutation rate is also capped at 10^−2^ so that a successful modifier lineages always competes against the background population while small. The sequence is started at *U*_*m,i* 1_ = 1 *T* ^∗^ and allowed to “burn-in” for 10 iterations before the *T*_*i*_’s are recorded. The fixation probability is calculated from the maximum likelihood estimator,

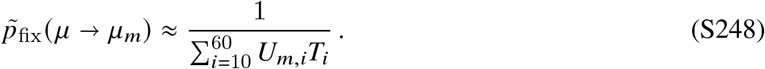

A copy of our implementation in Python is available in the associated Github repository (https://github.com/bgoodlab/evolution_of_evolvability).

### 8.2 Theoretical predictions

The theoretical predictions in Figs. 2-4 were generated using the following procedures.

***Figure 2***. To generate the theoretical predictions in Fig. 2, we first solved for *x*_*cm*_ numerically as a function of 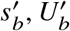, and *v*. We found that numerical accuracy was improved by using a modified version of Eqs. (S56) and (S118),

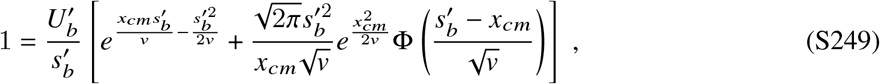

which has the same asymptotic behavior, but smoothly captures the transition between the multiple-mutations 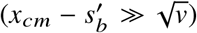 and quasi-sweeps regimes 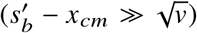. Numerical solutions to Eq. (S249) were obtained using the fsolve function from the SciPy library (107), with the measured value of *v* obtained from our forward-time simulations. We used the same procedure to solve for the wildtype interference threshold *x*_*c*_ using the analogous version of Eq. (S249). If 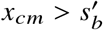, we estimated the fixation probability of the modifier using the multiple-mutations expression in Eq. (S84), while the quasi-sweeps prediction in Eq. (S135) was used for 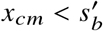. We used this procedure to generate all of the theory lines in panels A-C.

The boundary for the gray region in Fig. 2D was obtained by identifying the values of 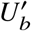 and 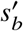 that minimize the true fixation probability 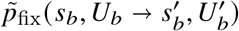 at a fixed value of 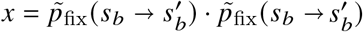. Using the leading-order approximations in Eqs. (S88), (S89), and (S90), the fixation probability of the compound modifier can be approximated as

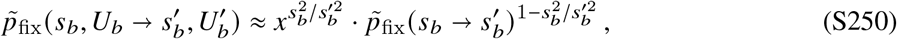

which eliminates the explicit dependence on 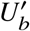. Minimizing this function with respect to 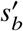 yields an analytical approximation for the boundary,

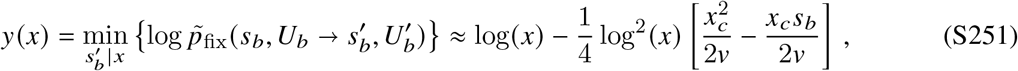

which was used to define the gray region in Fig. 2D.

***Figure 3***. The theoretical predictions for Fig. 3 were obtained using a similar procedure as in Fig. 2. For a modifier with a direct cost (*s*_*m*_ *<* 0; Fig. 3A,C), we used the multiple-mutations expression in Eq. (S99) if 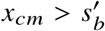, and the quasi-sweeps expression in Eq. (S139) if 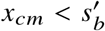; the integrals in Eq. (S139) were computed numerically using the quad function from the SciPy library (107). For a modifier with a direct benefit (*s*_*m*_ *>* 0; Fig. 3B,C), we used the corresponding predictions in Eq. (S97) when *x*_*cm*_ *< x*^∗^, or the dead-end predictions in Eq. (S115) otherwise, with *x*^∗^ defined by Eq. (S111).

The transition line in Fig. 3C was obtained by solving for the critical value of *s*_*m*_ that satisfies

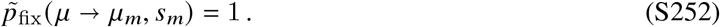

The theoretical line was obtained by numerically solving Eq. (S252) using the fixation probability predictions described above. The simulated values were calculated by linearly interpolating the observed fixation probabilities using the polyfit package in the Numpy library.

***Figure 4***. Predictions for the continuous distributions of fitness effects in Fig. 4 were obtained using the single-s mapping described in SI Section 5. We first calculated the value of *x*_*c*_ by numerically solving the wildtype version of Eq. (S249), with the effective parameters *s*_*b*_ and *U*_*b*_ defined by Eqs. (S151) and (S154). Using these estimates, we next checked whether the modifier was in the perturbative regime by numerically solving for *δx*_*c*_ using Eq. (S168). If *δx*_*c*_ *< v*/*s*_*b*_, we used the corresponding predictions from the perturbative regime in Eq. (S174), with 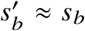_*b*_. If *δx*_*c*_ *> v s*_*b*_, we turned to the corresponding predictions from the modifier-dominated regime in SI Sections 5.3 and 5.4.

To do so, we numerically solved for *x*_*cm*_ using Eq. (S249) with 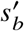 and 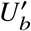 defined by Eqs. (S176) and (S179). For the bimodal distributions in Fig. 4, the solutions to Eqs. (S176) and (S179) can be expressed as a piecewise function,

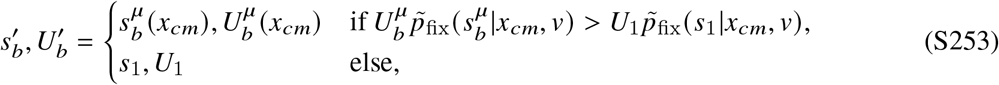

where 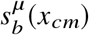 and 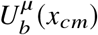 refer to the effective parameters of the wildtype distribution, but evaluated at *x*_*cm*_ rather than *x*_*c*_. The fixation probability of the modifier was then estimated as

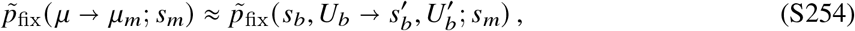

where the right hand side was calculated using the methods described for Figs. 2 and 3 above.

To capture the transition region between the perturbative- and modifier-dominated regimes, we continued to use our perturbative approximation if the modifier-dominated estimate of |*x*_*cm*_− *x*_*c*_| was less than the perturbative calculation of |*δx*_*c*_|. This convention ensures that our theoretical predictions are continuous at the border between the two regimes.

***Supplementary Figure S2***. The theoretical predictions for Fig. S2B,C were obtained using Eq. (S73). The theoretical predictions for Fig. S2D,E were obtained using the procedure used in Fig. 2 for a beneficial modifier without a short term cost.

***Supplementary Figure S3***. The theoretical predictions for Fig. S3A,B were obtained using the same procedure in Fig. 2 for a beneficial modifier with no direct cost or benefit.

***Supplementary Figure S4***. The forward-time simulations with diminishing returns epistasis in Fig. S4 followed a similar procedure to that outlined in SI Section 8.1. To enhance reproducibility, the simulations were allowed to “burn-in” for Δ*t* = 2 · 10^4^ generations before initiating diminishing returns. After this burn in period, diminishing returns was initiated and the mean fitness and typical selection coefficient, 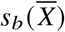, were recorded every 50 generations. The simulation results in Fig. S4A,B are the mean and average of 10 simulations.

After the typical selection coefficient in the population had declined to the chosen value 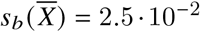, the fitness and abundance of each lineage was saved. Modifier lineages were then introduced at a constant rate for *t*_*U*_ = 1000 generations. This value was capped to ensure a successful modifier arose in a population with the chosen selection coefficient. To generate the comparisons of the neutral modifier and background selection coefficients in Fig. S4C, 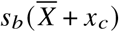 was obtained using an estimate of *x*_*c*_ from the model without diminishing returns. This estimate was obtained using a similar procedure to that in Fig. 2 with *v* numerically evaluated in a population with a constant selection coefficient, *s*_*b*_ = 2.5· 10^−2^. The results in Fig. S4C were then obtained using Eq. (S239).

After *t*_*U*_ = 1000, *U*_*m*_ was set to zero and the simulation continued until a modifier took over or all modifier lineages where purged. If a modifier did not take over, the population reverted back to the saved lineage distribution and this process was repeated *n* times until modifier took over. This allowed us to apply the same procedure in SI Section 8.1 to calculate the fixation probability of a modifier with *T* = *n* · *t*_*U*_. The theoretical predictions for the *θ* = *θ*_*m*_ line were obtained using the same procedure in Fig. 2 for a selection strength modifier with 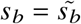 and 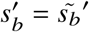^′^. The theoretical predictions for the *θ* = ∞ line were obtained using min{Eq. (S242), Eq. (S135)}, where *s*_*b*,0_ · *τ*_est,0_ was calculated from the relation 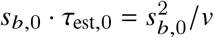. In this case, *v* was evaluated numerically in Wright Fisher simulations without diminishing returns for 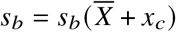.

### 8.3 Empirical example from Ref. (8)

The relative fitness estimates for the modifier example in Fig. 1A were obtained from Ref. (8). This study examined two strains of *E. coli* that were isolated from generation 500 of Lenski’s long-term evolution experiment (42). Relative fitnesses of the two strains at the first timepoint in Fig. 1A were obtained from head-to-head competitions under the same conditions as the original experiment. Ref. (8) reported these relative fitness values using the metric,

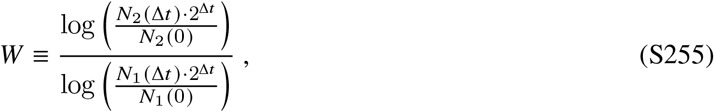

where *N*_*i*_ (0) is the number of colonies of strain *i* observed at the beginning of the competition, *N*_*i*_ (Δ*t*) is the (adjusted) number of colonies observed at the end of the competition, and Δ*t* is the length of the competition in generations. To enable more direct comparisons with our theory, we converted these *W* estimates to the relative (log) fitness,

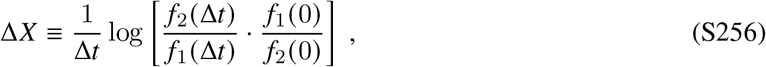

where *f*_*i*_ *(t*) is the relative frequency of strain *i* at generation *t*.

To perform this conversion, we assumed that the colony counts in Ref. (8) were adjusted so that a similar overall number of colonies were measured at both timepoints. This implies that *N*_*i*_ (Δ*t*) *N*_*i*_ (0) = *f*_*i*_ (Δ*t*) *f*_*i*_ (0). If we also assume that the competition experiments were started at a 1:1 ratio, so that *f*_1_ (0) ≈ *f*_2_ (0) ≈ 1/ 2, we can obtain a relation between the Δ*X* and *W* metrics,

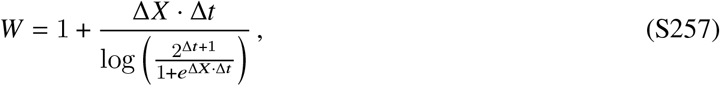

which depends on the duration Δ*t*. We used this expression to convert the average and 95% confidence intervals reported in Ref. (8) into the relative fitness values in Fig. 1A:

#### Initial timepoint

The fitness measurements for the initial timepoint were conducted over Δ*t* = 47 generations, and yielded a fitness disadvantage of *W* =0.937 (0.934,0.941) [mean and 95% confidence intervals from replicate competition assays]. Numerical conversion using Eq. (S257) yielded a log fitness difference of Δ*X*= -4.4% (-4.17%,-4.66%).

#### Second timepoint

The fitness measurements at the second timepoint were obtained after evolving each isolate for an additional 883 generations in 20 independent replay experiments. Pooled fitness measurements of the evolved populations were performed over Δ*t* = 6.6 generations, and yielded a fitness benefit of *W* =1.02 (1.003, 1.039). Numerical conversion using Eq. (S257) yielded a log fitness difference of Δ*X*=1.4% (0.2%, 2.6%).

